# A quantitative landscape of cell fate transitions identifies principles of cellular decision-making

**DOI:** 10.1101/2021.03.11.434982

**Authors:** M. Sáez, R. Blassberg, E. Camacho-Aguilar, E. D. Siggia, D. Rand, J. Briscoe

## Abstract

Fate decisions in developing tissues involve cells transitioning between a set of discrete cell states, each defined by a distinct gene expression profile. Geometric models, often referred to as Waddington landscapes, in which developmental paths are given by the gradient and cell states by the minima of the model, are an appealing way to describe differentiation dynamics and developmental decisions. To construct and validate accurate dynamical landscapes, quantitative methods based on experimental data are necessary. To this end we took advantage of the differentiation of neural and mesodermal cells from pluripotent mouse embryonic stem cells exposed to different combinations and durations of signalling factors. We developed a principled statistical approach using flow cytometry data to quantify differentiating cell states. Then, using a framework based on Catastrophe Theory and approximate Bayesian computation, we constructed the corresponding dynamical landscape. The result was a quantitative model that accurately predicted the proportions of neural and mesodermal cells differentiating in response to specific signalling regimes. Analysis of the geometry of the landscape revealed two distinct ways in which cells make a binary choice between one of two fates. We discuss the biological relevance of these mechanisms and suggest that they represent general archetypal designs for developmental decisions. Taken together, the approach we describe is broadly applicable for the quantitative analysis of differentiation dynamics and for determining the logic of developmental cell fate decisions.

## Introduction

Cell fate decisions in developing tissues involve gene regulatory networks comprising multiple genes, many molecular components and elaborate signalling dynamics. Despite the complexity, the outcome of cellular decisions is relatively simple: cells transition between a limited set of discrete cell fates, each defined by a distinct gene expression profile (Enver et al., 2009; MacArthur et al., 2009; Schiebinger et al., 2019). These transitions occur in a characteristic sequence regulated by extrinsic signalling. While quantitative models that describe signalling pathways and gene regulatory networks in great detail have been used to investigate cell differentiation and decision making, these suffer from a plethora of parameters and their behaviour is difficult to predict without case-by-case simulation. Hence, quantitative methods based on experimental data that represent and permit analysis of developmental processes at the scale of cell fate decisions would provide insight into the underlying principles and allow quantitative and testable predictions.

A popular and intuitive metaphor for the process of developmental decision making is the Waddington landscape, in which the differentiation trajectory of a cell is conceived as a ball rolling down a landscape of branching valleys, representing specific cell fates (Waddington, 1957). This can be mathematically formalised, with only minor changes, using dynamical systems theory (Camacho-Aguilar et al., 2021; Corson and Siggia, 2012, 2017; Huang, 2012; Mojtahedi et al., 2016). In this formulation the relevant dynamical systems are gradient-like and the system’s trajectories, which represent the developmental path of a cell, move downhill in this landscape. Thus, Waddington’s valleys correspond to the attractors of the system and sit at the minima of the landscape. Moreover, variation in the parameters of the dynamical system, caused by changes in the signals the cell receives, alter the landscape and give rise to bifurcations that destroy or create attractors. The transition to a new fate is signified by a cell entering a new basin of attraction, caused either by a signal-induced bifurcation or by a stochastic fluctuation resulting in a cell jumping from one attractor’s pull to another. In both cases the route from the old to the new cell state is defined by a saddle point in the landscape. This approach enables a rigorous link between the dynamical systems underlying gene regulatory networks and Waddington’s landscapes.

There are several advantages to a geometrical viewpoint. Firstly, since motion is always downhill, it gives a hierarchical structure to the dynamics and an intuitive understanding of the eventual fates. Secondly, a body of dynamical systems theory indicates that the qualitative structure of the landscape can be described by a relatively small corpus of universal normal forms (Zeeman, 1976). In particular, Rene Thom’s Catastrophe Theory provides a powerful classification scheme of the relevant bifurcations of such systems that is facilitated by the existence of the gradient-like structure (Smale, 1961; Thom, 1972). This theory suggests that although a system’s dimension might be large, the bifurcations can be described by low-dimensional systems. For example, in developmental systems it is common that differentiating cells transition from a progenitor state to one of two progeny fates. Such a decision can be an all-or-nothing one in which all cells make the same choice, or it can be one in which some cells make one choice and some the other one, enabling the allocation of a cell population to both fates. We introduce two 3-attractor landscapes, the binary choice (all-or-nothing) and the binary flip (allocation), that are the simplest archetypes underlying these two decision types.

To transform a developmental process from a metaphorical landscape description into a geometric model that allows quantitative and qualitative experimental predictions, quantitative data are needed together with methods to connect experimental measurements to a parameterised dynamical landscape. A framework based on Catastrophe theory and approximate Bayesian computation (ABC) has been developed and applied to experiments measuring the final outcome of a developmental process (Camacho-Aguilar et al., 2021). However, to test the full power of this approach and assess its general usefulness, quantitative measurements and perturbations during a differentiation process are necessary. To this end, we took advantage of the differentiation of neural and mesodermal cells from pluripotent progenitors using mouse embryonic stem cells (ESCs) exposed to different combinations and durations of signalling factors (Fig 1A; (Gouti et al., 2014; Tsakiridis et al., 2014; Wymeersch et al., 2021). ABC methods for parameter estimation based on matching summary statistics of temporally ordered data to those produced from simulations of candidate models naturally lend itself to our data and we use the proportion of cells in cell states at different time points as the summary statistic.

**Figure 1:**
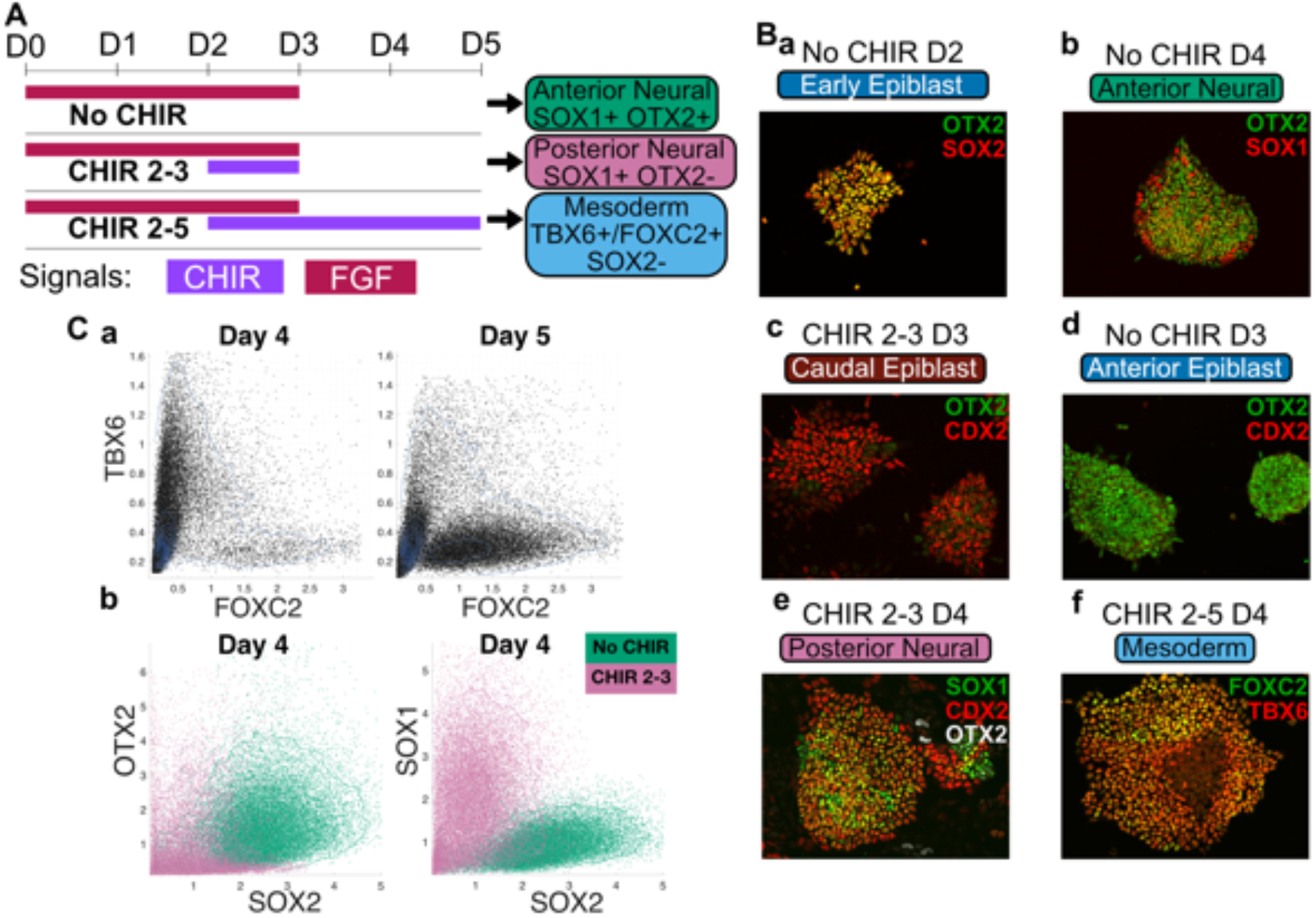
Single cell resolution analysis of the directed differentiation of ES cells to neural and mesodermal identities. A. Schematic of mESC differentiation. Embryonic stem cells differentiated in defined FGF and WNT signalling regimes adopt either Anterior Neural (AN), Posterior Neural (PN) or Paraxial Mesoderm (M) progenitor identities. Coloured bars show the times at which cells were exposed to CHIR and/or FGF. The expected dominant cells type at Day 5 and associated marker gene expression is given on the right. CHIR, CHIRON99021; D, Day. B. Representative immunofluorescence images of markers used to identify the different cell types. Progenitors differentiated for 2 days in FGF co-express the early epiblast markers SOX2 and OTX2 (Ba), and adopt SOX1+ OTX2+ AN identity by day 4 (D4) following withdrawal of FGF (Bb). Activation of WNT signalling at D2 with CHIR results in the upregulation of the posterior marker CDX2 (Bc) and the downregulation of OTX2 (Bd) by D3. Subsequent removal of CHIR at D3 leads to differentiation of SOX1+/CDX2+/OTX2-PN progenitors (Be), while sustained WNT signalling results in differentiation to TBX6+ early and FOXC2+ late paraxial mesoderm progenitors. C. Flow cytometry data. Flow cytometry analysis of individual progenitors shows that the progression from TBX6+ early paraxial mesoderm identity at D4 to FOXC2+ late paraxial mesoderm identity at D5 (Ca) under the sustained WNT signalling regime. At D4 OTX2 and SOX2 remain expressed at high levels in AN progenitors differentiated in the absence of CHIR, and are both reduced in PN progenitors induced by transient CHIR, whereas SOX1 expression is highest in PN progenitors (Cb).

Initiating the differentiation of ESCs, by withdrawing media supplements that sustain pluripotency, results in cells adopting a state that mimics the post-implantation epiblast. In the absence of further signals, these cells differentiate into neural progenitors with a molecular identity of the anterior nervous system (Ying et al., 2008). However, exposure to Wnt signalling at the epiblast stage blocks ESCs from adopting an anterior neural fate and instead cells acquire a caudal epiblast identity (Tsakiridis et al. 2014; Gouti et al. 2014a). This cell type, which in the embryo fuels axis elongation and the formation of trunk tissue, is responsible for generating the progenitors of the spinal cord and paraxial mesoderm (Wymeersch et al., 2021). Similarly, in vitro, caudal epiblast-like cells differentiate into spinal cord and mesoderm progenitors, with longer durations of Wnt signalling resulting in a higher proportion of mesodermal cells at the expense of neural cells (Blassberg et al., 2020; Gouti et al., 2017). Each of the cell states in this differentiation process is recognizable from well-defined gene expression profiles that can be assayed using representative marker genes. Hence the differentiation of ES cells into neural and mesodermal derivatives offers a well-characterised system in which to develop and test a dynamical landscape model of developing tissues and cellular decision making.

We used flow cytometry to measure the expression levels of representative marker genes in differentiating ESCs exposed to different signalling dynamics. These data provided an informative low-dimensional representation of cell states from which we developed a principled statistical approach to identify the attractors and the geometric form of the dynamical landscape. Using the ABC approach based on summary statistics we parameterised and refined the landscape and its changes with extensive training and validation datasets and tested it against specific experiments prompted by predictions from the model. The result was a quantitative model of the differentiation of neural and mesodermal tissue from pluripotent progenitors. Strikingly, the geometry of the landscape revealed two distinct decision-making mechanisms – a ‘binary choice’ as a one-or-other decision that commits cells to either anterior neural or caudal epiblast, and a ‘binary flip’ that simultaneously allocates cells to posterior neural and mesodermal fates. We discuss the biological relevance of these different mechanisms and suggest that they represent two dynamical archetypes that play a general role in differentiation dynamics. Taken together, the approach is broadly applicable for the quantitative analysis of cell fate dynamics and determining the design logic of developmental decisions.

## Results

### An in vitro system to quantify cell fate decisions

To develop a quantitative landscape model of cell fate decision making we took advantage of an in vitro system in which pluripotent mouse ESCs are directed to differentiate to distinct neural and mesodermal fates in response to defined signalling dynamics (Fig. 1A; (Gouti et al., 2014)). ESCs removed from pluripotency conditions and grown in basal media containing FGF2 (FGF) and an inhibitor of endogenous WNT secretion (LGK974), adopt a post-implantation epiblast-like (EPI) identity by Day (D)2 of culture (Fig. 1Ba; (Blassberg et al., 2020)), and subsequently, following removal of exogenous FGF, adopt an OTX2+/ SOX1+ molecular identity characteristic of anterior neural progenitors (AN) (Fig. 1Bb). By contrast, if WNT signalling is activated at D2 by the GSK3β antagonist CHIRON99021 (CHIR) concurrently with FGF signalling, EPI cells no longer differentiate to AN but instead, acquire a CDX2+ caudal epiblast (CE) identity at D3 (Fig. 1Bc). These differentiate to SOX1+ posterior neural progenitors (PN) in response to withdrawal of FGF and WNT signal activation (Fig. 1Be), while sustained WNT signalling drives the progressive differentiation of CE to TBX6+ early paraxial mesoderm (EM) and FOXC2+ late paraxial mesoderm (LM) identities (Fig. 1Bf and 1Ca).

To develop a quantitative description of the differentiation process we developed a flow-cytometry assay to measure simultaneously the expression of multiple marker proteins in individual cells. This enabled us to generate time-course data at single-cell resolution in a manner that was sufficiently scalable to explore the differentiation outcomes resulting from combinatorial modulation of FGF and WNT signalling dynamics. By inhibiting endogenous WNT secretion in all culture conditions with LGK974 we ensured that WNT signalling dynamics were entirely dependent upon the timing and concentration of CHIR addition. Moreover, as WNT signalling is known to induce FGF ligand production in CE cells (Amin et al., 2016) we tightly controlled FGF signalling dynamics through a combination of exogenous FGF withdrawal and inhibition of downstream FGF signalling with the small-molecule PD0325901 (PD). In total, we obtained 149 datasets in 4 experimental series, comprising 23 different signalling conditions assayed at 7 timepoints each (Supp Table T1).

We first defined a minimal set of markers sufficient to classify cells into distinct progenitor types. We employed the anterior marker OTX2 to distinguish between AN and PN progenitors, both of which express the neural progenitor markers SOX2 and SOX1. As expected (Gouti et al., 2014), OTX2 expression was permanently extinguished in SOX2+ SOX1+ PN progenitors following combined activation of FGF and WNT signalling between D2 and D3, whereas its expression remained high in SOX1+ AN progenitors (Fig. 1Cb). Moreover, the level of SOX1 was higher in PN cells, whereas SOX2 was higher in AN (Fig. 1Cb). We reasoned that SOX2 and SOX1 levels might be sufficient for classification of AN and PN progenitors in the absence of OTX2 measurement. This was subsequently confirmed experimentally (see below). In addition, we confirmed that exposure to WNT signalling induced the CE markers T/BRA and CDX2 in many cells at D3/D3.5 (Fig. S1C). Continued exposure to WNT signalling resulted in the loss of both T/BRA and SOX2 expression from differentiating CE progenitors at D4 and a high proportion of cells expressed TBX6+, characteristic of EM identity (Fig. S1C), which itself began to decrease at D5 as cells acquired FOXC2+ LM identity (Fig. 1Ca). As SOX2 expression is extinguished as cells commit to paraxial mesoderm identity ((Takemoto et al., 2011); Fig. S1B) we reasoned that either the presence of TBX6 and absence of SOX2 expression, or the absence of both TBX6 and SOX2 was sufficient to classify M progenitors.

### Clustering flow cytometry data identifies landscape attractors

In order to quantify the differentiation process, we hypothesized that each cell state represented an attractor of a dynamical landscape and we set out to develop a principled procedure to allocate cells to an identity. To this end, we used an algorithm based on fitting a Gaussian mixture model (GMM) to the multidimensional flow cytometry data (McLachlan et al., 2003) (fitgmdist in MATLAB). The GMM consists of a set of weighted multivariate normal distributions. We regarded each of these distributions as defining a cluster. A cell belonged to a cluster if its probability for the corresponding distribution was greater than a predefined threshold. Close to an attractor, the state variables (levels of gene expression in this case) are expected to have an approximately multivariate normal distribution since the dynamics near attractors are expected to be approximately linear (Kurtz, 1981).

To mitigate batch effects, we included a reference set consisting of the three signalling regimes comprising No CHIR, CHIR 2-3 and CHIR 2-5 (as described in Fig. 1A) in each experimental series (Initial, Test, and Prediction; see Supp. Table T1), and used the corresponding pooled data from these three conditions to fit and define the GMM probability distributions at each timepoint (Fig. 2A, SI Sect. 2). To choose the number of distributions in the GMM, m, we increased it sequentially from 1 until the scalar distributions for each cluster were clearly unimodal in each of the dimensions (Fig. 2B). Cells from all other samples within the experimental series were then classified by assigning them to the clusters defined by the GMM model using a threshold probability of p=0.65. Otherwise, cells were considered to be in transition between clusters (Unclassified Transitioning, UT). The process was insensitive to the choice of threshold probability p (SI Sect. 2).

**Figure 2:**
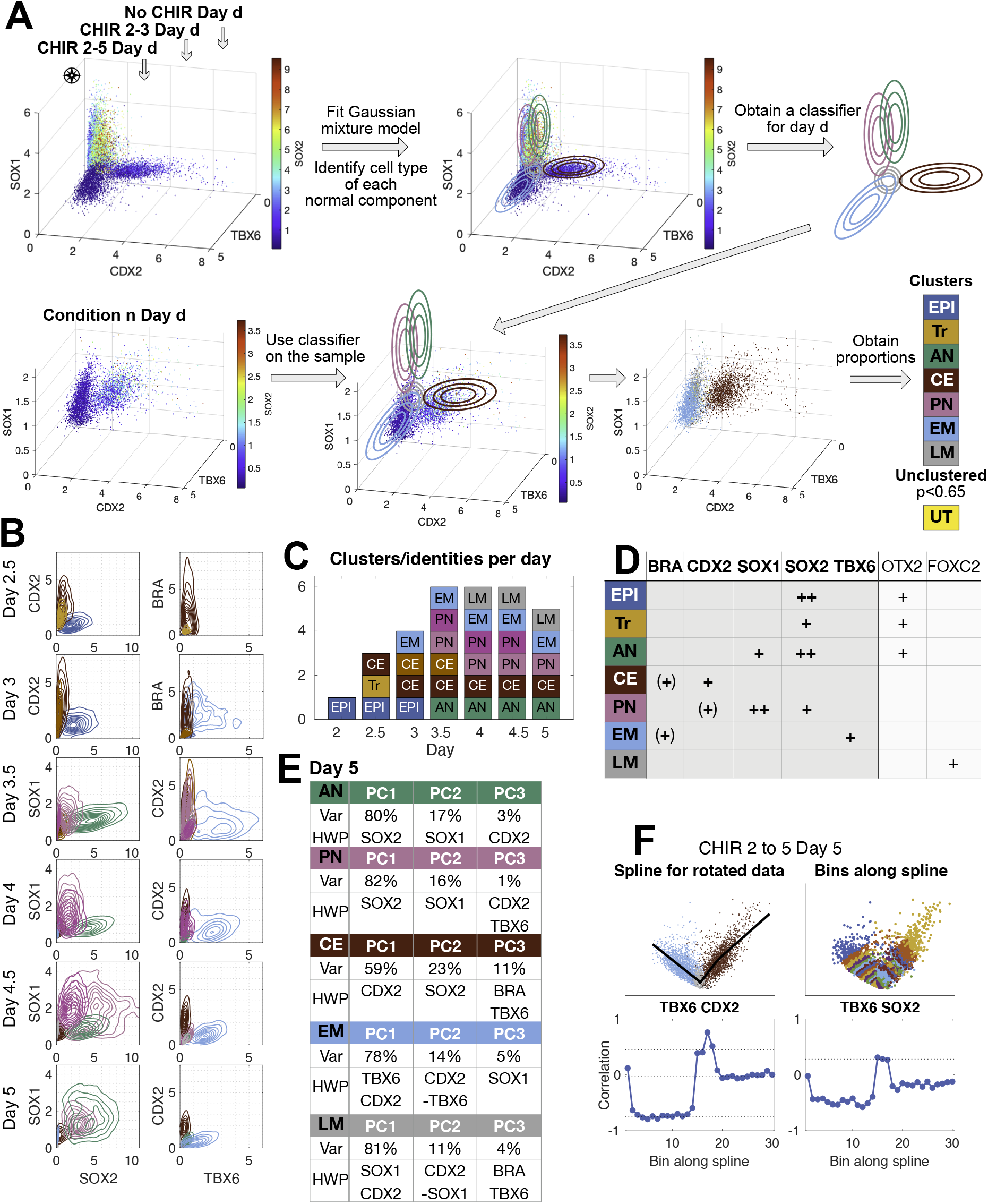
Clustering using Gaussian Mixture Models defines cell identities. A. Overview of the clustering method. Flow cytometry data from the three reference conditions (No CHIR, CHIR 2-3 and CHIR 2-5) for a specific timepoint were pooled and used to define a Gaussian Mixture Model such that each component corresponds to a cell identity. The model was used to cluster cells in samples from the same timepoint and experimental series. The number of cells assigned to each cluster was used to quantify the population proportions. Axes correspond to flow cytometry measurements (Units are in the order of ten thousand). B. Two dimensional joint distributions of flow cytometry data from the initial reference datasets illustrating the clusters identified in the pooled data. Colours correspond to the label assigned to the cluster as in (C). If the same label is assigned to several clusters different shades are used. Contours indicate cell density in each cluster. Axes correspond to flow cytometry measurements (Units are in the order of ten thousand). At day 5, from the total volume of the 5-dimensional cube that encompasses gene expression space, only 0.02% is occupied by the GMM that represents 90% of the cells. C. Number of cell types identified on each day of differentiation. Colours correspond to the assigned labels (D). If the same label is assigned to several clusters different shades are used. D. Table of cell type labels assigned to each cluster using marker protein expression: ++ denotes high levels of the markers, + denotes moderate levels of the markers, (+) denotes marker expression is optional, due to transitioning populations. The populations are Epiblast (EPI), Transition from EPI to posterior identity (Tr), Anterior Neural (AN), Caudal Epiblast (CE), Posterior Neural (PN), Early Paraxial Mesoderm (EM) and Late Paraxial Mesoderm (LM). The core set of markers used are indicated in bold. The extra information on additional markers is included in the right most columns. E. Principal component (PC) analysis of clusters at day 5 with variance explained by each of the first three PCs (Var) indicated. The highest weighted proteins in each PC (HWP) are shown. The PCs comprise mainly one or two proteins only. When more than one weight is relevant, we order them by magnitude. The relevant proteins in each cluster are different for different cells identities. F. Analysis of the local correlation structure in and between CE, LM and EM clusters at day 5. The data is rotated so that the first dimension extends across the two clusters. We show an illustrative projection of the rotated data on the two top panels. A smoothing spline is fit through the data (top-left). The data is binned along the spline into 30 bins (top-right) and gene-gene correlations in each bin computed (bottom). Correlations are maintained in bins within a cluster but change abruptly in bins around the cluster boundaries.

Once cells had been allocated to clusters, we analysed marker proteins and assigned cell identity accordingly (Fig. 2B and 2D). There was a good correspondence between the clusters found by the algorithm and the cell-type specific marker expression known to be present at each day of the three reference signalling regimes (Fig. 1A). The number of clusters needed to obtain unimodal distributions was larger for D3-D4.5, where there is more than one cluster corresponding to the same cell identity (Fig. 2C). For example, at D3 and D3.5 there are two clusters corresponding to CE cells: one defined by BRA+ and CDX2+ cells (so called neuromesodermal progenitors; (Wymeersch et al., 2021) and one with only CDX2+ cells (Fig. 2B-C and Fig. SI4).

The clustering algorithm identified the expected cell states in each of the datasets. Moreover, in line with our preliminary analysis, we were able to assign FOXC2+ late paraxial mesoderm identity on the basis of a SOX2-/TBX6-marker profile (Fig. S1B and S2C). Moreover, the clustering algorithm distinguished between OTX2+ AN and OTX2-PN progenitors based on their distinct levels of SOX1 and SOX2, as we had observed previously (Fig. S2B). However, as neither OTX2 nor SOX1/SOX2 expression levels were able to distinguish between anterior and posterior neural progenitors at D4.5 and D5 we extrapolated the proportions obtained at D4 to quantify their proportions in this case.

At D2.5 under FGF + CHIR signalling conditions, an additional population of cells with intermediate levels of SOX2 and low expression of BRA, CDX2, SOX1 and TBX6 was detected by the clustering (Fig. S2A). As these cells had begun to downregulate SOX2 this indicated that they were transitioning from EPI to CE, but had yet to acquire CDX2+ posterior identity (Blassberg et al., 2020). We considered them as a separate group and labelled them Transitioning (Tr). Thus, we were able to define clusters, corresponding to the attractors in the landscape, representing epiblast (EPI), anterior neural (AN), caudal epiblast (CE), posterior neural (PN) and mesoderm (M), and an additional transitioning population (Tr) using a minimal set of 5 marker genes (BRA, CDX2, SOX1, SOX2, TBX6) (Fig. 2D).

To test the validity of our cluster assignments, we examined the correlation of gene expression within a cluster. Principal component analysis of individual clusters identified distinct associations of genes with principal components for each cluster (Fig. 2E). Each principal component was associated with one or two genes. To test the hypothesis of multivariate normality we calculated local gene-gene correlations within each cluster and found smoothly varying patterns (Fig. 2F) in the correlation for each cluster. Correlations between gene expression reflect regulatory interactions between the components of the underlying gene regulatory network. Hence, consistency of the correlation structure across each cluster and differences in the structure in different clusters is evidence that the clusters correspond to distinct biological states. Moreover, the boundaries between adjacent clusters were characterised by abrupt changes in this correlation structure (Fig. 2F, SI Sect. 2). This is consistent with the idea that cell states represent high-dimensional attractors in gene expression space that are separated by saddle points.

### Two distinct binary decision landscapes

The data indicate that, in the absence of CHIR, EPI cells transition to AN after removal of FGF on D3, suggesting that FGF withdrawal causes a loss of the EPI attractor in such a way that cells escape towards the AN attractor (Fig. 3Ca No CHIR). On the other hand, if CHIR is added on D2 a substantial proportion of cells transition to CE by D3 (Fig. 3Ca CHIR 2-3). This suggests that, on addition of CHIR, the disappearance of the EPI attractor allows cells to escape towards the CE attractor. The distinct fate outcomes associated with these decisions suggests that each is caused by the bifurcation of the EPI attractor but with escape routes in different directions. In both cases, all cells escape in the same direction towards the same attractor.

**Figure 3:**
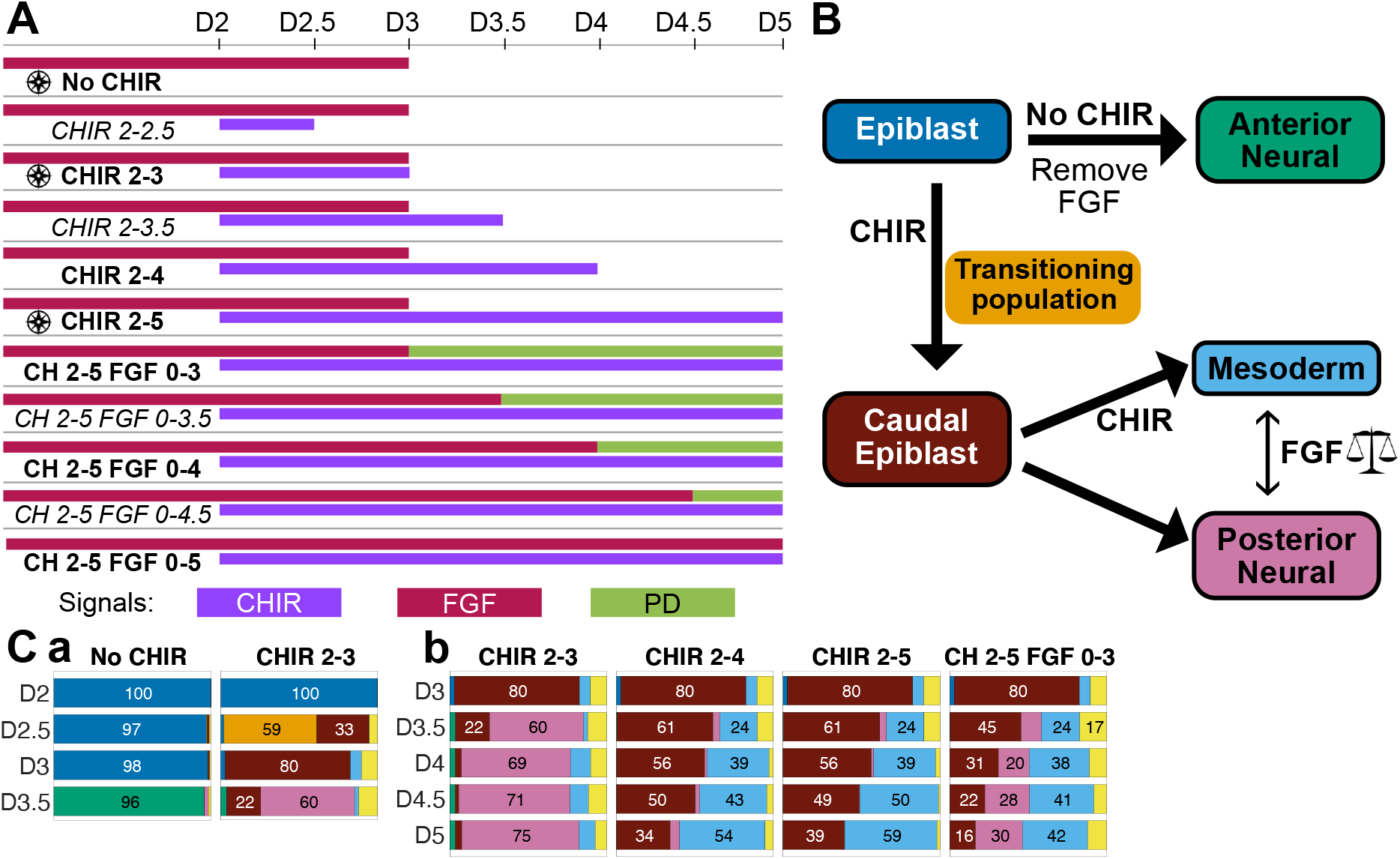
Initial experimental series. A. Schematic of the 11 experimental conditions used to acquire the initial dataset. Experimental conditions marked in bold comprise the training sets; the datasets for the conditions in italics formed the validation dataset and were not used for model fitting. The conditions marked with a symbol form the reference set. Coloured bars show the times at which cells were exposed to each of three signals. CHIR, CHIRON99021 (purple); FGF (red); PD, PD0325901 (green). B. Summary of the different cell types and transitions identified in the system. These define five attractors (with contour line) in the landscape together with a well-defined intermediate transition from Epiblast to Caudal Epiblast (no contour line). Thick arrows indicate the differentiation routes between cell types and the signals that drive them. C. The subset of fate proportions from the initial experimental series data that inform the geometry of the landscape: (a) informs the first decision from EPI to AN and CE; (b) informs the second decision from CE to PN and M. Colours correspond to cell identities (B). Cells with a low probability of belonging to any cell type were considered transitioning cells (UT) and labelled in light yellow.

There are three possible generic 3-attractor landscapes (SI Sect. 3) and others are combinations of them. These can each be modelled by simple two-dimensional dynamical systems (Rand et al., 2021). Only one of these is compatible with the above observations. This is related to Thom’s butterfly catastrophe (Thom, 1972): the three attractors, AN, EPI and CE, are separated by two saddle points (Fig. 4A) with one of them in the middle, EPI. We call this the *binary choice landscape*. Changing levels of signal cause bifurcations: cells in the central attractor EPI can transition to either CE or AN (Fig. 4Aa and Ac) but those in states CE and AN are only able to transition to EPI (Fig. 4Ab). This models an one-or-other/all-or-nothing decision where a population of cells in EPI chooses between the fates CE or AN. If cells commence the transition from EPI to CE and signals change pushing them towards AN, they need to return to EPI first before transitioning to AN.

**Figure 4:**
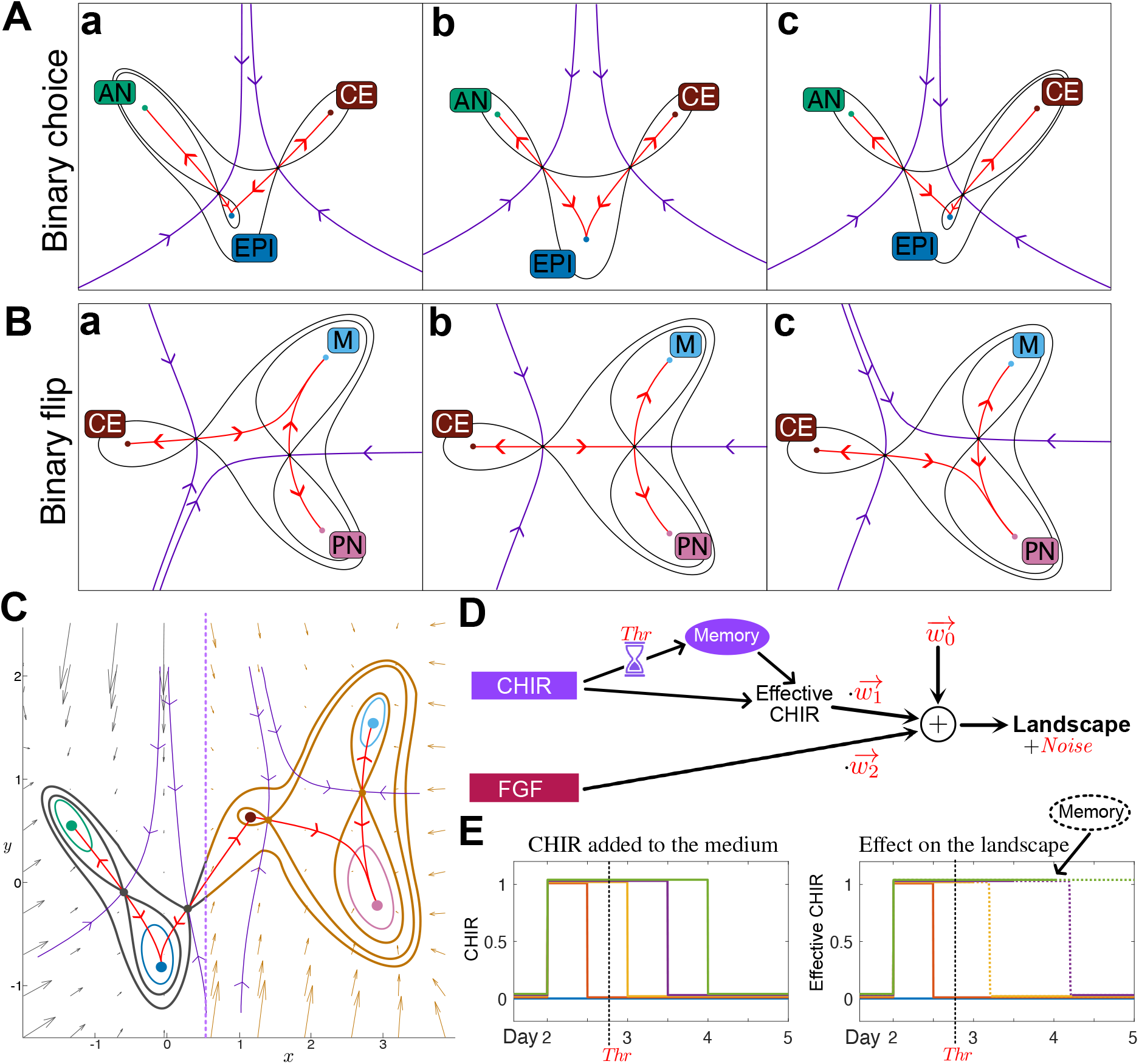
Two archetypal landscapes for binary cell fate decisions. A and B: Contour maps for three parameter sets (a,b and c) of two distinct three attractor potentials. The black lines denote contour curves passing through saddles, coloured dots are attractors, purple curves indicate the separatrices of the basins of attraction (stable manifolds of the saddles with the downhill direction indicated) and red curves are skeleton trajectories (unstable manifolds of the saddles with the downhill direction indicated). A. Binary choice landscape. Attractor EPI remains the central attractor for all three parameter sets (A abc). The curve defined by the unstable manifolds can be both smooth or cusp-like, as in the figure. Changes in parameters can result in a bifurcation either between attractor EPI and the saddle point separating AN and EPI (Aa) or between attractor EPI and the saddle point separating EPI and CE (Ac). These bifurcations leave the remaining saddle point intact. Hence this landscape models an all-or-nothing decision. B. Binary flip landscape. In this landscape a saddle point separates attractor CE from a basin that contains both attractors M and PN. The escape route (unstable manifold) emanating from the saddle associated with attractor CE can lead to either attractor M or PN. Varying parameters results in a swap (flip) in the attractor favoured by the escape route (Ba and Bc). This landscape models a decision in which cells differentiating from state CE can be flexibly distributed to either M or PN. C. The model was constructed from two binary decision landscapes smoothly connected through the common CE attractor. The first landscape is indicated in grey, the second in ochre. The two landscapes are different in character and allow different types of transitions. The model produces a gradient dynamical system with each part of the landscape defined by a parametrised height function that, with the addition of noise, determines the flow of the cells in the landscape. The two parts of the model each depend on 3 parameters (1 for velocity of cells and 2 for the shape of the landscape) for a total of 6 parameters. These parameters are functions of the signals in the medium as indicated in (D). D. Schematic of the effect of signals on model parameters. The effect of the signal concentration on the landscape is linear. A memory effect was incorporated for the effect of CHIR. Exposure to CHIR for longer than a Threshold (Thr) activates a memory term that triggers the persistence of the CHIR effect after its removal for a time proportional to the period of CHIR exposure (Memory = CHIRTime-Thr). The w’s are the weights of these effects and are 6 dimensional (w_0_ determines the landscape if there is no CHIR and PD). Noise amplitude is also a model parameter, in total the model comprises 20 parameters, these were estimated using ABC SMC with the training data. E. Example of the memory effect for different CHIR durations. If the duration of CHIR exposure is greater than the threshold (18h), the landscape will take CHIRTime-Thr to change after removal of CHIR. Thus, the total time CHIR affects the landscape is 2·CHIRTime-Thr.

Following the transition to CE, the data suggest a second decision associated with cells leaving the CE attractor adopting a mixture of M and PN fates. The ratio M:PN cells increases with the length of CHIR exposure and is not an all-or-nothing response as in the previous decision (Fig. 3Cb). Moreover, the way in which cells leave the CE state appears markedly different from the way that they leave the EPI state (Fig. 3Cb CHIR 2-5). Cells appear to leave the CE state at an approximately constant rate in CHIR induction conditions. This raises the possibility that transitions are caused not by bifurcation but by fluctuation driven escape from the CE attractor basin.

These data are compatible with a different generic three-attractor landscape (Fig. 4B). We call this the *binary flip landscape* because it allows signals to flip the escape route of cells leaving the CE state, tipping them into either state M or PN (Fig. 4Ba and Bc). When combined with stochastic noise, this allows cells leaving CE to be distributed with different ratios between M and PN (Fig. 4Bb).

Finally, we connect the two landscapes using the common CE attractor (Fig. 4C). This is done using a transition function to allow the cell’s trajectory to transition smoothly from one landscape to the other (SI Sect. 3).

To construct models, we took advantage of the normal forms of these two landscapes provided by catastrophe and dynamical systems theory (Arnold, V.I., Afrajmovich, V.S., Ilyashenko, Y.S., Shilnikov, L.P., 1994; Zeeman, 1976) as these minimise the number of parameters and state variables needed to capture the changing geometries. In addition, we postulated a linear relationship between the signals and model parameters and later validated this using the data (Fig. 4D). All cells in a sample were assumed to be affected by the same landscape, determined by independent effects of each signal (CHIR, FGF). To fit the data, we found it necessary to add a memory effect to this relationship (Fig. 4E). The data showed that removing CHIR at D3 or D3.5 gave different PN:M balances, while the PN:M balance remained near invariant if CHIR was removed at D4 or kept until D5 (Fig. 3Cb CHIR 2-4 and CHIR 2-5). This indicated that the effect of CHIR persisted for some time after its removal. To account for this memory effect, we maintained a CHIR signal in the model for a period of time proportional to the time CHIR had been in the system longer than a time threshold which was estimated in the fitting (Video CHIR 2-3.5). For FGF, since FGF signalling has been shown to generate a positive feedback and induce expression of FGF ligands (Amin et al., 2016; Blassberg et al., 2020), we maintained a level of 90% FGF when exogenous FGF was withdrawn, unless the inhibitor PD0325901 was added, in which case FGF signalling was set to 0.

The resulting system depends on 20 parameters: 3×6 for the relationship from the 3 signal changes (no signalling (no CHIR + PD), addition of CHIR and addition of FGF: the parameters w in Fig. 4D) to the six landscape model parameters, a memory threshold and a noise level (SI Sect. 3). In order to fit these, we used a training dataset of 7 treatment regimes (see Fig. 3A and 6) measured at 7 time points that distinguish 7 populations: EPI, Tr, AN, CE, PN, M, UT (a total of 343 measurements of cell fate proportions). We kept separate, for validation, a dataset of 4 experimental conditions with a total of 196 measurements (see Fig. 3A and 7). We took advantage of the approximate Bayesian computation (ABC) framework (Camacho-Aguilar et al., 2021; Toni et al., 2009) to generate posterior distributions of the parameters (see SI Sect. 4).

### Fitted landscape captures cell fate decisions

By extensive sampling from the resulting posterior parameter distribution for each signal combination we obtained simulation time series for cells in the landscape using the flow defined by the stochastic dynamical system (Fig. S3B). We computed the proportion of cells around each attractor and compared these with the corresponding experimental proportions (see SI Sect. 5 for details). For each timepoint and signal combination this produced a distribution of cell states and resulted in a 2×3 table of landscapes (Fig. S3A and Fig. 8A) corresponding to the different signal combinations used in the training experiments: CHIR on/off and FGF on/off or inhibited by PD.

Even though the parameters come from a distribution, the distribution of cell states was relatively tight about their mean with an average coefficient of variation of 7%. The larger size of some coefficients of variation was due to misidentification of clusters for a small number of experimental conditions. To compare simulated and experimental proportions we therefore used the mean simulated proportion. Comparison with the training datasets showed excellent overall agreement (Fig. 5; see SI for a detailed analysis). The largest differences tended to be associated with situations with a significant number of unclassified cells in the experimental data (UT), which we reasoned were cells transitioning between states. Importantly, the simulations reproduced subtle features of the data such as the constant rate at which cells leave the CE state in response to CHIR.

**Figure 5:**
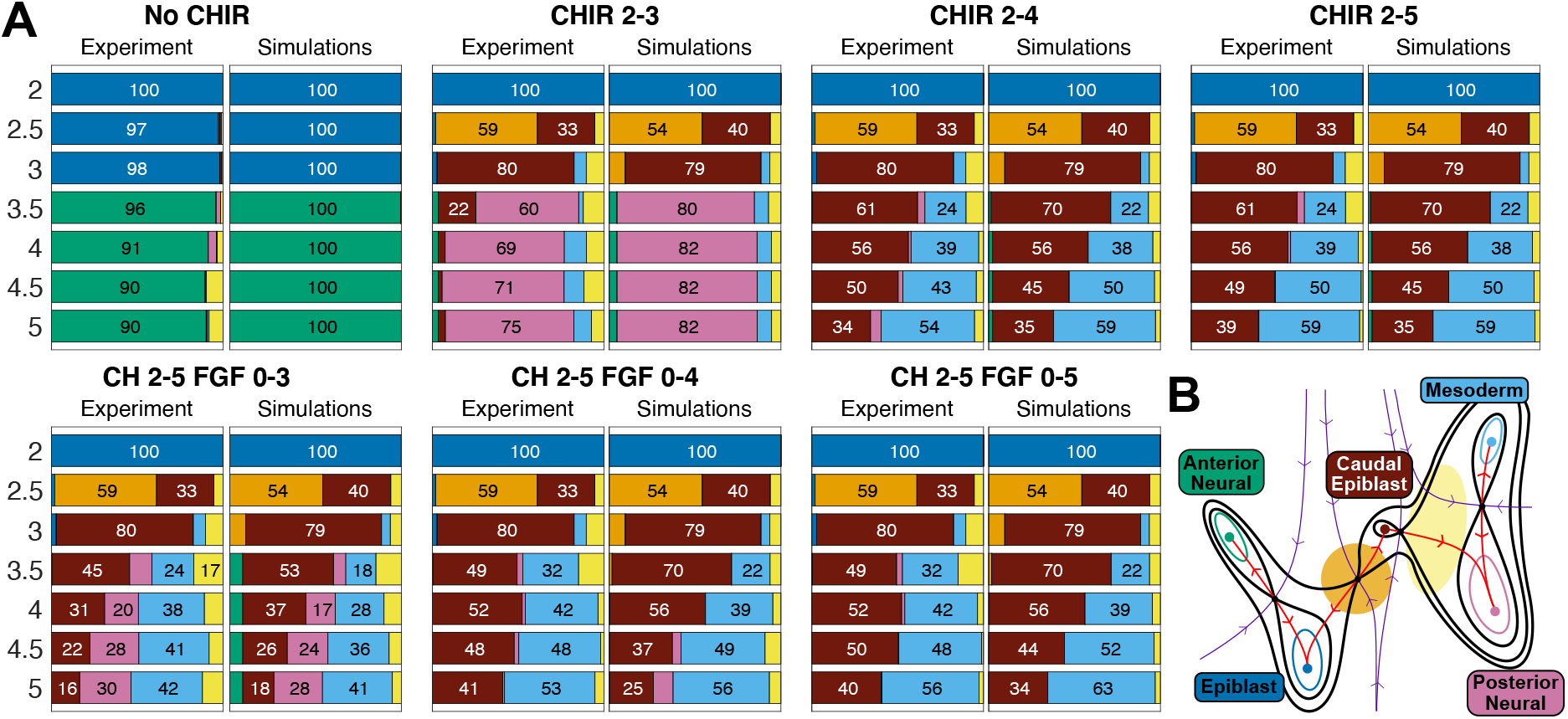
Comparison of model simulations to experimental data. A. For each condition at each time point the proportions of cells assigned to each cell identity by the clustering method (Experiment) were compared with the proportion predicted by the model (Simulations). For the simulated data the proportions of each cell type were obtained by averaging the proportions of cell types obtained by simulating the model using all 10000 parameter sets found by the fitting algorithm. Colours correspond to cell identities (B). Cells with a low probability of belonging to any cell type were considered transitioning cells (UT) and labelled in light yellow. B. Qualitative form of the global landscape model used in the fitting. Cell identities correspond to attractors in the landscape. Different signalling regimes change the particular form of the landscape.

We next examined simulations using signal combinations from the validation datasets computed with the model parameter obtained from the fitting to the training data. Again, there was good overall agreement (Fig 6). Strikingly, the simulations showed that the cells transitioning at D2.5 were “recaptured” by the EPI attractor after removal of CHIR (Video CHIR 2-2.5), as observed in the experimental data, where a proportion of cells transitioning to CE at D2.5 return to an epiblast state at D3. Since the data used to fit the model did not contain any conditions in which the EPI attractor was repopulated after cells had left it, these simulations provide a non-trivial validation of the geometry of the landscape and its parameterisation.

**Figure 6.**
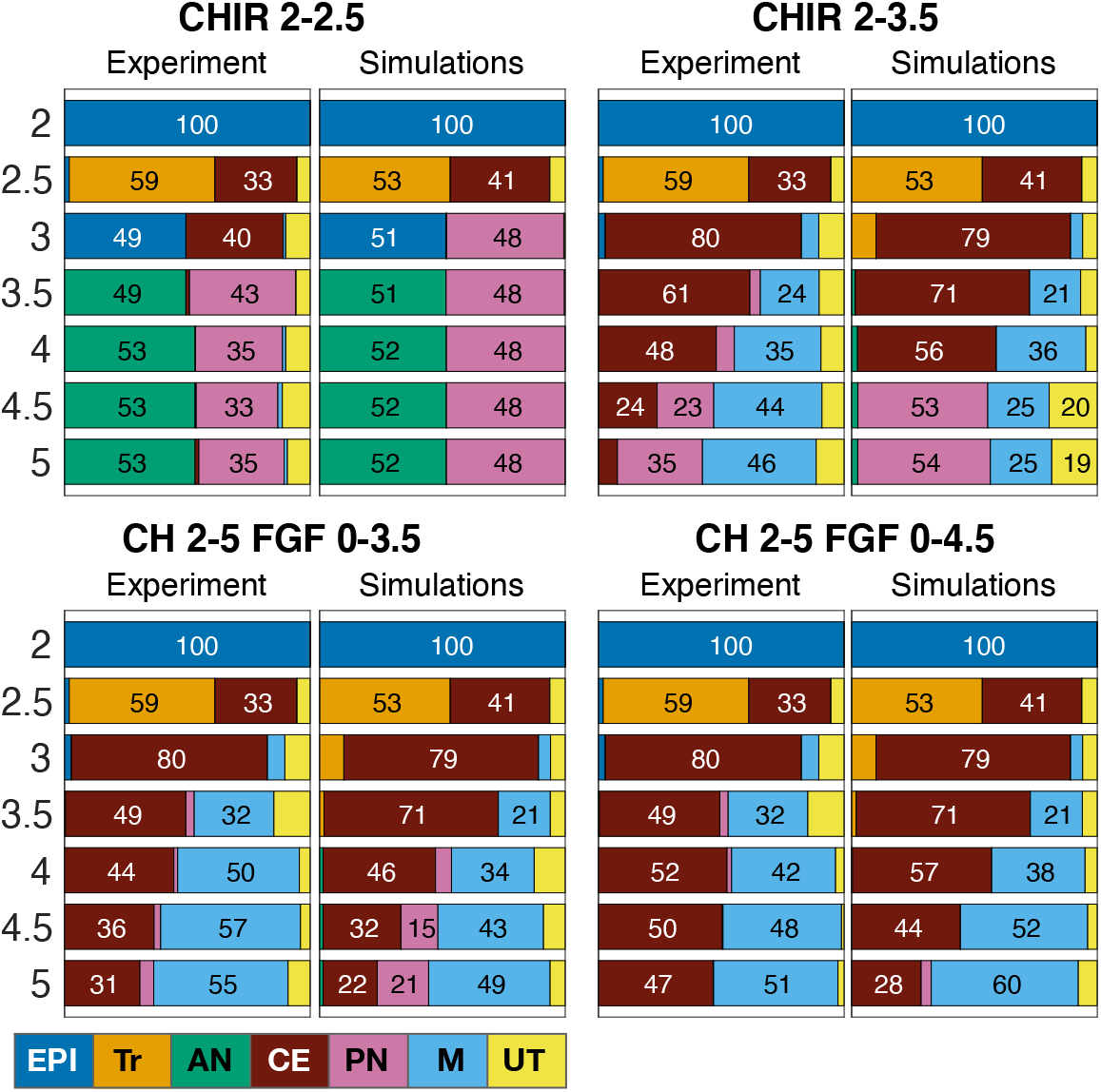
Comparison of model simulations to validation data. Experimental conditions not used in the fitting were compared with model simulations. For each validation condition at each time point the proportions of cells assigned to each cell identity by the clustering method (Experiment) were compared with the proportion predicted by the model (Simulations). For the simulated data the proportions of each cell type were obtained by averaging the proportions of cell types obtained by simulating the model using all the parameter sets found by the fitting algorithm. Colours correspond to cell identities as detailed. Overall, the model performed well at predicting the experimental results. We note that when CHIR is removed after 12h (CHIR 2-2.5) many cells that were in transition (Tr; orange) at D2.5 were recaptured by the EPI attractor both in the experiments and the simulations at D3. However, the simulations underestimated the CE the proportion of cells that remained CE at D3.

Closer inspection, however, revealed some differences between simulations and experimental data. Most notably, there was an underestimation of the CE population in some CHIR conditions (CHIR 2-2.5, CHIR 2-3, CHIR 2-3.5), together with an underestimation of M and overestimation of PN. Taking the results together, there was good agreement between the simulations and experimental data with the model accurately replicating the key decision processes in the differentiation pathway, this included the recapturing of cells by the EPI attractor after short CHIR exposure and the leakage of CE cell into M. Nevertheless, discrepancies in the exact proportions of cells between simulations and experiment prompted us to investigate further refinement to the model fit.

### Refined model accurately recapitulates experimental data

To test the model and determine the value of refining the fit, we took advantage of the model to design experiments. Simulations testing the effect of alternating periods of CHIR induction with periods of no CHIR suggested that the informative results were obtained when cells were induced into the transition state by exposure to CHIR for 12h at D2 and then subjected to either 1 or 2 pulses comprising 3h with no CHIR followed by 3h CHIR. After the pulses, cells were kept in continuous CHIR (Fig. 7A). Simulations of the single pulse predicted a mixture of anterior and posterior fates at D5 (Fig. 7B Initial Sim). By contrast, simulations of the double pulse predicted approximately 50% of the cells would adopt an AN identity and almost no cells would remain CE at D5 (Fig. 7B Initial Sim). That is, the model predicted that 3h of no CHIR and no FGF would be sufficient for cells to become AN and that two 3h pulses of no CHIR conditions would be sufficient for CE cells to become PN. Moreover, the model predicted that the transitioning population (Tr) would be present at day 3 under both pulsing conditions.

**Figure 7:**
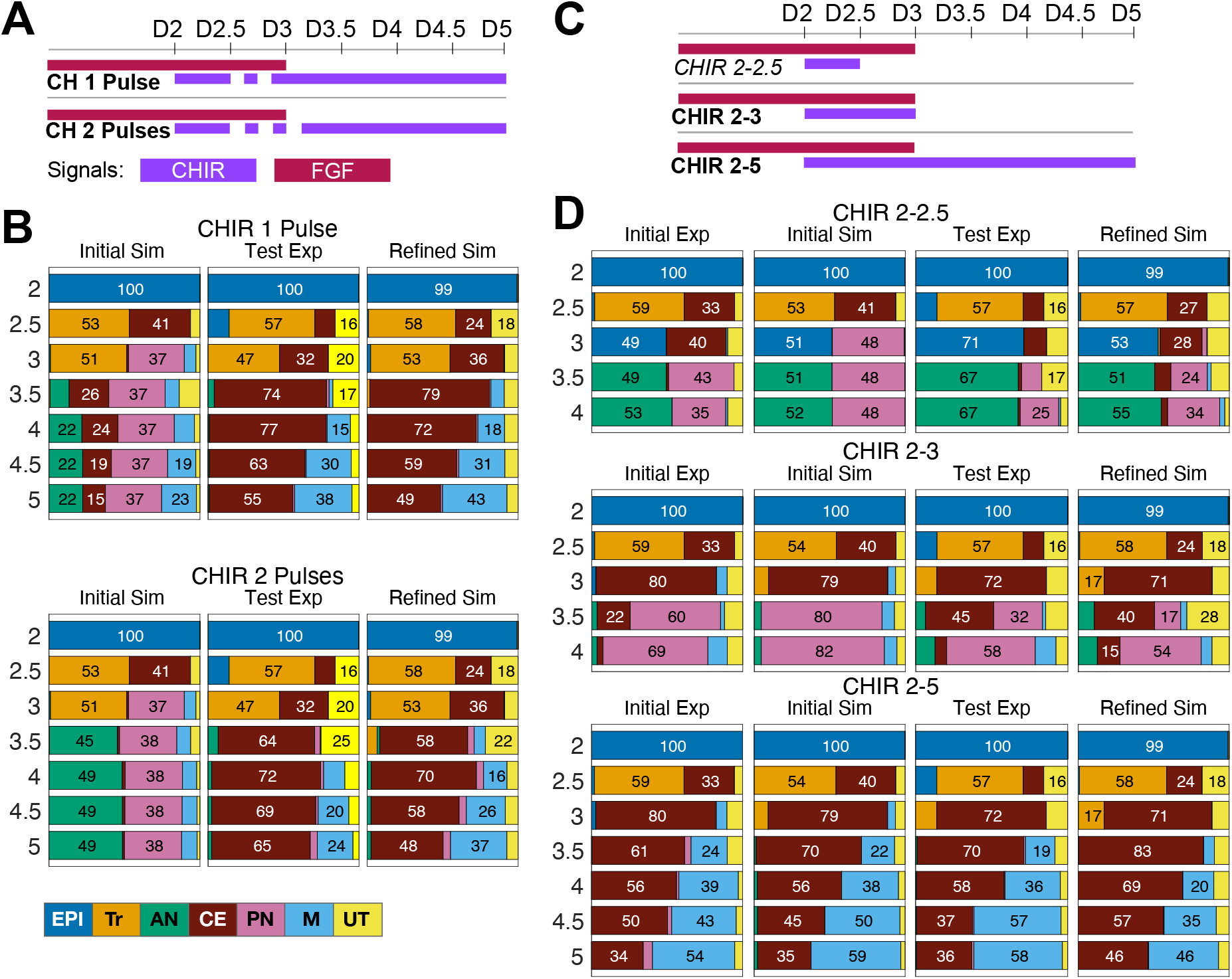
A refined model improves the accuracy of predictions. A. Details for the 2 experimental conditions designed to refine the model. Coloured bars show the times at which the CHIR (purple) or FGF (red) were added and removed from the medium. B. Comparison between the mean proportions of cell types predicted by simulations of the initial model (Initial Sim) and the refined model (Refined Sim), and the proportions of cell types obtained experimentally (Test Exp) for the conditions in (A). Colours correspond to cell identities as detailed. For the pulsing experiments, substantial differences were observed between the predictions of the initially fitted model and the experimental data. After using these experimental data together with the reference set to refine the model, the agreement between simulation results and experimental data was improved. C. Details for the 3 experimental conditions used to compare the accuracy between the initial model and the refined model. Coloured bars show the times at which the CHIR (purple) or FGF (red) were added and removed from the medium. D. Comparison of simulations from the initial model and refined model. The refined model performed better at predicting the outcome of short CHIR duration experiments (C).

We performed these two experiments (Fig. 7B Test Exp) as part of the Test experimental series. As expected, we observed the transitioning population was present at D3 in both conditions. At later days the experimental data differed from our predictions. First, few cells adopted an AN identity and second, a substantial CE population remained throughout D3.5 to D5. The continued presence of CE cells at D5 was consistent with the idea that CE represents an attractor in the dynamical landscape and it pointed to the initial fitting underestimating its stability. The discrepancy between simulations and experiments prompted us to refine the model fit.

To this end, we included the data from the pulsing conditions together with the reference set of conditions to retrain the model (see SI Sect. 6 for details). Adjusting these parameters markedly improved the performance of the model (Fig. 7B,D Refined Sim). The AN population under the two pulse conditions was almost non-existent and the CE population was increased, matching the experimental data. Data from CHIR 2-2.5 condition which was not used in the training was used as a validation and showed a good agreement with the experimental data. Comparison of the refined landscape (Fig. 8) with the initially fitted landscape (Supp. Fig. 3) revealed that the CE attractor did not bifurcate under No CHIR + FGF conditions and was very close to bifurcation under No CHIR + No FGF conditions allowing cells to remain in that area for longer. Moreover, the saddle point separating the AN attractor from the rest of the landscape had moved closer to the AN attractor, increasing the amount of time necessary for cells to commit to the AN fate under No CHIR + No FGF conditions.

**Figure 8:**
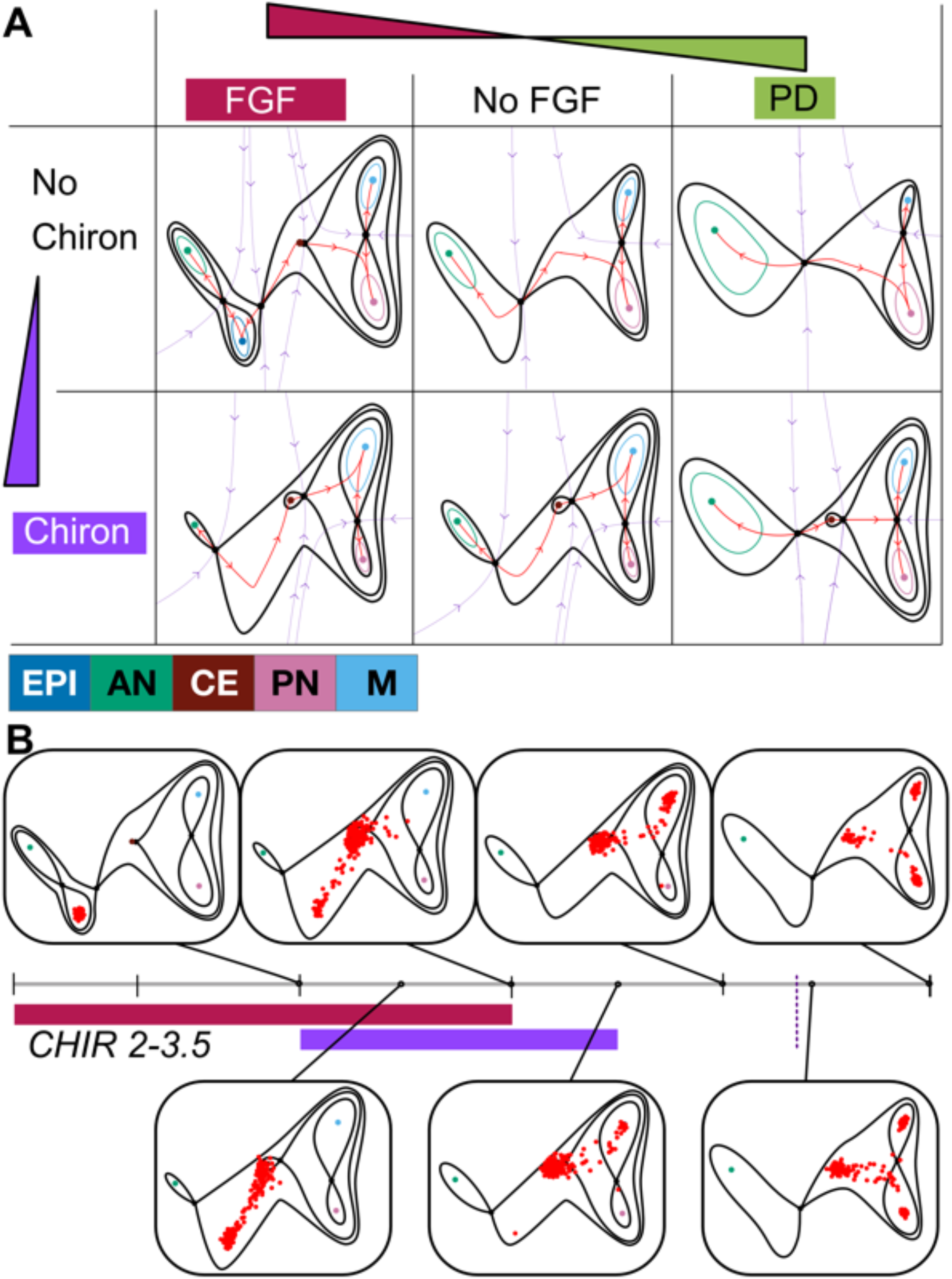
Landscape geometry and the effect of signalling. A. The landscapes produced by different combinations of signals are portrayed in the table. The changes in the landscape that result from different signal combinations arise by bifurcations of attractors and flips in dynamical trajectories (unstable manifolds) of the parameterised landscape family. Note that the landscape corresponding to No CHIR+PD (top right) is an extrapolation of the fitting and not based directly on data, it therefore represents an untested prediction of the model (see Fig. 10). Colours correspond to cell identities as detailed. B. Example of a simulation time series of the model for the signalling regime CHIR 2-3.5. Red points represent the location of cells in the landscape at the specified time points. Cells are initialised in the EPI attractor at D2. Their location evolves as given by the stochastic dynamical system defined by the landscape in Fig. 4C, which precise geometry is determined by the signalling regime. Three changes in the landscape are apparent. The addition of CHIR at D2 results in the bifurcation of the EPI attractor and cells follow the unstable manifold towards the location of the newly available CE attractor. At D2.5 many cells are still transitioning between EPI and CE. At later time points noise causes cells to abandon the CE attractor and follow the corresponding unstable manifold towards the M attractor. At D3 the removal of FGF has small effect in the landscape. At D3.5 CHIR is removed but the memory effect causes the landscape to remain the same. The termination of the CHIR memory effect (dotted purple line) results in the bifurcation of the CE attractor and the flip of the unstable manifold so that cells from the CE attractor transition into the PN attractor. To compare simulations to experimental data, simulated cells were clustered and compared with the experimental data at corresponding timepoints.

We conclude that in all conditions the CE attractor is close to its bifurcation point. As a result, when it is a genuine attractor its basin is shallow, hence it is easy for noise driven fluctuations to allow a cell’s trajectory to escape and proceed downhill to the PN or M attractors (Video CHIR 2-3.5). Thus, we see a gradual loss of CE cells at a roughly constant rate per unit time.

The fitted parameterised landscape provides insight into the capacity for cells to revert to a previous state: in developmental biology terminology the commitment of cells (Waddington, 1957). The model parameters determined by the fit mean that combinations of FGF and WNT signalling produce a constrained set of landscapes. This applies not just to the signal combinations used in the experiments but other reasonably conceivable combinations. This constraint means that certain transitions are not possible. For example, once cells have transitioned to the AN state this is irreversible as this attractor is deep and no bifurcation is allowed that destroys this state, thus cells are committed to AN and can no longer become CE in response to WNT signalling (Video CHIR 4-5)(Metzis et al., 2018). Similarly, the bifurcation of the EPI attractor in response to CHIR provides insight into the inability of CHIR + FGF inhibition with PD to revert epiblast cells to the ESC state (Video CHIR 2-5 FGF 0-3) (Guo et al., 2009; Ying et al., 2008).

Moreover, the model predicted a landscape for no CHIR with PD (Fig. 8A top-right), a condition not used in the experiments (Fig. 10). The table can also be used to predict the effect of different timings of signal changes (Fig. 7) and extrapolated to different concentrations of signals (Fig. 9).

**Figure 9:**
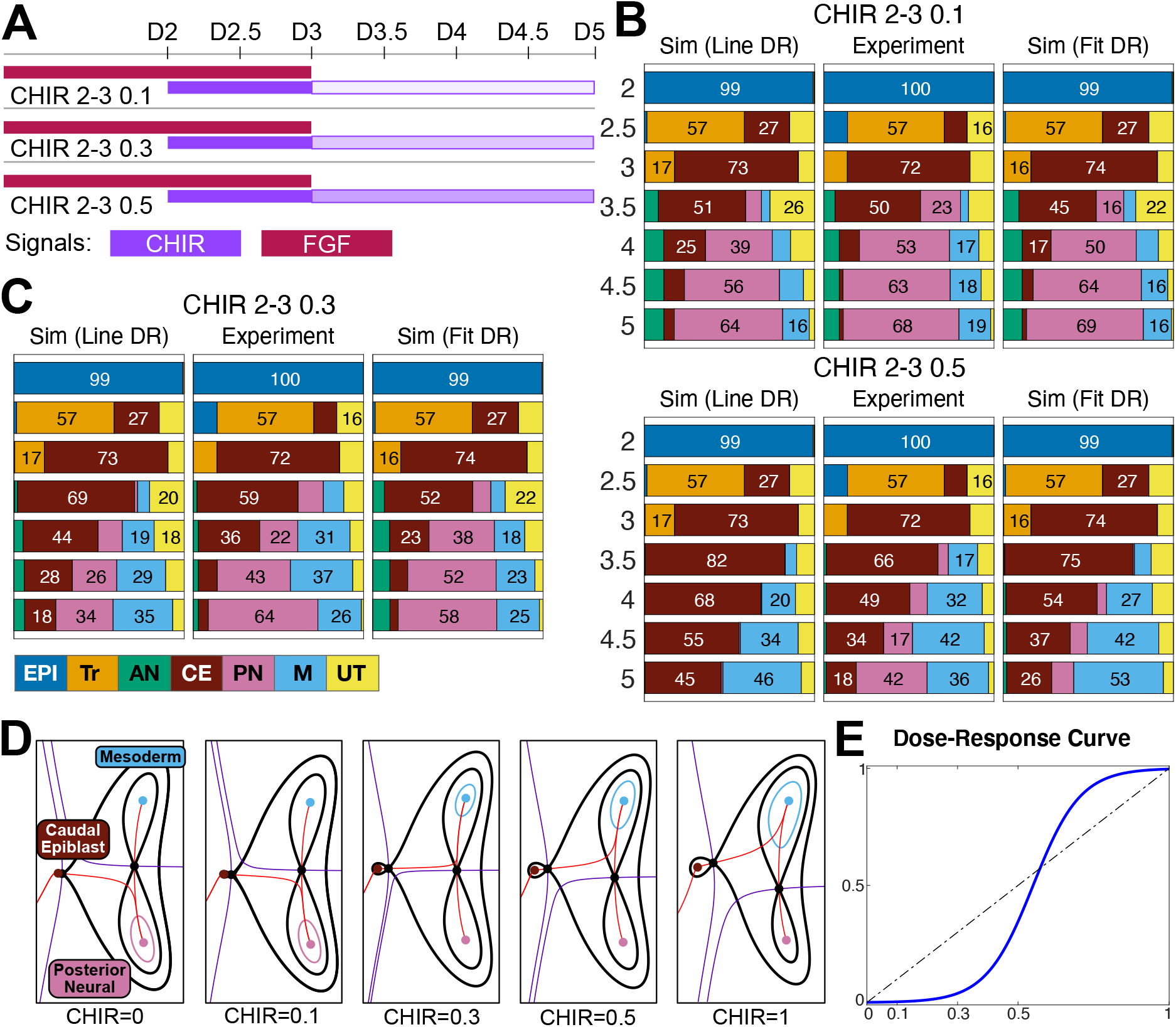
Incorporating the dose-response to WNT signalling. A. Three experimental conditions were designed to assess how the balance between PN and M fates are affected by CHIR concentration. Coloured bars indicate the times at which cells were exposed to CHIR (purple) and FGF (red). The intensity of shading indicates the concentration of CHIR, high (0.5mM), medium (0.3mM) or low (0.1mM) used between Day (D)3-D5. B and C. Comparison between the mean proportions of cell types predicted by simulations of the unrefined model (Sim (Line DR)) and the model refitted to the CHIR dose-response data (Sim (Fit DR), with the proportions of cell types obtained experimentally (Experiment). Colours correspond to cell identities as detailed. D. Landscapes for different CHIR concentrations. The stability of the CE attractor increases with CHIR concentration and the unstable manifold from the CE saddle flips from PN to M at ~0.3mM CHIR. E. Fitted sigmoidal dose-response curve compared with the linear effect used before refinement.

**Figure 10:**
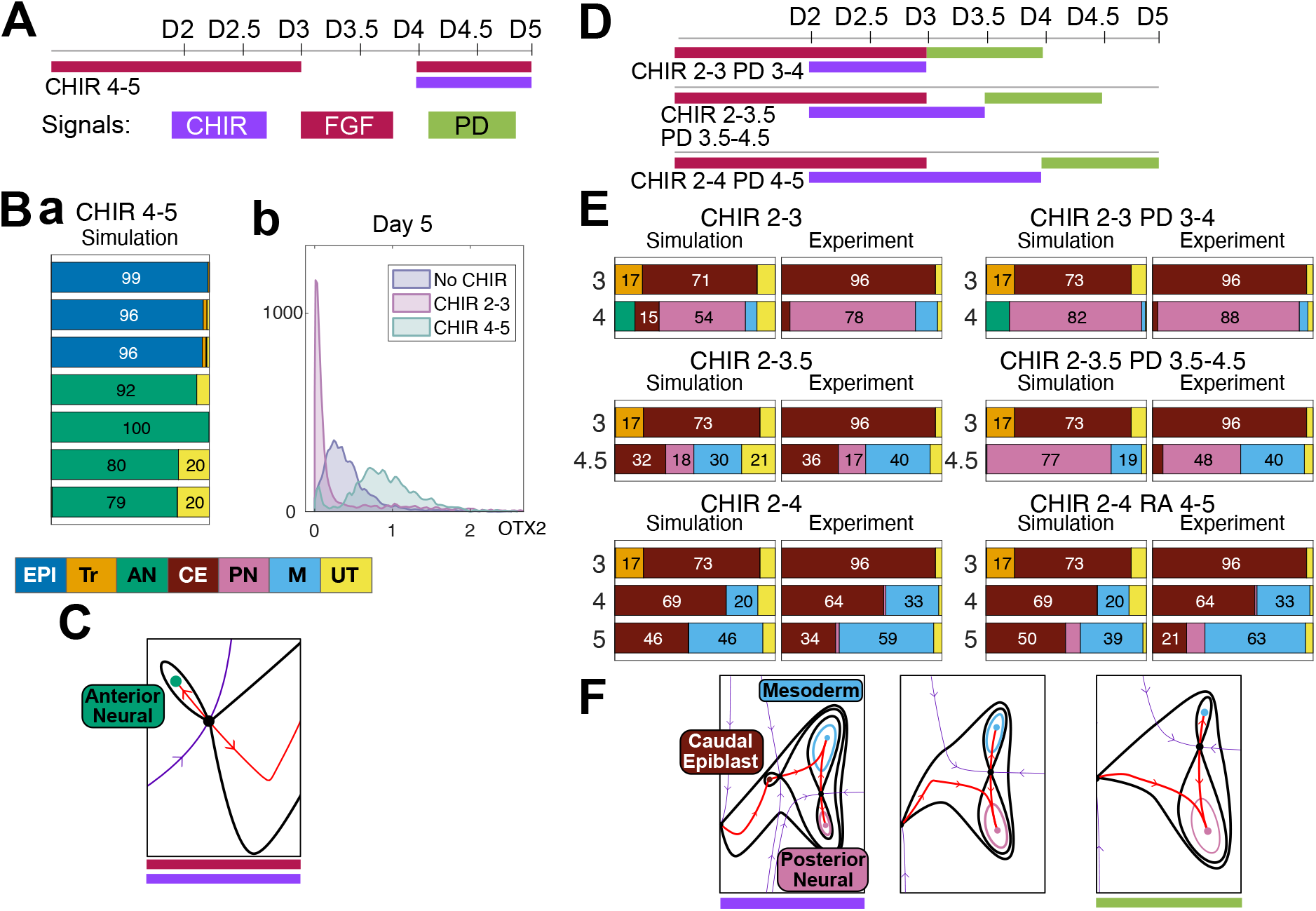
Quantitative predictions test the accuracy of the landscape. A. Details of the experimental condition used to assess the bifurcation of the AN attractor. WNT and FGF are introduced at later time points to challenge the commitment to AN. B. a. Mean proportions of cell types predicted by simulations for the condition indicated in (A). Colours correspond to cell identities as detailed. The model predicts that cells that have reached AN identity remain in this state if subsequently exposed to CHIR and FGF. b. Experimental data supports the prediction as AN cells retain high levels of OTX2 after CHIR induction ((Bb) and Fig. SI63). Colours correspond to cell identities (Fig. 3B). Cells with a low probability of belonging to any cell type were considered transitioning cells (UT) and labelled in yellow. C. Part of the landscape being tested in (A) and (B). D. Details of three experimental conditions used to assess the destabilisation of CE by the effect of PD with no CHIR. E. Comparison between the mean proportions of cell types predicted by the model (Simulation) and the experimentally observed proportions of cell types (Experiment). The data confirm the destabilisation of the CE attractor in PD conditions compared to control conditions and supports the memory effect of CHIR showing more CE cells for longer CHIR induction durations. Moreover, the effect of retinoic acid (RA; CHIR2-4 RA4-5) is also accurately recapitulated by simulating the inhibition of FGF signalling in the model. F. Part of the landscape being tested in (D) and (E).

### Identification of the dose-response curve for CHIR concentration

To investigate how the balance between M and PN was controlled in the second decision, we performed dose response experiments with CHIR after D3. We first simulated the model assuming a linear relationship between the effective CHIR signal level and CHIR concentrations (Fig. 9). Experimental results suggested a good correspondence for the low concentration (0.1mM), but larger differences were evident for higher concentrations (0.5mM) with a difference of 40% in PN, overestimating the CE and M proportions.

These results suggested that the model needed to be further refined to account for the dose-response effects of CHIR concentration. We tested whether the addition of a sigmoidal dose-response curve improved the model performance. For the fitting of the dose-response effect, we kept the distribution of all other parameters the same and fit the parameters of the sigmoidal curve. This substantially improved model performance. Although the half-response parameter did not produce a clear distribution, suggesting that additional concentrations would be necessary for a precise estimation of the dose-response curve, the estimation of cell fates in response to both 0.3mM and 0.5mM were close to the empirical data.

### Testing the geometry of the landscape

Having fit, refined and validated the landscape we set out to test its accuracy and investigate its implications by identifying predictions for which we had not previously analysed experimental data.

First, we noted that, in contrast to the EPI attractor that bifurcates with the saddle connected to the CE attractor in response to FGF + CHIR, the AN attractor was not destroyed by the addition of FGF + CHIR – a saddle remains separating AN and CE (Fig. 8A). This predicted that once cells had committed to AN identity, they would remain anterior and not adopt a CE identity after the addition of CHIR (Fig. 10Ba). We tested this hypothesis by adding FGF + CHIR at D4 to cells that had adopted an AN identity. In the standard AN condition (No CHIR + FGF D0-3), cells remained anterior in response to addition of FGF + CHIR at D4 as indicated by the high levels of expression of the marker OTX2 at D5 (Fig. 10Bb). These results were consistent with the constructed landscape, confirming that the saddle separating AN from CE does not bifurcate away in response to FGF + CHIR, and indicated that once EPI cells are committed, they remain anterior.

Second, because the landscapes define a family parametrised by two signals (CHIR and FGF) they include a hypothesised landscape for No CHIR + PD conditions, even though none of the datasets used to construct the model corresponded to this condition (Fig. 8A top-right). In this case the model predicts that, in the absence of WNT, the inhibition of FGF signalling results in a further destabilisation of the CE attractor and an expansion of the PN attractor (Fig. 10F). We tested this by removing CHIR and adding the FGF inhibitor, PD, to cells that had adopted a CE identity in response to different durations of CHIR exposure. The prediction was that cells in the CE attractor at the time of CHIR removal would transition to PN, while cells that had already adopted an M identity would remain in this state. Consistent with these predictions, the removal of CHIR and inhibition of FGF signalling after 24h (CHIR 2-3 + PD 3-4) resulted in a moderate increase in the PN population compared to control conditions. Maintaining the CHIR for 36h (CHIR 2-3.5 + PD 3.5-4.5) resulted in a marked increase in PN population at the expense of CE identity compared with control conditions (Fig. 10D), whereas the M population was less affected. This observation supports the idea that the CE represents a relatively shallow attractor that is maintained by CHIR signalling and destabilised by the removal or inhibition of FGF.

In the embryo, retinoic acid (RA) emanating from the somites, is proposed to inhibit FGF signalling in differentiating CE cells to promote neural differentiation (del Corral et al., 2003). We therefore compared the effect of RA addition to the effect of FGF inhibition. The addition of RA after 24h of CHIR (CHIR 2-3 + RA 3-4) or 36h (CHIR 2-3.5 + RA 3.5-4.5) resulted in almost identical proportions of cell types as obtained for the equivalent experiments with PD (Fig. SI64). Moreover, we also predicted that if cells were exposed to CHIR from D2-4, the memory effect of CHIR would result in the maintenance of a CE population at D5 even after the addition of PD or RA. Indeed, exposure of cells to RA after 48h of CHIR induction (CHIR 2-4 + RA 4-5) agreed with model predictions, with a large number of cells adopting M identity by D5. Importantly, in these experiments a substantial proportion of cells remains CE at 24h after removal of CHIR and addition of RA, consistent with the memory effect of CHIR hypothesised for the construction of the model.

Taken together, the results provide a quantitative model for the differentiation of neural and mesodermal cells from pluripotent progenitors that accurately predicts the effect of signals on the proportions of cell types generated. Moreover, the quantified landscape offers an intuitive visualisation of the differentiation process that highlights the distinctions between two types of decisions (binary choice and binary flip) that would be obscured in more complex gene-centric models.

## Discussion

It is increasingly easy to control cell fate and to generate quantitative data, raising the prospect of predictive engineering of organs and tissues (Kim et al., 2020; Shi et al., 2017). To support this endeavour, quantitative models that assimilate data and represent the successive steps in development are needed. Models are commonly constructed around genes and their interactions, typically defined with mass action kinetics, but their complexity and the large number of parameters involved can limit the scope of such models and obscure the underlying decision mechanisms. The Waddington landscape metaphor is an appealing simplification (Waddington, 1957), yet until recently it lacked both a mathematical foundation and meaningful engagement with data. We demonstrate an approach to construct, parameterise and analyse dynamical landscapes. We show that it makes specific and testable predictions as well as providing a visualisation of the developmental process that offers insight into the underlying mechanisms. This revealed two distinct strategies by which cells decide between one of two fates. We propose that they represent archetypes for developmental decisions that play a general role in developmental dynamics.

### Developmental transitions are represented by low-dimensional dynamics

Gene network models and real cellular systems develop by successive transitions among a small panel of fates, so the mathematics of dynamical systems and catastrophe theory can be applied (Arnold, V.I., Afrajmovich, V.S., Ilyashenko, Y.S., Shilnikov, L.P., 1994; Guckenheimer et al., 2013; Thom, 1969, 1972). Moreover, there is formal mathematical theory for such systems that reflects, extends and clarifies Waddington’s picture (Rand et al., 2021). This theory indicates that even when the state space (gene expression) is high dimensional, developmental transitions comprise the simplest saddles (those with 1-dimensional unstable manifolds) and their attractors. Consequently, the topology and behaviour of such models can be represented in two dimensions, so that it is conceptually meaningful to work in the low dimensional space and assert that relevant biology takes place on a two-dimensional manifold (Corson and Siggia, 2017). The final crucial step is to make the model quantitative by fitting a rich set of its summary statistics to those of the data. This links the powerful qualitative theories to quantitative models and provides an intuitive but rigorous means to analyse developmental dynamics.

We applied these ideas to the differentiation of embryonic stem cells in response to two signals, WNT and FGF, that together direct differentiation into precursors of neural and mesodermal tissues via intermediates that have received prior study (Blassberg et al., 2020; Gouti et al., 2014; Tsakiridis et al., 2014; Wymeersch et al., 2021). This system has the virtue that cell identity is controlled by external signals, hence can be modelled without the complexity of cell interactions. Landscape models describe the cells as they transition between valleys in the Waddington metaphor and the most informative data is obtained during transitions. We show that quantitative data using just five molecular markers and a carefully designed array of conditions and times was sufficient to define the cellular states and obtain informative summary statistics: the proportions of cells of a given type at each time point. This information was sufficient to identify the underlying geometrical model. The fixed points of the model were abstract and acquired meaning when gene expression patterns were related to in-vivo experiments. The morphogens control the basins of attraction of the fixed points and ultimately the saddle node bifurcations or flips that define cellular decisions. Strikingly, all the relevant cellular decisions could be captured in two dimensions. This strategy for constructing models of cellular differentiation contrasts with single-cell transcriptome analyses of tissue development that simultaneously profile 1000’s of genes (Schiebinger et al., 2019; Tanay and Regev, 2017) but often lack the resolution and diversity of conditions necessary to discern relevant cell fate transitions and the underlying structure of the developmental process.

### Quantitative descriptions of experimental embryology

The results support the notion that cell states correspond to attractors of a dynamical system. For the EPI, AN, PN and M states their assignment as attractors was unambiguous and the differentiation of cells between states involved bifurcations of attractors. For CE the results were more subtle and revealing. For some morphogen combinations the CE state appeared as an attractor, albeit a shallow one with a relatively small basin of attraction close to bifurcation, while for other morphogen combinations the CE state had bifurcated away. The consequence of the shallowness of the attractor is a gradual leakage of cells from the CE state and this, in contrast to the bifurcation of an attractor, explains the transitory nature of the CE state. Cells leave the CE attractor by jumping over a saddle into the basin of a different attractor. This observation not only illustrates the subtlety of the method, but also provides an explanation for the progressive elaboration of the anterior-posterior axis of vertebrate embryos (Henrique et al., 2015; Wymeersch et al., 2021). This process involves the production of trunk mesoderm and spinal cord over an extended period of developmental time from bipotential progenitors (CE) located at the tail end of the elongating body. Ensuring the CE state is poised close to bifurcation allows the constant differentiation of a fraction of cells to neural and mesodermal tissue while retaining an uncommitted progenitor population.

The model we constructed terminates in one of three fates: mesoderm, anterior neural, or posterior neural. Cells in these states are committed, they cannot leave these attractors and remerge on our fate plane, but they are not endpoints as cells continue to differentiate. We envision that longer differentiation trajectories could be constructed, branch by branch, with the modular and hierarchical property of the models facilitating the gradual elaboration of the full differentiation tree. By fitting and refining the saddles and attractors as they move in response to extrinsic signals, such models generate quantitative predictions and suggest complex temporal stimuli that might stabilize or disrupt populations of cells in the dynamical landscape. In this way, landscape models delineate phenomenological features of experimental embryology in quantitative form.

### Two archetypal decision mechanisms

The landscape that resulted from our analysis comprises two sub-landscapes. These are generic in the sense that they occur naturally in dynamical systems without any special conditions and when they occur, they are structurally stable and robust to misspecification or perturbation of the dynamical system. Each encodes a fundamentally different type of binary decision. For the AN-EPI-CE decision, the data support the notion that the EPI state is destabilised by collision of a saddle point between it and either the CE or AN attractor. Importantly, after either bifurcation, the saddle separating the EPI attractor from the alternative fate remains intact. The result is an all-or-nothing choice that commits cells to a fate. This agrees with previous experimental observations that epiblast cells commit to an anterior or posterior identity prior to acquiring a neural fate (Metzis et al., 2018). Since the overall effect of this landscape is that signalling results in all cells in the EPI state choosing and committing to either the CE or AN state, we call this landscape the *binary choice landscape*.

The decision for the differentiation of CE to either PN or M is different. The landscape indicates that the proportion of PN and M cells generated in response to different amounts of WNT signalling is a consequence of the location of the unstable manifold of the saddle that separates the CE attractor from PN and M. For prolonged and high CHIR conditions, the escape route connects to the M attractor, whereas in the absence of WNT signalling it flips to PN. For intermediate conditions stochasticity produces a mixed outcome with a progressive increase in the ratio of M to PN as WNT signalling increases. Because the flip in the unstable manifold is the essential dynamical phenomena controlling the allocation of cells, we call this the *binary flip landscape.* Previously, a pitchfork bifurcation had been postulated for this transition (Steventon and Martinez Arias, 2017). However, in contrast to our landscapes, this is not generic but only occurs with an additional constraint equivalent to symmetry between M and PN and it is reversible such that cell states are only maintained if signalling is maintained.

The results help clarify the idea of cell commitment (Barresi and Glibert, Scott F., 2020; Waddington, 1957), by explaining when cells that have escaped a basin of a bifurcated attractor can or cannot be recaptured by reversing the signal and reintroducing the attractor. For the EPI to CE transition, recapture is possible for a limited time after which they are committed to the transition. For the EPI to AN transition, once cells have entered the AN state they cannot escape because the AN attractor never bifurcates away. In this view, therefore, commitment becomes a dynamical property of the landscape.

Crucially, theoretical considerations (Rand et al., 2021) indicate that there are relatively few generic three-attractor landscapes. Thus, the binary choice and binary flip landscapes are likely to be encountered repeatedly in developmental decisions. Moreover, as we show here, these landscapes can be linked together to provide an ordered hierarchy that accounts for a differentiation pathway comprising multiple decisions. Together this leads us to suggest that the binary choice and binary flip landscapes represent design archetypes that underpin cellular decisions throughout embryonic development.

### Outlook

In our approach, we did not fit flow cytometry data directly to the dynamical variables, but instead relied on the assignment of cells to discrete fates. We sampled the population frequently in time to capture the transitions among the fates in the topological model and thus obtained quantitative predictions about what matters most, the discrete cell states. A future goal will be to develop tools that directly link gene expression measurements and detailed gene centred models to landscape models. The minimal spatial organisation in the differentiating stem cell colonies, meant only the response of individual cells to exogenous signals needed to be considered. To apply these models to cells of an intact embryo, we would surmise that the cell intrinsic landscape remains the same, but the spatial-temporal behaviour of the morphogens would need to be measured or inferred. To probe such models, it will be necessary to perturb signalling pathways with sufficient temporal resolution to catch cell fate transitions, experimental manipulations that do not control time are manifestly less useful. Thus one can hope to reconstruct development in space-time analogous to the reconstruction of differentiation from single cell expression data (e.g. (Karaiskos et al., 2017; Nitzan et al., 2019) but in a way that provides quantitative predictions and insight into the underlying mechanisms.

## Supporting information

Movie_CHIR_2-3p5

Movie_CHIR_2-2p5

Movie_CHIR_2-5_FGF_0-3

Movie_CHIR_4-5

## Acknowledgements

We thank Despina Stamataki for experimental support and Francis Corson, Joaquina Delas, Ashley Libby, Rory Maizels and Archishman Raju for critical feedback on the manuscript. We thank Paul Brown for technical support and Yashin Dicente Cid for critical discussions. We are grateful to the Crick Science Technology Platforms in particular to Flow Cytometry Facility and High-Performance Computing.

## Author Contributions

RB, JB, ECA and DAR conceived the project. RB established the in vitro differentiation system and flowcytometry workflow, performed all experiments, and pre-processed data. MS and ECA analysed data. MS designed and performed the clustering procedure with input from ECA and RB. All authors interpreted data and designed experiments. MS, DAR and EDS designed the mathematical model with contributions from ECA. ECA designed and implemented the software to simulate and fit the mathematical model with input from MS. MS performed the fitting with input from ECA. MS, DAR, & JB wrote the manuscript with contributions from RB and ECA. EDS reviewed and edited the manuscript.

## Materials and Methods

### Cell lines

The XY mouse ES HM1 TetON line (Serafimidis et al., 2008) was used for all experiments. ES cells were maintained at 37°C with 5% carbon dioxide (CO2) and routinely tested for mycoplasma.

### ES cell culture and differentiation

All mouse ESCs were propagated on mitotically inactivated mouse embryonic fibroblasts (feeders) in DMEM knockout medium supplemented with 1000U/ml LIF (Chemicon), 10% cell-culture validated fetal bovine serum, penicillin/streptomycin, 2mM L-glutamine (GIBCO). To obtain EpiLCs and CEpiLCs, ESCs were differentiated as previously described (Gouti et al., 2014) with the addition of the porcupine inhibitor LGK974 (Cayman) in all culture medium. Briefly, ESCs were dissociated with 0.05% trypsin, and plated on tissue-culture treated plates for two sequential 20-minute periods in ESC medium to separate them from their feeder layer cells which adhere to the plastic. To start the differentiation, cells remaining in the supernatant were pelleted by centrifugation, counted, and resuspended in N2B27 medium containing 10ng/ml bFGF (Peprotech) + 5μM LGK974, and 50,000 cells per 35mm gelatin-coated CELLBIND dish (Corning) were plated. N2B27 medium contained at 1:1 ratio of DMEM/F12:Neurobasal medium (GIBCO) supplemented with 1xN2 (GIBCO), 1xB27 (GIBCO), 2mM L-glutamine (GIBCO), 40mg/ml BSA (Sigma), penicillin/streptomycin and 0.1mM 2-mercaptoethanol. To generate epiblast (Epi) cells, the cells were grown for 72 hrs in N2B27 + 10ng/ml bFGF + 5μM LGK974. To generate caudal epiblast (CE), cells were cultured with N2B27 + 10ng/ml bFGF + 5μM LGK974 for 48 hrs, then N2B27 + 10ng/ml bFGF + 5μM LGK974 + 5mM CHIR99021 (Axon) (FLC-medium) for a further 24hrs. CE were differentiated to posterior neural progenitors (NP) by removal of bFGF and CHIR from culture medium at 72hrs, and to paraxial mesoderm (M) by removal of bFGF and maintenance of 5mM CHIR from 72hrs onwards. When investigating the role of endogenous FGF producton either the MEK inhibitor PD0325901 (500nM) was employed to inhibit downstream FGF signalling or retinoic acid (10nM) was added to inhibit expression of FGF. For all experiments described, cells were cultured for 48hrs before changing medium. Medium changes were then made every 12 hours.

### Immunofluorescence

Cells were washed in PBS and fixed in 4% paraformaldehyde in PBS for 15min at 4°C, followed by two washes in PBS and one wash in PBST (0.1% Triton X-100 diluted in PBS). Primary antibodies were applied overnight at 4°C diluted in filter-sterilized blocking solution (1% BSA diluted in PBST). Cells were washed 3x in PBST and incubated with secondary antibodies at room temperature, for 1hr. Cells were washed 3x in PBST, incubated with DAPI for 5 min in PBS and washed twice before mounting with Prolong Gold (Invitrogen). Cells were imaged on a Zeiss Imager.Z2 microscope using the ApoTome.2 structured illumination platform. Z stacks were acquired and represented as maximum intensity projections using ImageJ software. The same settings were applied to all images. Immunofluorescence was performed on a minimum of 2 biological replicates, from independent experiments. Primary antibodies used: SOX2 (Mouse, Santa Cruz sc-365823), OTX2 (Goat, R&D AF1979), CDX2 (Rabbit, Abcam ab76541), TBX6 (Goat, R&D AF4744), FOXC2 (Goat, R&D AF6989). Secondary antibodies raised in donkey coupled to AlexaFluor-488, 568, or 647 fluorophores (Molecular Probes) were used at 1:1000 dilution throughout. SOX1 was labelled with AlexaFluor-647 conjugated antibody (Mouse, BD 562224).

### Intracellular flow cytometry

Cells were washed in PBS and dissociated with minimal accutase (GIBCO). Once detached cells were collected into 1.5mL Eppendorf tubes by dissociating in N2B27 and pelleted. Cells were resuspended in PBS, pelleted and resuspended in 4% paraformaldehyde in PBS. Following 15min incubation at 4 C, cells were centrifuged, resuspended in PBS, and stored at 4°C for future analysis. On the day of flow cytometry, cells were counted and equal cell numbers were transferred for staining in v-bottom 96 well plates. Samples were pelleted and resuspended in 50μL FACS block (PBS + 0.2% Triton + 3% BSA). After 10min incubation at room temperature antibodies were added to the sample and incubated overnight at 4°C. Cells were pelleted at 700rcf for 5min and resuspended in 50μL FACS block. One additional wash was performed before acquisition on a Fortessa flow cytometer (BD) equipped with a high-throughput sampler using FACSDiva software. Commercially available conjugated antibodies used were SOX2-V450 (Mouse, BD-561610), SOX1-PerCP-Cy.5 (Mouse, BD 561549), CDX2-PE (Mouse, BD 563428), BRA-APC (Goat, R&D IC2085A). Anti TBX6 (Goat, R&D AF4744) was conjugated to AlexaFluor-488, and anti FOXC2 (Goat, R&D AF6989) was conjugated to AlexaFluor-647 using Molecular Probes Antibody Labelling Kit (Invitrogen/ ThermoFisher). When labelling OTX2, samples were incubated with unconjugated OTX2 antibody (Goat, R&D AF1979) for 1 hour at room temperature in FACS block. Following 2 washes with FACS block AlexaFluor-488 anti-goat antibody (1:1000) was included in overnight incubations with fluorescently conjugated mouse antibodies.

### Flow cytometry data pre-processing

FCS files were imported into the R programming environment using the ‘flowCore’ package (Hahne et al., 2009). Individual cells were gated using a standard two-step gating strategy with an additional gate to remove outlier events with especially low fluorescence values as shown in (Figure S4). A custom R script was used to remove outlier events with fluorescent values exceeding a 3-sigma threshold from the mean of the average calculated across all samples of a given experiment. Raw fluorescence-intensity values were used in all downstream analysis and graphical representations of the data.

---

**Figure S1.**
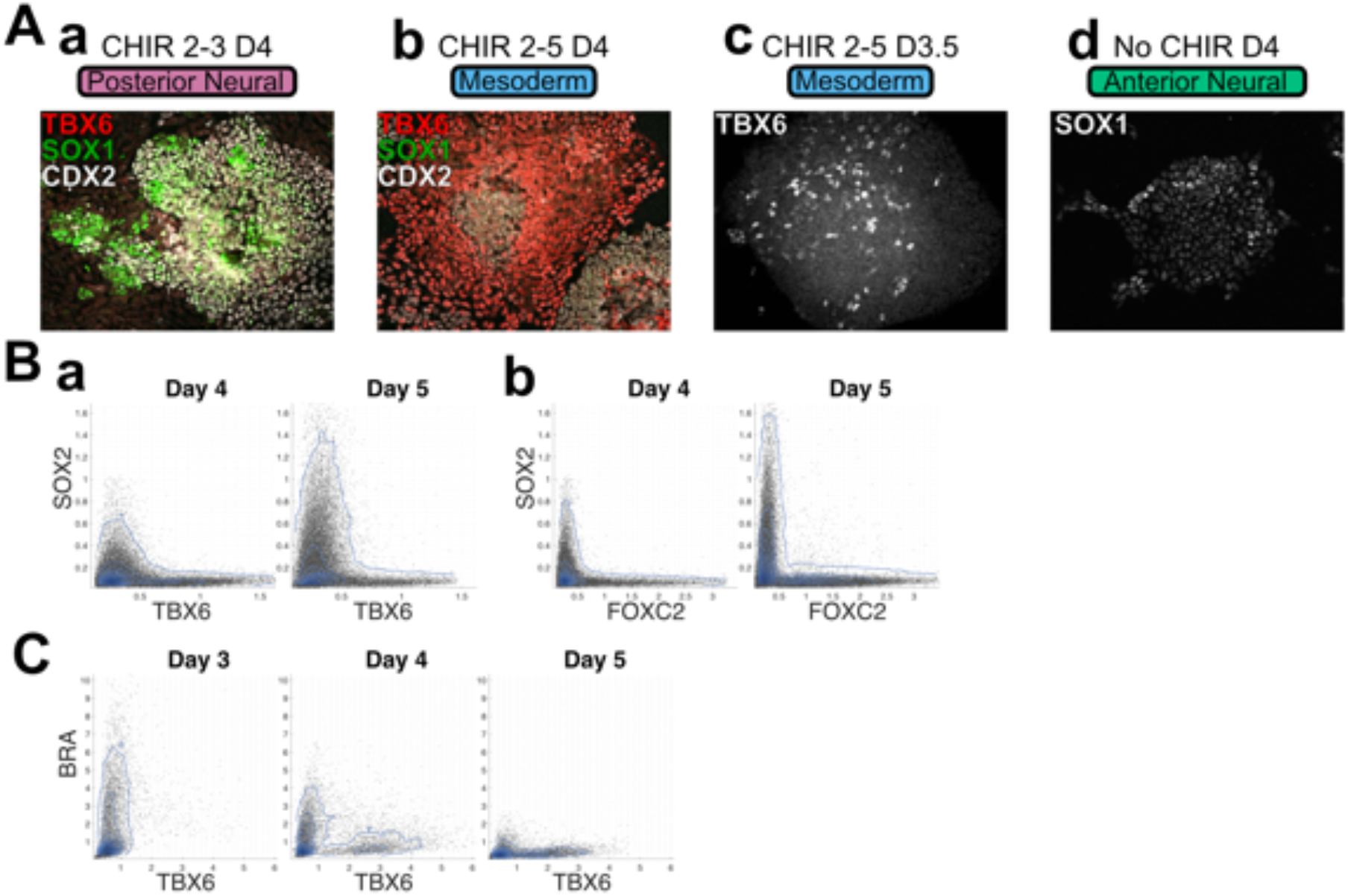
(A) Immunofluorescence assays indicate that CDX2 expressing CE progenitors at D4 adopt either SOX1 expressing posterior neural identity under the transient WNT signalling regime (Aa), or TBX6+ paraxial mesoderm identity under the sustained CHIR signalling regime (Ab). The expression of both TBX6 (Ac) and SOX1 (Ad) is initially not spatially organized. (B) Flow cytometry analysis of individual progenitors indicates that SOX2 levels are inversely correlated with both TBX6 (Ba) and FOXC2 (Bb) over the period of paraxial mesoderm differentiation. (C) Flow cytometry analysis indicates that the increase in the proportion of TBX6+ paraxial mesoderm progenitors over time is correlated with the loss of BRA+ paraxial mesoderm precursors.

**Figure S2.**
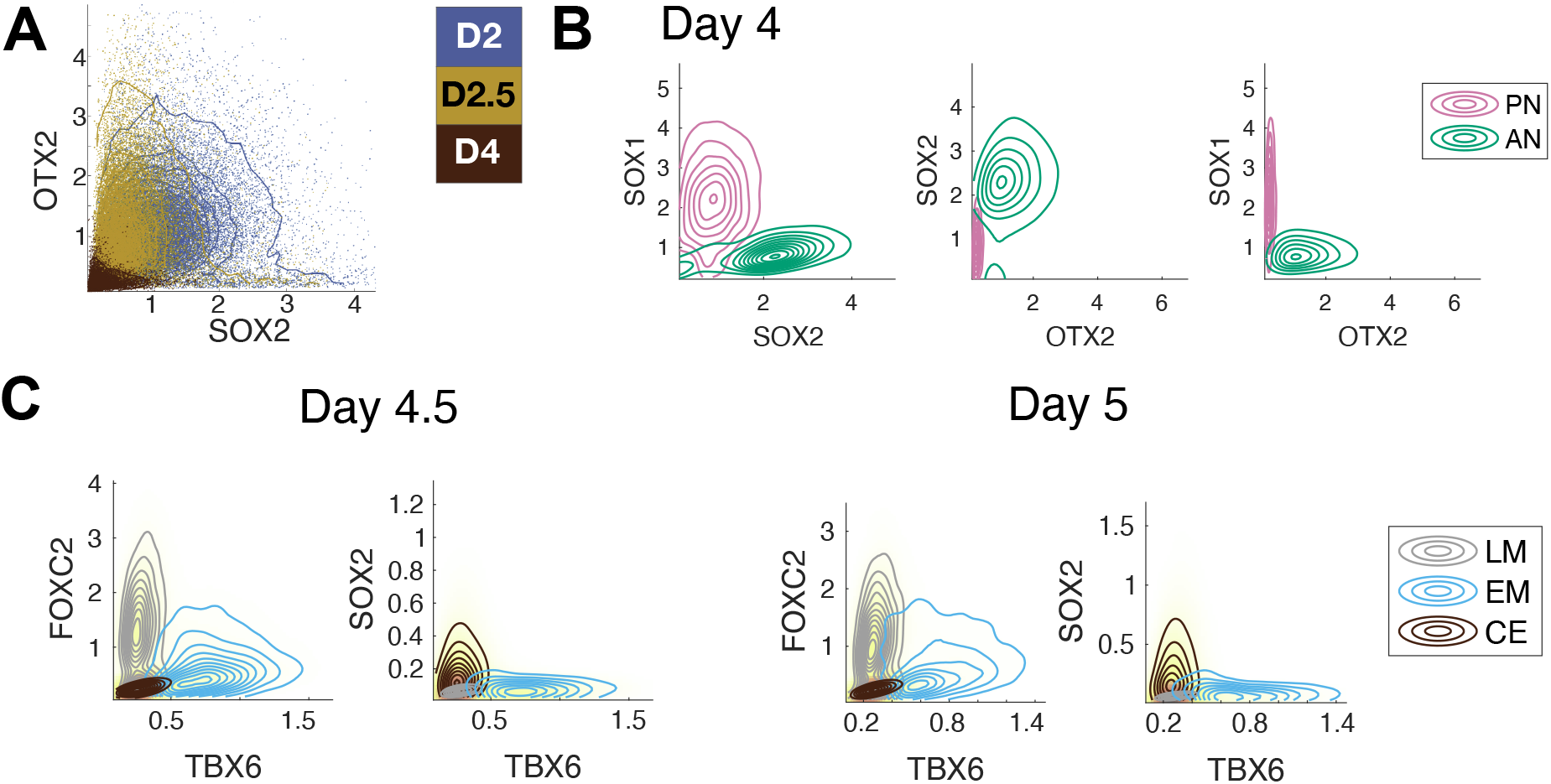
(A) Flow cytometry analysis of individual progenitors exposed to CHIR from D2 indicates SOX2 expression is reduced to intermediate levels by D2.5, whereas OTX2 remains expressed. SOX2 is further downregulated by D4 under the sustained CHIR by which point OTX2 expression is also repressed. (B) Clustering of the reference set of D4 samples with markers CDX2, OTX2, SOX1, SOX2. The clustering method identifies OTX2+ AN and OTX2 - PN clusters which express distinct levels of SOX1 and SOX2 with distributions that are almost identical to those obtained by the clusters defined with the minimal set of 5 markers used for model-fitting (Fig. 2B). (C) Clustering of D4.5 and D5 samples exposed to continuous CHIR induction assayed with CDX2, FOXC2, SOX1, SOX2, TBX6. The clustering identifies a late mesodermal population expressing high levels of FOXC2 and very low levels of SOX2.

**Figure S3.**
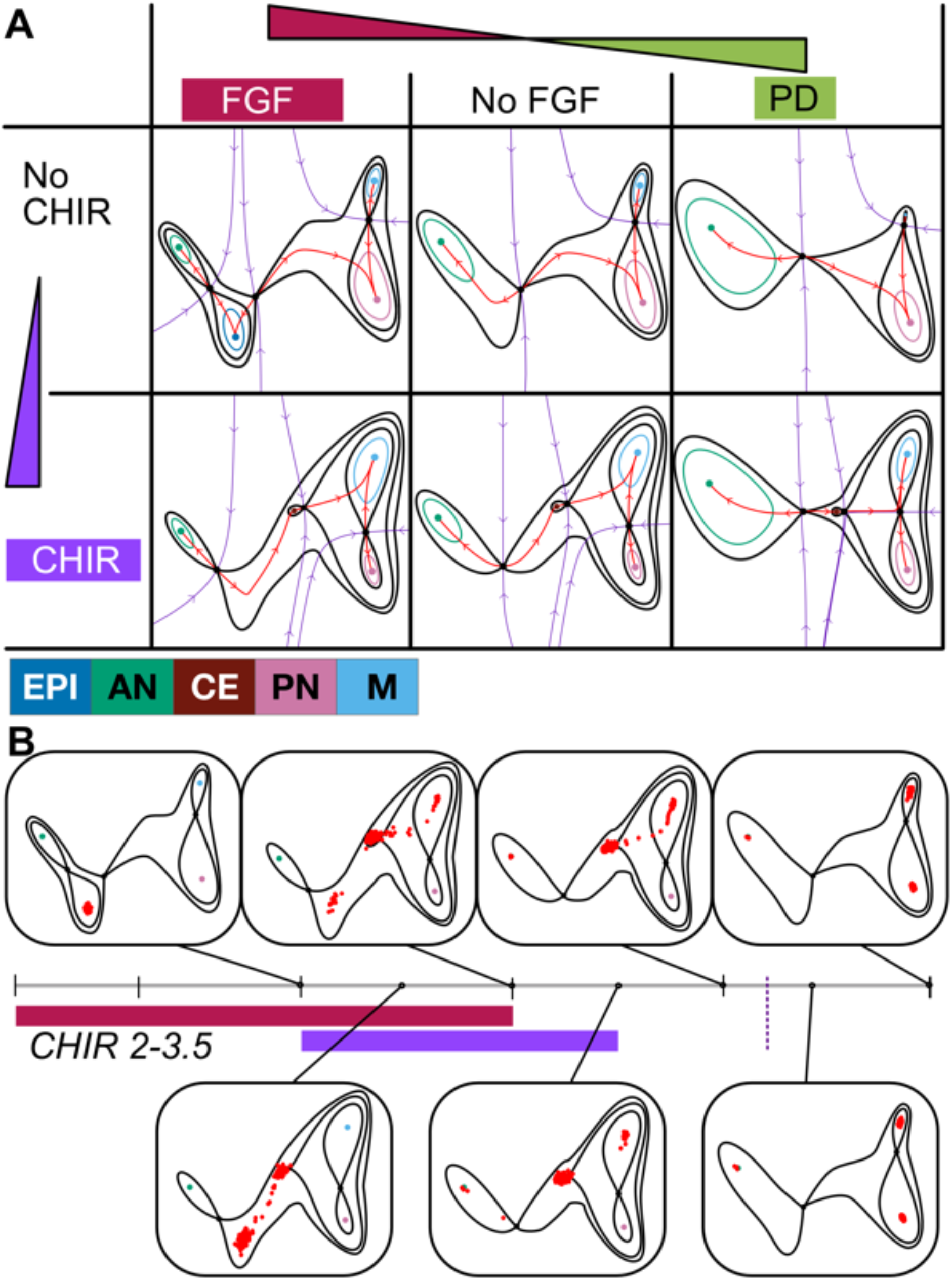
Landscape geometry and the effect of signalling after initial fitting. A. The landscapes produced by different combinations of signals are portrayed in the table. The parameters obtained by the initial fitting are used for the plot. The changes in the landscape that result from different signal combinations arise by bifurcations of attractors and flips in dynamical trajectories (unstable manifolds) of the parameterised landscape family. Note that the landscape corresponding to No CHIR+PD (top right) is an extrapolation of the fitting and not based directly on data, it therefore represents an untested prediction of the model (see Fig. 10). Colours correspond to cell identities as detailed. B. Example of a simulation time series of the model for the signalling regime CHIR 2-3.5. Red points represent the location of cells in the landscape at the specified time points. Cells are initialised in the EPI attractor at D2. Their location evolves as given by the stochastic dynamical system defined by the landscape in Fig. 4C, which precise geometry is determined by the signalling regime. Three changes in the landscape are apparent.

**Figure S4.**
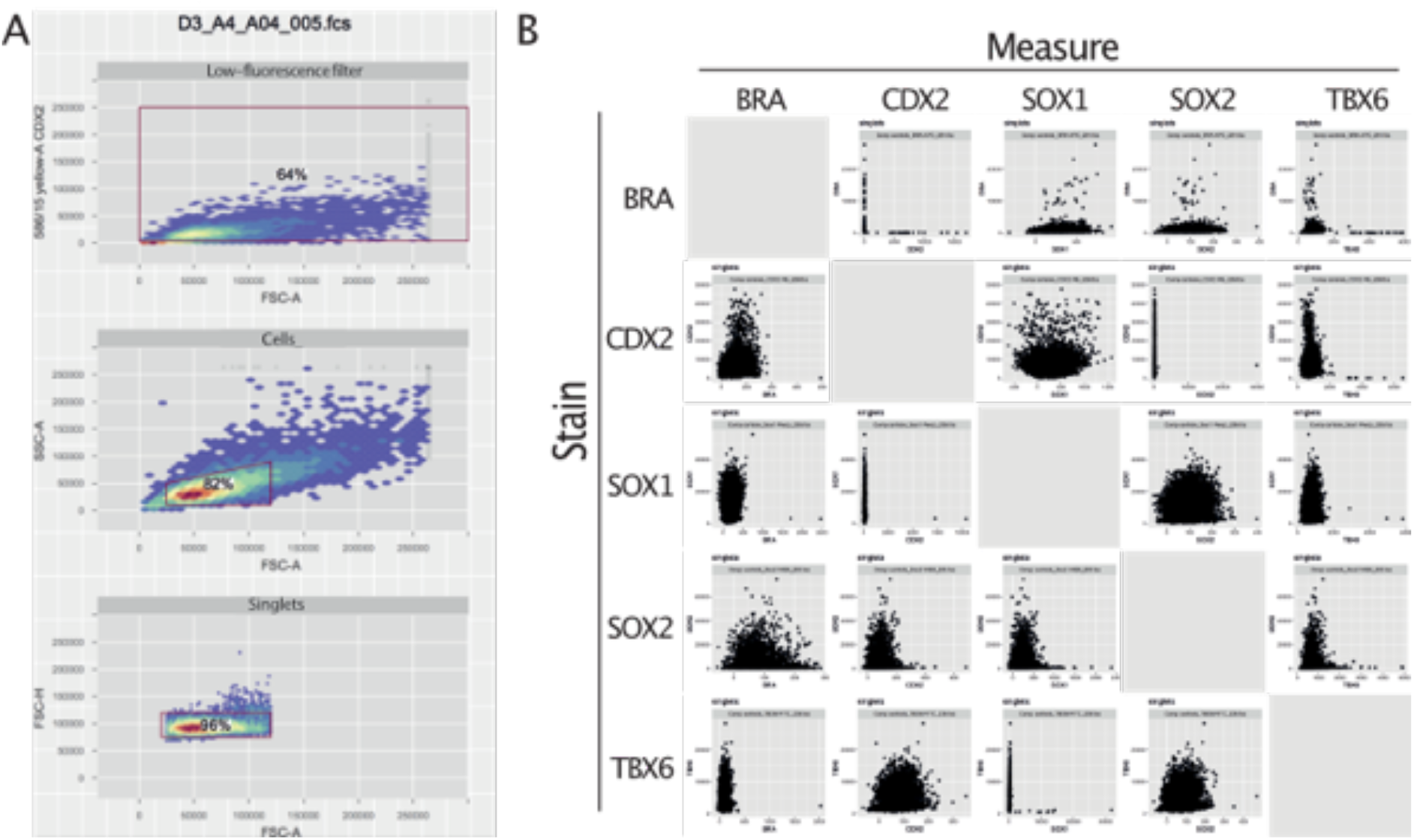
(A) ‘FlowCore’ gating series applied to flow-cytometry data to identify single-cells for analysis. Removing events with extremely low values in the 586/15 channel improved subsequent clustering. (B) Fluorescence measurements from samples individually stained with each of the 5 markers were acquired in all analysis channels and compensation was applied to account for the spectral overlap between fluorochromes. Appropriate compensation is demonstrated by the absence of spurious correlations between marker signals.

