## Supplementary Information for "A quantitative landscape of cell fate transitions identifies principles of cellular decision-making"

March 5, 2021

### 1 Landscape models

It is commonly accepted that cell fate decisions are made by gene regulatory networks comprising transcription factors and intercellular signals that define a complex stochastic dynamical system. These regulatory networks are usually modelled by mass-action equations: A system of ordinary differential equations which relates the evolution of concentrations of the mRNAs and proteins of the genes involved. Such systems require a lot of conjecture since it is very rare that more than a small proportion of the interactions are known in any detail. Moreover, they typically involve large numbers of parameters (notably rate constants) for which reliable values are difficult to obtain.

On the other hand, the transitions observed in developmental systems are typically between relatively small numbers of possible outcomes and for many systems we are essentially interested about the transitions between stationary states. Thus we can expect that although the regulatory network is complex, the resulting dynamics is relatively simple.

Things are also further simplified by using the idea of a generic property [1, 2] , a precise notion that was invented by algebraic geometers and topologists but which will also be very useful in biology. The assumption of only studying systems with given generic properties and the consequent avoidance of having to deal with aberrant, atypical, unlikely and/or degenerate systems eases the progress in understanding and classification. At the same time it seems very reasonable to assume that biological systems are also unlikely to possess such non-generic properties. Such a genericity assumption will underlie our choice of landscape structure.

If we demand that a dynamical system is quasi-gradient (i.e. has a finite number of restpoints and no oscillating or other more complex recurrent behaviour) and also insist that it is structurally stable (i.e. does not change its qualitative form when the parameters are perturbed) then they have a precise mathematical characterisation and these systems are called Morse-Smale. They always have a hierarchical structure reminiscent of Waddington landscapes [3, 4].

We are interested not just in a single dynamical system but a parameterised family where the parameters $\theta$ can be understood as functions of the signals that the cell is receiving. We call these parameterised landscapes (PLs).

Then the cellular transitions are described by bifurcations of the PL. The bifurcation set divides the parameter space up into components consisting of topologically equivalent quasi-gradient Morse-Smale systems. We call these MS-components. These are separated by components of the bifurcation set that in general have a stratified structure where codimension 1 hypersurfaces join together in sets of higher codimension. In the case of 2 and 3 parameters, which we consider here, the hypersurfaces are curves and surfaces respectively.

A combination of catastrophe theory and dynamical systems approaches (e.g. [5, 6, 7]) allows one to characterise the local structure of these bifurcations subject to a mild and precise genericity assumption if the number of parameters is no greater than 5.

For example, in 2 parameter PLs generically the bifurcation set will consist of curves of fold points with a finite number of cusps on them. Not only does the theory provide a qualitative description of the local behaviour but it also provides a normal form. To explain this, suppose that the "true" model has state variables $x$ and parameters $\theta$. The normal form will have state variable $y$ and parameters $\mu$ and the two will be related by a functional relationship of the form

$$y = \phi(x, \theta), \quad \mu = \psi(\theta) \tag{1}$$

so that a trajectory $x(t)$ of the true model in the $x$ coordinates will correspond to a trajectory $y(t) = \phi(x(t), \theta)$ in the $y$ coordinates of the normal form. Moreover, there exists a normal form such that the state variable $y$ is two dimensional.

These results are only local but for 2 parameter PLs we can use the local results and some topology to deduce a characterisation of global structures [4]. In Sect. 3.1 we describe how we use this to hypothesise the qualitative form of the PL and its normal form. In particular we describe the MS components and their boundary.

The normal form will be related to the true model via a relation as in (1) and this opens up the possibility of fitting the biological data to the normal form because summary statistics of the true model and the normal form should be the same. Therefore, if we can fit the normal form to the experimental data, then we have determined the qualitative form of the true model and those quantitative aspects of the true model that are preserved under a transformation as in (1).

### 2 Clustering

We developed a method to identify and quantify the populations present at different time points under different experimental conditions using the flow cytometry data collected from different samples, under different signaling conditions, at different time points. The proposed method allowed us to identify the proportions corresponding to the different cell identities present at each time point for each experimental series (ES). The method is based on the hypothesis that cell identities are attractors of a dynamical system, and therefore the population of cells with a particular identity should form an approximately multivariate normal distribution around the attractor because close to the attractor the dynamics are essentially linear.

In each each of the three experimental series (initial, test and prediction), we collected flow cytometry data for each temporal signaling profile. For the two main experimental series data were collected every 12h from day 2 to day 5. For the other experiments refer to Fig. 10 and Supp. Table T1 for the time points. For each temporal signaling profile $s$, within the experimental series $e$, and each time point $d$, we obtained a dataset $D_d^{e,s}$ using flow cytometry to measure the expression levels of the chosen core set of markers (BRA, CDX2, SOX1, SOX2, TBX6) at single cell level. Each data set involved 7200, 50000 or 10000 cells respectively. We refer to these datasets as sample datasets.

We defined a *reference set* of experiments consisting of the signaling regimes *No CHIR, CHIR 2-3 and CHIR 2-5.* These three experimental conditions were present in all our ES. We used the sample datasets $D_d^{e,s}$ corresponding to these signaling regimes in the reference set as a baseline to be able to compare the results between different experimental series. We refer to them as reference datasets.

The cell populations identified in these reference datasets were used to classify cells in other datasets (other samples, coming from other signaling profiles). Since we measured the expression level for 5 genes in each cell, the sample datasets consisted of a 5-dimensional vector for each cell (Figure SI1). That is, if $x \in D_d^{e,s}$ is the state of a cell, then $x \in \mathbb{R}^5$.

Firstly, for each time point $d$ and ES $e$ separately, data in the reference datasets for time point $d$ and ES were pooled, and the pooled dataset $\widehat{D^{e,d}}$ was clustered in the 5-dimensional data space. This gives a set of clusters $\{C_i^{e,d}\}$ for each time point $d$ and ES $e$ based just on the reference data. Using this, cells from all experiments were allocated to cell populations corresponding to the EPI, Tr, AN, CE, PN, EM and LM states described in the main paper (Fig. 2D) using the clusters for the appropriate time point and ES. The way in which cells were allocated to clusters is described below.

The reason for using only the reference data for clustering and doing it separately for each time point and ES is to use a consistent set of data across experimental series and time points (i.e. the reference set) while allowing flexibility to deal with batch effects (i.e. doing it separately for each time point and ES).

Note that we collected only one sample dataset at day 2 since all experimental conditions were equivalent at that point, so we had only one dataset corresponding to that day. For days 2.5 and 3 the pooled data contained two samples: CHIR or No CHIR (cells had been either exposed or not exposed to CHIR up to this point) and from day 3.5 onwards the pooled data contained 3 different sets of samples, each coming from one of the three conditions in the reference set.

For each time point $d$ separately we fitted a *Gaussian mixture model* (GMM) $Y^{e,d}$ with $m(d)$ components to the pooled reference data $\widehat{D^{e,d}}$ described above using the MATLAB algorithm `fitgmdist` ([8]). A GMM $Y$ consists of a weighted sum of $m$ multivariate normal distributions (MVNs). These MVNs have probability distributions $P_i(x)$, $i = 1, \ldots, m$ defined by their means $\mu_i$ and covariance matrices $\Sigma_i$.

The clusters $C_i^{e,d}$ are in 1-1 correspondence with the MVNs in the GMM and we regard a cell as belonging to one of the $m$ clusters $C_i^{e,d}$ if its 5-dimensional state has probability greater than some threshold probability $p$ for the corresponding MVN i.e. $P_i(x) > p$. In cases where a cell could belong to more than one cluster it was allocated to the one where it had the highest probability. That is, let $x \in \widehat{D^{e,d}}$ be a cell, then Once the reference data was clustered, we took each cluster and examined the scalar distributions of the single gene expression levels for each of the 5 genes to check if any of these were not unimodal. We started with $m = 1$ and increased $m$ until they were all unimodal. This gave the number of clusters $m = m(d)$ for that time point and ES. We wanted the minimal such $m(d)$.

As expected, the number of clusters $m(d)$ obtained at each time point $d$ varied. At intermediate days of the experiment we needed a higher number of clusters to obtain unimodal cluster distributions, while at early and late days this number was smaller.

By randomising the seed for the GMM algorithm we ensured that the MVNs found by the GMM algorithm and the determination of $m(d)$ were not biased by knowledge of previously found clusters. The goal was to identify the populations present in the data and avoid any bias introduced by previous knowledge of the system.

We then analysed the protein levels in each cluster and used this to label it with the cell identity it best represented. For certain days, more than one cluster was labelled with the same cell identity. Several considerations were necessary while performing

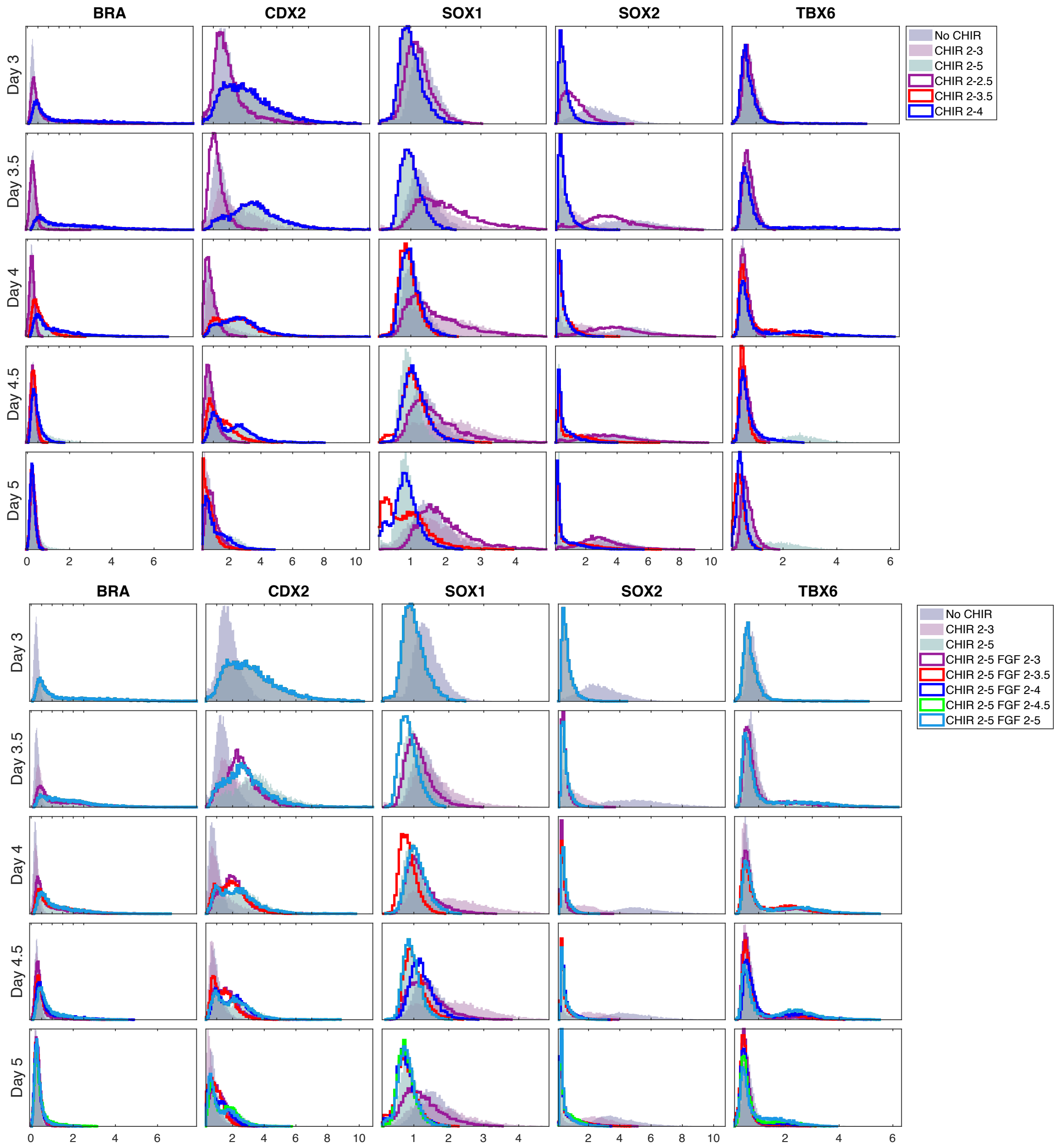

Figure SI1: Distributions comparing the three reference experimental conditions with the distributions for the other 8 experimental conditions in the initial experimental series. Shaded distributions correspond to the reference set, line distributions correspond to non-reference conditions as detailed in the legend. The reference set values cover all populations present in the different conditions.

this step and we discuss them in section 2.1. The fact that the clusters obtained by the automatic algorithm are coherent with the already known cell identities is a validation of the method used.

Finally, we used the MVNs $P_i$ obtained from computed from $\widehat{D_{e,d}}$ to allocate a cell identity to all cells from all samples. For cells from a given time point $d$ and ES $e$ the corresponding MVNs for that time point and ES were used, and the cell was considered for membership of a cluster if the posterior probability of the cell state wrt

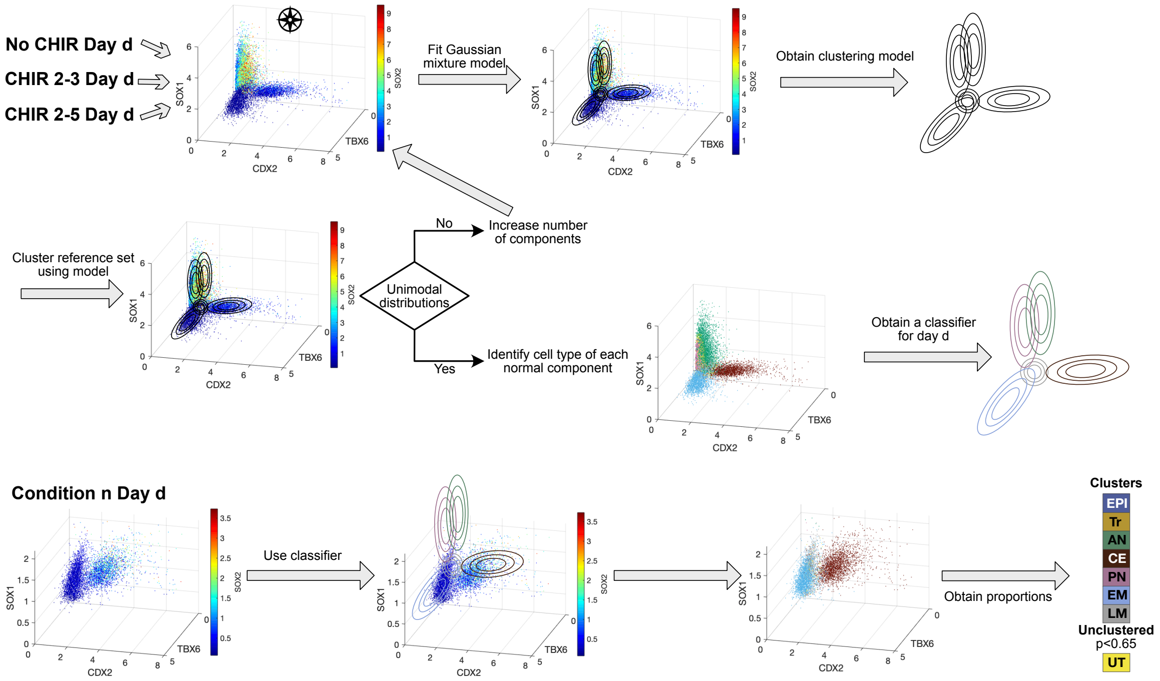

Figure SI2: Method used to perform the clustering as detailed in the text.

that MVN was greater than $p = 0.65$. The cell would then be added to the cluster maximising this probability. That is, if $x \in D_d^{e,s}$ is the state of a cell, then the cell belongs to $C_i^{e,d}$ if and only if $P_i(x) > p$ and $P_i(x) > P_j(x)$ for all $j \neq i$.

The number of cells in each cluster was stable in relation to the chosen threshold probability $p$ as shown in Fig. SI3. Any cells with a lower probability for all clusters remained as unclassified and we considered them as transitioning between clusters (UT).

Once the cells had been allocated to clusters, we computed the population proportions at each day for that experimental condition by adding up the proportions in all clusters corresponding to the same cell identity (EPI, Tr, AN, CE, PN,M,UT) as detailed in section 2.1. The proportions for each of the datasets corresponding to each cluster can be found in Supplementary Table T2 together with the aggregated proportions for each cell identity. These proportions are the data that we used to construct and parametrise the landscape model.

For the reference set of the *Predictions 2* experimental series (Fig. 10D and Supp. Table T1) only samples for days 3, 4 and 5 were collected. For some of the experimental conditions (CHIR 2-5 PD/RA 3.5-4.5) samples corresponding to day 4.5 were collected, but data for that day was not present in the reference conditions. Therefore, to cluster the day 4.5 samples we used the GMM model obtained for day 4 samples.

#### 2.1 Fate assignment

We performed the clustering method described above in datasets coming from three different ES (Initial, Test and Prediction). The details of the experimental conditions used in each ES can be found in Table T1. Figure SI4 shows the cumulative distributions for each cluster in the data pooled from the reference set for the different ES

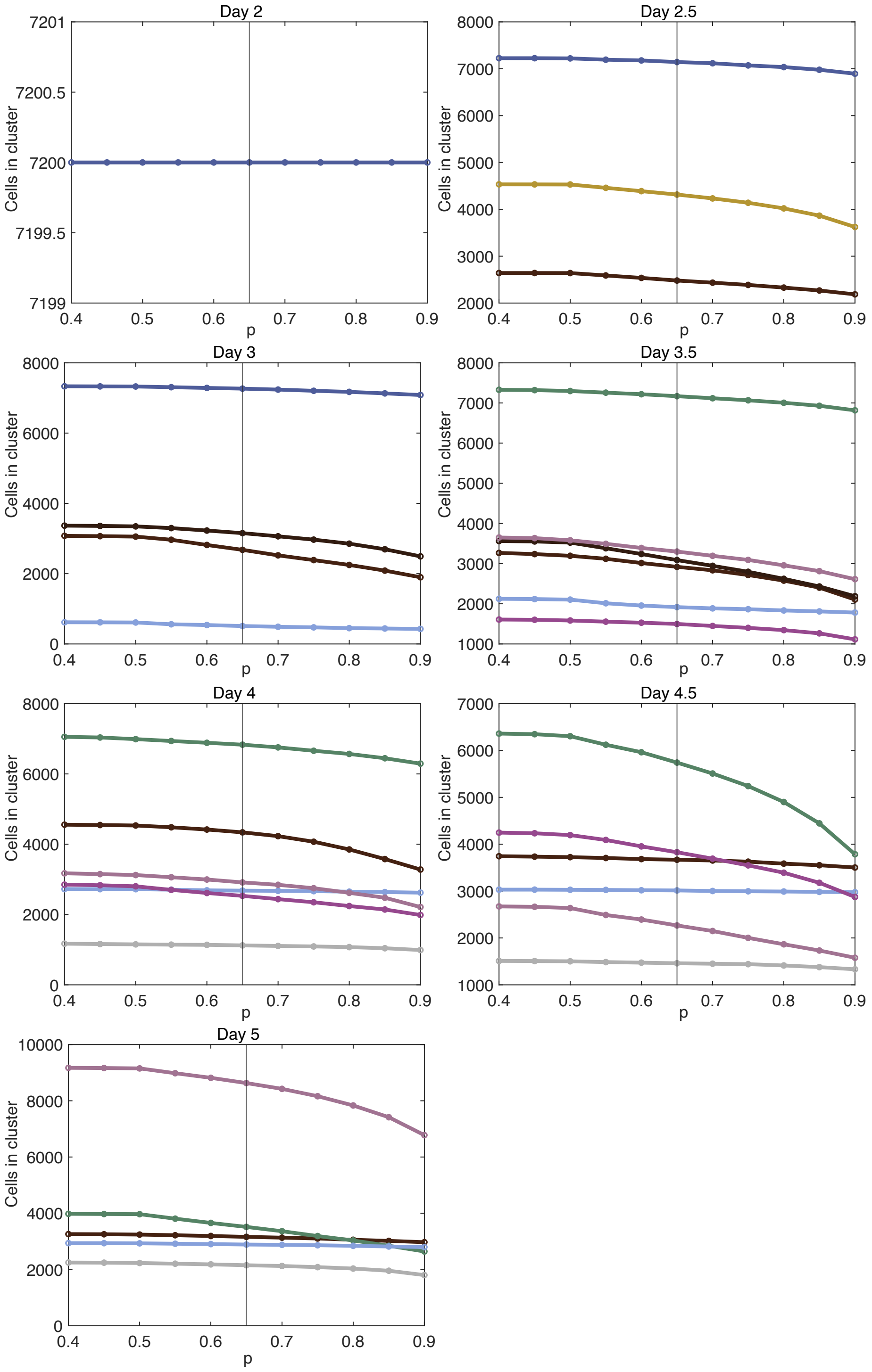

Figure SI3: Analysis of the number of cells from the three reference conditions (pooled data) in each cluster of each day. The $x$-axis corresponds to the minimal posterior probability $p$ chosen to perform the clustering. The colours of the lines correspond to the cell identity assigned to the cluster as in Fig. SI4.

with the corresponding identity assigned to each cluster. The main criteria to assign a cell identity was the levels of protein expression as detailed in Fig. 2C in the main text. The ideal situation would be that the algorithm finds one GMM component for each known cell identity in each day with a clear expression of the known markers. This was not the case and we dealt with the different situations encountered while assigning cell identities.

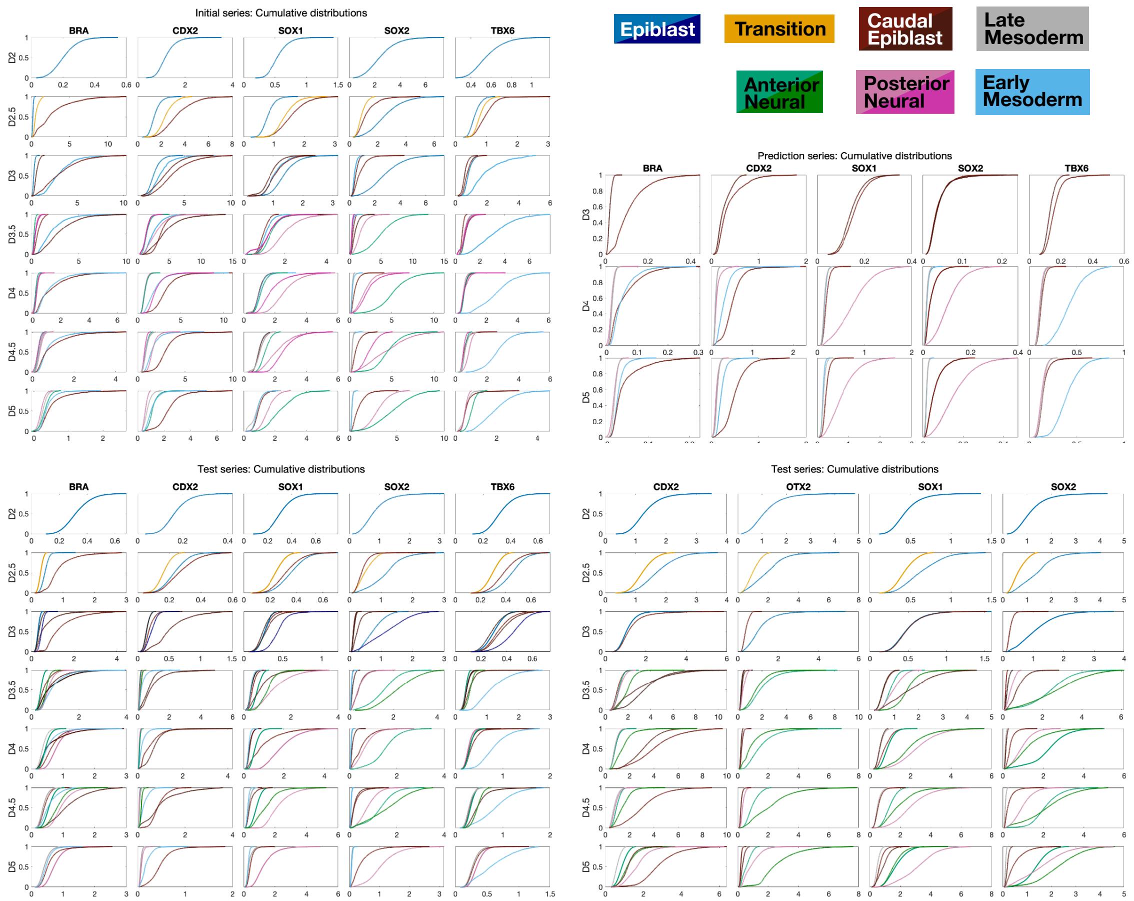

Figure SI4: Cumulative distributions for each cluster in each reference set for the different experimental series for the different markers. Colours correspond to the fate assigned to that cluster. When multiple clusters are assigned to the same identity different shades of colour are used. The cumulative plots show an S shape when the distribution is unimodal. The later the S raises, the higher the mean value for the distribution is.

**Transitioning population at day 2.5** At day 2.5 under CHIR induction, a population of cells with medium levels of SOX2 and low levels of all other proteins in the core set of markers (BRA, CDX2, SOX1, TBX6) was detected, but it was no longer detected at day 3. As discussed in the main paper, this population expressed high levels of OTX2 (see Fig. 1). We classified the cells in this cluster as transitioning cells between EPI and CE (Tr). We rationalised them as cells that were abandoning the EPI attractor but had yet to down-regulate OTX2 and up-regulate CDX2 or BRA.

**Anterior vs posterior neural** As discussed in the main text and shown in Fig. 1 and Supp. Fig. S2, we were able to distinguish AN from PN by the levels of expression of SOX1 and SOX2. We confirmed this observation by performing the clustering using the core set of markers (BRA, CDX2, SOX1, SOX2, TBX6) and an alternative set (CDX2, OTX2, SOX1, SOX2) in the Test experimental series (Supp. Fig. S2). The proportions obtained with the two different sets for the two neural identities were very similar (see Table T2). To further validate this distinction, we analysed the clusters that were characterised by high levels of SOX1 and SOX2 and observed that at days 3.5 and 4 the cells from No CHIR conditions samples were assigned to one cluster and the cells from CHIR 2-3 conditions samples to a different one. Since we expected that under No CHIR conditions only AN cells would be present and under CHIR 2-3 conditions mostly PN cells would be present. This validates our distinction.

At days 4.5 and 5 the levels of SOX1 and SOX2 became much more similar and the separation of the AN/PN clusters was lost. Therefore, we assumed that no more cells would becoming AN after day 4 (since the EPI population had disappeared) and that any increase in the neural population corresponds to an increase of PN cells.

**More than one cluster for the same identity** As shown in Fig. 2C, for the initial experimental series, at days 3 and 3.5 there were two separate clusters assigned to the CE identity. In the pooled data corresponding to these days, a population with high levels of both BRA and CDX2 (NMP) and a population with high levels of CDX2 only (other CE cells) were identified. Both clusters corresponded to CE cells (multiple brown curves in Fig. SI4).

There was also more than one cluster assigned to PN on the intermediate days (multiple pink curves in Fig. SI4). At days 4.5 and 5, since the levels of SOX1 and SOX2 of AN and PN had become very similar it was difficult for the algorithm to find well defined clusters and it found an extra cluster for neural at day 4.5. At day 3.5 the second PN cluster is small with low levels of all proteins except SOX1, which prompted us to define it as PN. At day 4 the second PN cluster has a higher level of CDX2 compared with other neural populations, which may mean that it was defined by a large transitioning population that had been captured by the algorithm as a separate cluster. The levels of SOX1 and SOX2 justify the definition of PN identity over CE.

**Late Paraxial Mesoderm** As mentioned in the main text, late paraxial mesoderm cells downregulate TBX6 and upregulate FOXC2. Since FOXC2 is not part of our core set of markers, this population defines a cluster with low levels of all markers (LM). It was detected in data from day 4 and it became more abundant in data from consecutive days (Fig.1 and Supp. Fig. 2 and 3).

As part of the test experimental series we ran the CHIR 2-5 samples for a set of markers including FOXC2 (i.e. CDX2, FOXC2, SOX1, SOX2, TBX6). Here, the clustering identified a cluster positive in FOXC2 and negative for all other markers (Supp. Fig. 2). The population proportions for this cluster is almost identical to the proportions for the LM cluster obtained using the core set of markers. In this case, we didn't have a complete reference set of experimental conditions so the clusters were used only for comparison.

We combine the proportions for Early Mesoderm (EM) and Late Mesoderm (LM) to quantify the population of Mesoderm cells (M) used in the construction of the landscape.

Table SI1: Proportions of LM under CHIR 2-5 conditions using two different sets of markers: core set (BRA, CDX2, SOX1, SOX2, TBX6) or a set including FOXC2 (CDX2, FOXC2, SOX1, SOX2, TBX6)

| Day | Core set | With FOXC2 |
|---|---|---|
| 4 | 0.11 | 0 |
| 4.5 | 0.37 | 0.33 |
| 5 | 0.47 | 0.43 |

#### 2.2 Cluster structure: local correlations

We analysed the structure of the clusters obtained in order to test our cell identity assignment. We considered the pooled data from the reference set in the initial experimental series at each day separately. For each day and cluster we considered the cells that had been assigned to it by the clustering algorithm with a minimum probability of 0.65. Figures SI5 and SI6 show the mean value and principal component analysis of all clusters identified at different days. We noticed that each principal component has only a small number of non-zero weights for the different proteins, and that most of the variance is explained by the first principal component alone. Moreover, the weights are clearly different for different cell identities and the highest weights in the first component correspond to the highly expressed proteins in each case.

Next, we analysed the normality of the clusters obtained. We checked that the structure of the cluster remains constant along the cluster. We computed the correlation structure of the cells in each cluster. In particular, to test whether these clusters were indeed approximately multivariate normal, we analysed the local correlations. We computed a smoothing spline that approximates the cluster data using the first PC as parametrising variable. We ordered the points according to the position of their projection along the spline. Once ordered, we split them in 25 bins (Figure SI7). For each bin we computed the gene-to-gene correlations of the cells that fell in it.

If the cluster is indeed a multivariate normal, these bins should have a similar structure along the first PC, giving similar correlation values. The results of this computation show that these local correlations take stable values along the spline for all clusters in all days, which supports the hypothesis that the clusters are multivariate Gaussian distributions corresponding to well defined cell states (Fig. SI8). Moreover, the values obtained are very different for different clusters (Fig. SI9 and SI10), with clusters corresponding to the same cell identity on different days giving similar values in general.

The computation of these local correlations along the first PC corresponds to the computation of the gene-to-gene correlation after projecting on the orthogonal of the first PC (Fig. SI7). Since we are conditioning the correlation on a value along the first PC, it is similar to removing the information from the first PC and looking at the relationships between the genes in the complementary space (PC2, . . . , PC5). We show the values of this computation in Figure SI9. We observe that when the local values are tight, the correlation for the projected data has a very similar value. When the variance of the local correlations is larger the correlation for the projected data takes a more extreme value than the local ones.

For example, EPI and AN show very similar local correlations along all days until

Day 2 EPI

| | | | | | | |
|---|---|---|---|---|---|---|
| VE | | 0.88 | 0.10 | 0.01 | 0 | 0 |
| BRA | 0.21 | 0.02 | 0.09 | 0.08 | 0.59 | 0.80 |
| CDX2 | 1.21 | 0.13 | 0.91 | -0.38 | -0.10 | 0 |
| SOX1 | 0.55 | 0.04 | 0.25 | 0.76 | -0.52 | 0.28 |
| SOX2 | 2.51 | 0.99 | -0.15 | -0.01 | -0.01 | 0 |
| TBX6 | 0.56 | 0.05 | 0.27 | 0.52 | 0.60 | -0.54 |

Day 2.5 EPI

| | | | | | | |
|---|---|---|---|---|---|---|
| VE | | 0.85 | 0.13 | 0.02 | 0 | 0 |
| BRA | 0.19 | 0.02 | 0.06 | 0.03 | 0.51 | 0.86 |
| CDX2 | 1.47 | 0.20 | 0.89 | -0.39 | -0.13 | 0.02 |
| SOX1 | 0.80 | 0.08 | 0.30 | 0.85 | -0.39 | 0.18 |
| SOX2 | 2.56 | 0.97 | -0.23 | -0.02 | -0.01 | 0 |
| TBX6 | 0.64 | 0.08 | 0.26 | 0.35 | 0.76 | -0.48 |

Day 2.5 Tr

| | | | | | | |
|---|---|---|---|---|---|---|
| VE | | 0.67 | 0.13 | 0.11 | 0.09 | 0.01 |
| BRA | 0.62 | 0.12 | 0.92 | 0.08 | 0.37 | -0.03 |
| CDX2 | 2.39 | 0.91 | -0.11 | -0.39 | 0.06 | -0.03 |
| SOX1 | 1.39 | 0.31 | 0.13 | 0.61 | -0.59 | -0.41 |
| SOX2 | 0.87 | 0.18 | -0.35 | 0.62 | 0.68 | -0.03 |
| TBX6 | 0.81 | 0.18 | 0.08 | 0.29 | -0.23 | 0.91 |

Day 2.5 CE

| | | | | | | |
|---|---|---|---|---|---|---|
| VE | | 0.70 | 0.21 | 0.05 | 0.02 | 0.01 |
| BRA | 2.89 | 0.99 | 0.08 | 0.09 | -0.03 | 0.01 |
| CDX2 | 3.39 | -0.07 | 0.98 | -0.16 | -0.13 | 0 |
| SOX1 | 1.58 | 0 | 0.15 | 0.25 | 0.82 | -0.49 |
| SOX2 | 1.27 | -0.11 | 0.11 | 0.95 | -0.27 | 0.05 |
| TBX6 | 0.98 | -0.01 | 0.07 | 0.08 | 0.48 | 0.87 |

Day 3 EPI

| | | | | | | |
|---|---|---|---|---|---|---|
| VE | | 0.75 | 0.21 | 0.03 | 0.01 | 0 |
| BRA | 0.26 | 0.03 | 0.06 | -0.01 | 0.55 | 0.83 |
| CDX2 | 1.78 | 0.18 | 0.85 | -0.45 | -0.19 | 0.06 |
| SOX1 | 1.39 | 0.14 | 0.38 | 0.88 | -0.22 | 0.12 |
| SOX2 | 2.82 | 0.97 | -0.24 | -0.06 | -0.02 | 0 |
| TBX6 | 0.83 | 0.09 | 0.26 | 0.14 | 0.78 | -0.54 |

Day 3 CE1

| | | | | | | |
|---|---|---|---|---|---|---|
| VE | | 0.92 | 0.05 | 0.02 | 0.01 | 0 |
| BRA | 0.55 | 0.06 | 0.45 | 0.87 | 0.19 | -0.10 |
| CDX2 | 2.98 | 0.99 | -0.15 | 0.01 | -0.03 | 0.02 |
| SOX1 | 0.91 | 0.12 | 0.72 | -0.37 | -0.29 | -0.49 |
| SOX2 | 0.52 | 0.06 | 0.19 | -0.31 | 0.93 | -0.03 |
| TBX6 | 0.60 | 0.05 | 0.47 | -0.13 | -0.11 | 0.86 |

Day 3 CE2

| | | | | | | |
|---|---|---|---|---|---|---|
| VE | | 0.52 | 0.44 | 0.03 | 0.01 | 0 |
| BRA | 3.14 | 0.93 | 0.36 | 0.04 | -0.07 | -0.01 |
| CDX2 | 3.89 | -0.36 | 0.92 | -0.13 | -0.07 | 0 |
| SOX1 | 1.09 | 0 | 0.10 | 0.30 | 0.77 | -0.55 |
| SOX2 | 0.87 | -0.09 | 0.07 | 0.93 | -0.36 | 0.02 |
| TBX6 | 0.75 | 0.01 | 0.07 | 0.18 | 0.52 | 0.83 |

Day 3 EM

| | | | | | | |
|---|---|---|---|---|---|---|
| VE | | 0.60 | 0.23 | 0.15 | 0.02 | 0 |
| BRA | 3.10 | 0.88 | -0.21 | -0.42 | -0.02 | 0 |
| CDX2 | 2.55 | 0.46 | 0.41 | 0.77 | -0.15 | -0.07 |
| SOX1 | 1.07 | 0.08 | 0.10 | 0.07 | 0.97 | -0.21 |
| SOX2 | 0.42 | 0.05 | 0.04 | 0.07 | 0.20 | 0.98 |
| TBX6 | 1.98 | -0.02 | 0.88 | -0.47 | -0.05 | 0.01 |

Day 3.5 AN

| | | | | | | |
|---|---|---|---|---|---|---|
| VE | | 0.92 | 0.07 | 0.01 | 0 | 0 |
| BRA | 0.29 | 0.02 | 0.07 | -0.02 | 0.59 | 0.80 |
| CDX2 | 1.48 | 0.04 | 0.86 | -0.44 | -0.22 | 0.08 |
| SOX1 | 1.34 | 0.10 | 0.40 | 0.89 | -0.17 | 0.11 |
| SOX2 | 5.13 | 0.99 | -0.09 | -0.07 | -0.02 | 0 |
| TBX6 | 0.79 | 0.04 | 0.29 | 0.08 | 0.76 | -0.58 |

Day 3.5 CE1

| | | | | | | |
|---|---|---|---|---|---|---|
| VE | | 0.83 | 0.09 | 0.05 | 0.03 | 0 |
| BRA | 0.72 | 0.10 | 0.99 | 0.04 | 0.01 | -0.06 |
| CDX2 | 3.72 | 0.99 | -0.10 | -0.09 | -0.08 | -0.01 |
| SOX1 | 0.94 | 0.10 | -0.06 | 0.61 | 0.69 | -0.37 |
| SOX2 | 0.58 | 0.02 | -0.03 | 0.75 | -0.66 | 0.03 |
| TBX6 | 0.56 | 0.06 | 0.04 | 0.22 | 0.30 | 0.92 |

Day 3.5 CE2

| | | | | | | |
|---|---|---|---|---|---|---|
| VE | | 0.62 | 0.32 | 0.03 | 0.02 | 0 |
| BRA | 3.01 | 0.12 | 0.99 | 0.07 | 0.03 | -0.02 |
| CDX2 | 4.46 | 0.99 | -0.11 | -0.04 | -0.03 | -0.01 |
| SOX1 | 1.27 | 0 | -0.04 | 0.62 | -0.59 | -0.51 |
| SOX2 | 0.96 | 0.04 | -0.08 | 0.68 | 0.72 | 0 |
| TBX6 | 0.86 | 0.01 | 0 | 0.37 | -0.35 | 0.86 |

Day 3.5 EM

| | | | | | | |
|---|---|---|---|---|---|---|
| VE | | 0.53 | 0.37 | 0.08 | 0.01 | 0 |
| BRA | 2.21 | 0.70 | -0.22 | -0.68 | -0.05 | 0.01 |
| CDX2 | 2.65 | 0.69 | -0.01 | 0.72 | -0.06 | -0.07 |
| SOX1 | 1.07 | 0.07 | 0.05 | -0.02 | 0.95 | -0.29 |
| SOX2 | 0.44 | 0.06 | -0.01 | 0.05 | 0.29 | 0.95 |
| TBX6 | 3.19 | 0.17 | 0.97 | -0.14 | -0.06 | 0.03 |

Day 3.5 PN1

| | | | | | | |
|---|---|---|---|---|---|---|
| VE | | 0.63 | 0.22 | 0.12 | 0.01 | 0 |
| BRA | 0.42 | 0.04 | 0.03 | 0.05 | 0.69 | 0.72 |
| CDX2 | 2.96 | 0.98 | -0.13 | 0.10 | -0.06 | 0.01 |
| SOX1 | 2.02 | -0.06 | 0.31 | 0.93 | -0.18 | 0.10 |
| SOX2 | 1.80 | 0.15 | 0.94 | -0.30 | -0.02 | 0 |
| TBX6 | 0.75 | 0.04 | 0.07 | 0.19 | 0.70 | -0.69 |

Day 3.5 PN2

| | | | | | | |
|---|---|---|---|---|---|---|
| VE | | 0.82 | 0.08 | 0.06 | 0.03 | 0.01 |
| BRA | 0.46 | 0.14 | -0.57 | -0.51 | -0.62 | -0.08 |
| CDX2 | 1.36 | 0.65 | -0.40 | 0.64 | -0.01 | -0.06 |
| SOX1 | 1.20 | 0.62 | 0.68 | -0.22 | -0.29 | -0.15 |
| SOX2 | 0.38 | 0.14 | 0.03 | -0.04 | -0.09 | 0.98 |
| TBX6 | 0.78 | 0.39 | -0.22 | -0.53 | 0.72 | -0.01 |

Figure SI5: Mean and principal components for all clusters identified in the initial dataset for days 2 to 3.5 days. Each panel corresponds to the cluster and day specified at the top. The first column contains the mean value for the cells in the cluster. Each of the other columns corresponds to a principal component and it is coloured to ease the reading. The intensity of the colour corresponds to the absolute magnitude of the weight. The first row contains the amount of variation explained by that component (VE) and the rows below contain the weights of each protein in that component.

day 4, at later days the AN and PN cells are difficult to differentiate and the clusters are not well distinguished. EM at days 3, 3.5 and 4 show very similar correlation structures. At days 4.5 and 5 the EM correlations change with respect to previous days but stay similar between them. The LM clusters are very similar on days 4 to 5. Clusters corresponding to PN and CE are harder to compare since there are several clusters in a day that correspond to the same identity. For example, at day 4 the two clusters labelled as PN show very similar local correlations except for CDX2 TBX6. There are visible similarities on some clusters corresponding to the same cell type in consecutive days.

To further validate the cluster structure and their differences we analysed the change in correlation along the different clusters in more detail. We took a sample corresponding to a particular day and experimental condition and we considered the two main clusters in it (those containing the largest proportions of cells). We took the cells classified as each of these main clusters and rotated the data so that the first coordinate expanded them (Fig. 2F). To do this we performed linear discriminant analysis of the cells labelled as these two main clusters and obtained a separating hyperplane $\Pi$. Then we rotated all data from that day and experimental condition so that the first component was perpendicular to $\Pi$. The other components were determined by the specific rotation. We then computed a spline going through the data and repeated the previous procedure: project points into the spline, order points according to the projection, bin the cells accordingly (30 bins) and compute the gene-to-gene correlation for each pair of genes in each bin. If we compare the result with the clustering we clearly see that the local correlation changes at the same region where the cluster identity changes. In experimental conditions with several clusters this analysis becomes harder. Restricting the analysis to cells belonging to the two main clusters in a given sample the change in cluster label corresponds to a change in the local correlations.

**Day 4 AN**

| | | | | | | |
|---|---|---|---|---|---|---|
| VE | | 0.95 | 0.03 | 0.02 | 0 | 0 |
| BRA | 0.21 | 0.02 | 0.05 | -0.02 | 0.71 | 0.70 |
| CDX2 | 0.86 | 0.05 | 0.82 | -0.50 | -0.22 | 0.16 |
| SOX1 | 1.11 | 0.11 | 0.48 | 0.86 | -0.09 | 0.08 |
| SOX2 | 5.22 | 0.99 | -0.11 | -0.07 | -0.03 | 0.01 |
| TBX6 | 0.59 | 0.05 | 0.29 | -0.04 | 0.66 | -0.69 |

**Day 4 CE**

| | | | | | | |
|---|---|---|---|---|---|---|
| VE | | 0.59 | 0.35 | 0.05 | 0.02 | 0 |
| BRA | 1.37 | 0.36 | 0.93 | -0.05 | -0.08 | -0.04 |
| CDX2 | 3.28 | 0.91 | -0.37 | -0.12 | -0.11 | 0 |
| SOX1 | 0.99 | 0.13 | 0.02 | 0.15 | 0.87 | -0.44 |
| SOX2 | 0.69 | 0.10 | -0.01 | 0.98 | -0.19 | 0 |
| TBX6 | 0.58 | 0.08 | 0.05 | 0.08 | 0.43 | 0.90 |

**Day 4 EM**

| | | | | | | |
|---|---|---|---|---|---|---|
| VE | | 0.55 | 0.36 | 0.07 | 0.02 | 0 |
| BRA | 1.22 | 0.56 | -0.34 | 0.76 | -0.04 | -0.01 |
| CDX2 | 2.02 | 0.78 | -0.05 | -0.61 | -0.13 | -0.07 |
| SOX1 | 1.02 | 0.12 | 0.06 | -0.02 | 0.95 | -0.27 |
| SOX2 | 0.38 | 0.09 | -0.02 | -0.05 | 0.26 | 0.96 |
| TBX6 | 2.68 | 0.24 | 0.94 | 0.24 | -0.08 | 0.03 |

**Day 4 PN1**

| | | | | | | |
|---|---|---|---|---|---|---|
| VE | | 0.59 | 0.35 | 0.04 | 0.01 | 0 |
| BRA | 0.23 | 0.02 | -0.02 | 0.28 | 0.95 | -0.12 |
| CDX2 | 0.85 | 0.09 | 0.10 | 0.88 | -0.30 | -0.33 |
| SOX1 | 2.65 | 0.60 | 0.78 | -0.15 | 0.04 | -0.04 |
| SOX2 | 1.84 | 0.79 | -0.61 | -0.02 | -0.02 | -0.01 |
| TBX6 | 0.52 | 0.07 | 0.05 | 0.34 | 0.02 | 0.94 |

**Day 4 PN2**

| | | | | | | |
|---|---|---|---|---|---|---|
| VE | | 0.51 | 0.28 | 0.20 | 0.02 | 0 |
| BRA | 0.28 | 0.02 | 0.04 | -0.01 | 0.28 | 0.96 |
| CDX2 | 2.18 | 0.02 | 0.93 | -0.36 | -0.04 | -0.03 |
| SOX1 | 1.84 | 0.28 | 0.33 | 0.90 | -0.08 | 0.01 |
| SOX2 | 2.24 | 0.96 | -0.12 | -0.25 | -0.03 | 0 |
| TBX6 | 0.60 | 0.05 | 0.05 | 0.05 | 0.96 | -0.28 |

**Day 4 LM**

| | | | | | | |
|---|---|---|---|---|---|---|
| VE | | 0.80 | 0.10 | 0.06 | 0.02 | 0.02 |
| BRA | 0.30 | 0.15 | 0.12 | 0.87 | -0.44 | -0.09 |
| CDX2 | 0.84 | 0.63 | 0.71 | -0.27 | -0.14 | 0.08 |
| SOX1 | 0.98 | 0.65 | -0.69 | -0.16 | -0.29 | -0.04 |
| SOX2 | 0.32 | 0.15 | -0.09 | 0.22 | 0.29 | 0.92 |
| TBX6 | 0.57 | 0.38 | -0.03 | 0.30 | 0.79 | -0.38 |

**Day 4.5 AN**

| | | | | | | |
|---|---|---|---|---|---|---|
| VE | | 0.94 | 0.04 | 0.02 | 0 | 0 |
| BRA | 0.27 | 0.02 | 0.05 | 0.07 | 0.90 | -0.42 |
| CDX2 | 0.86 | 0.10 | 0.27 | 0.91 | -0.20 | -0.25 |
| SOX1 | 1.36 | 0.11 | 0.93 | -0.33 | -0.07 | -0.11 |
| SOX2 | 4.58 | 0.99 | -0.14 | -0.07 | -0.01 | 0 |
| TBX6 | 0.60 | 0.06 | 0.21 | 0.25 | 0.37 | 0.87 |

**Day 4.5 CE**

| | | | | | | |
|---|---|---|---|---|---|---|
| VE | | 0.62 | 0.24 | 0.10 | 0.02 | 0.01 |
| BRA | 0.94 | 0.13 | 0.97 | 0.01 | -0.14 | -0.11 |
| CDX2 | 3.05 | 0.98 | -0.14 | -0.14 | -0.09 | 0 |
| SOX1 | 0.97 | 0.11 | 0.05 | 0.11 | 0.84 | -0.52 |
| SOX2 | 0.85 | 0.13 | -0.04 | 0.98 | -0.12 | 0.05 |
| TBX6 | 0.60 | 0.07 | 0.16 | 0.02 | 0.50 | 0.85 |

**Day 4.5 EM**

| | | | | | | |
|---|---|---|---|---|---|---|
| VE | | 0.62 | 0.28 | 0.07 | 0.03 | 0 |
| BRA | 0.81 | 0.36 | -0.49 | 0.80 | -0.03 | -0.03 |
| CDX2 | 1.59 | 0.64 | -0.48 | -0.59 | -0.13 | -0.08 |
| SOX1 | 0.97 | 0.17 | 0.03 | -0.03 | 0.96 | -0.22 |
| SOX2 | 0.34 | 0.08 | -0.07 | -0.04 | 0.21 | 0.97 |
| TBX6 | 2.66 | 0.66 | 0.73 | 0.14 | -0.13 | 0.03 |

**Day 4.5 PN1**

| | | | | | | |
|---|---|---|---|---|---|---|
| VE | | 0.84 | 0.12 | 0.03 | 0.01 | 0 |
| BRA | 0.36 | 0.01 | 0.02 | 0.06 | 0.44 | 0.90 |
| CDX2 | 1.59 | 0.03 | -0.10 | 0.96 | -0.25 | 0.06 |
| SOX1 | 2.19 | 0.04 | 0.99 | 0.09 | -0.04 | -0.01 |
| SOX2 | 4.38 | 1.00 | -0.03 | -0.04 | -0.03 | 0.01 |
| TBX6 | 0.84 | 0.04 | 0 | 0.26 | 0.86 | -0.44 |

**Day 4.5 PN2**

| | | | | | | |
|---|---|---|---|---|---|---|
| VE | | 0.63 | 0.33 | 0.02 | 0.01 | 0 |
| BRA | 0.26 | 0.02 | 0 | 0.31 | 0.94 | -0.14 |
| CDX2 | 0.74 | 0.10 | 0.09 | 0.88 | -0.34 | -0.30 |
| SOX1 | 2.60 | 0.39 | 0.91 | -0.14 | 0.03 | -0.06 |
| SOX2 | 2.35 | 0.91 | -0.40 | -0.07 | 0 | -0.02 |
| TBX6 | 0.51 | 0.07 | 0.08 | 0.32 | 0.03 | 0.94 |

**Day 4.5 LM**

| | | | | | | |
|---|---|---|---|---|---|---|
| VE | | 0.76 | 0.11 | 0.07 | 0.04 | 0.02 |
| BRA | 0.33 | 0.14 | -0.06 | 0.64 | 0.73 | -0.19 |
| CDX2 | 0.90 | 0.62 | -0.70 | -0.34 | 0.10 | -0.08 |
| SOX1 | 1.02 | 0.66 | 0.71 | -0.20 | 0.08 | -0.12 |
| SOX2 | 0.33 | 0.16 | 0.02 | 0.07 | 0.16 | 0.97 |
| TBX6 | 0.62 | 0.37 | -0.08 | 0.66 | -0.65 | 0 |

**Day 5 AN**

| | | | | | | |
|---|---|---|---|---|---|---|
| VE | | 0.80 | 0.17 | 0.03 | 0.01 | 0 |
| BRA | 0.32 | 0.02 | 0.01 | 0.07 | 0.58 | 0.81 |
| CDX2 | 1.14 | 0.05 | -0.05 | 0.94 | -0.30 | 0.13 |
| SOX1 | 2.37 | -0.03 | 1.00 | 0.04 | -0.04 | 0.01 |
| SOX2 | 4.56 | 1.00 | 0.03 | -0.06 | -0.03 | 0.01 |
| TBX6 | 0.79 | 0.05 | 0.02 | 0.32 | 0.75 | -0.57 |

**Day 5 CE**

| | | | | | | |
|---|---|---|---|---|---|---|
| VE | | 0.59 | 0.23 | 0.11 | 0.05 | 0.02 |
| BRA | 0.53 | 0.08 | -0.11 | 0.81 | -0.56 | 0.14 |
| CDX2 | 2.39 | 0.94 | -0.26 | -0.19 | -0.10 | -0.06 |
| SOX1 | 0.94 | 0.18 | 0.04 | 0.28 | 0.60 | 0.73 |
| SOX2 | 0.99 | 0.26 | 0.96 | 0.05 | -0.09 | -0.06 |
| TBX6 | 0.56 | 0.10 | -0.04 | 0.49 | 0.56 | -0.66 |

**Day 5 EM**

| | | | | | | |
|---|---|---|---|---|---|---|
| VE | | 0.78 | 0.14 | 0.05 | 0.02 | 0 |
| BRA | 0.39 | 0.14 | 0.23 | -0.10 | 0.96 | -0.07 |
| CDX2 | 1.11 | 0.43 | 0.79 | -0.32 | -0.29 | -0.07 |
| SOX1 | 0.87 | 0.17 | 0.27 | 0.93 | 0 | -0.18 |
| SOX2 | 0.28 | 0.07 | 0.12 | 0.14 | 0.05 | 0.98 |
| TBX6 | 1.96 | 0.87 | -0.49 | -0.02 | -0.01 | 0.01 |

**Day 5 PN**

| | | | | | | |
|---|---|---|---|---|---|---|
| VE | | 0.82 | 0.16 | 0.01 | 0.01 | 0 |
| BRA | 0.21 | 0.02 | 0.01 | 0.29 | 0.92 | -0.27 |
| CDX2 | 0.59 | 0.08 | 0.07 | 0.79 | -0.39 | -0.46 |
| SOX1 | 1.55 | -0.01 | 0.99 | -0.10 | 0.02 | -0.03 |
| SOX2 | 2.99 | 0.99 | 0 | -0.10 | 0.01 | -0.01 |
| TBX6 | 0.47 | 0.06 | 0.07 | 0.52 | 0.08 | 0.85 |

**Day 5 LM**

| | | | | | | |
|---|---|---|---|---|---|---|
| VE | | 0.81 | 0.11 | 0.04 | 0.03 | 0.01 |
| BRA | 0.26 | 0.10 | 0.09 | 0.84 | -0.50 | -0.17 |
| CDX2 | 0.75 | 0.50 | 0.85 | -0.16 | 0 | -0.05 |
| SOX1 | 0.79 | 0.76 | -0.50 | -0.25 | -0.34 | -0.06 |
| SOX2 | 0.26 | 0.14 | 0.01 | 0.17 | 0 | 0.97 |
| TBX6 | 0.50 | 0.38 | -0.16 | 0.42 | 0.80 | -0.13 |

Figure SI6: Mean and principal components for all clusters identified in the initial dataset for days 4 to 5 days. Each panel corresponds to the cluster and day specified at the top. The first column contains the mean value for the cells in the cluster. Each of the other columns corresponds to a principal component and it is coloured to ease the reading. The intensity of the colour corresponds to the absolute magnitude of the weight. The first row contains the amount of variation explained by that component (VE) and the rows below contain the weights of each protein in that component.

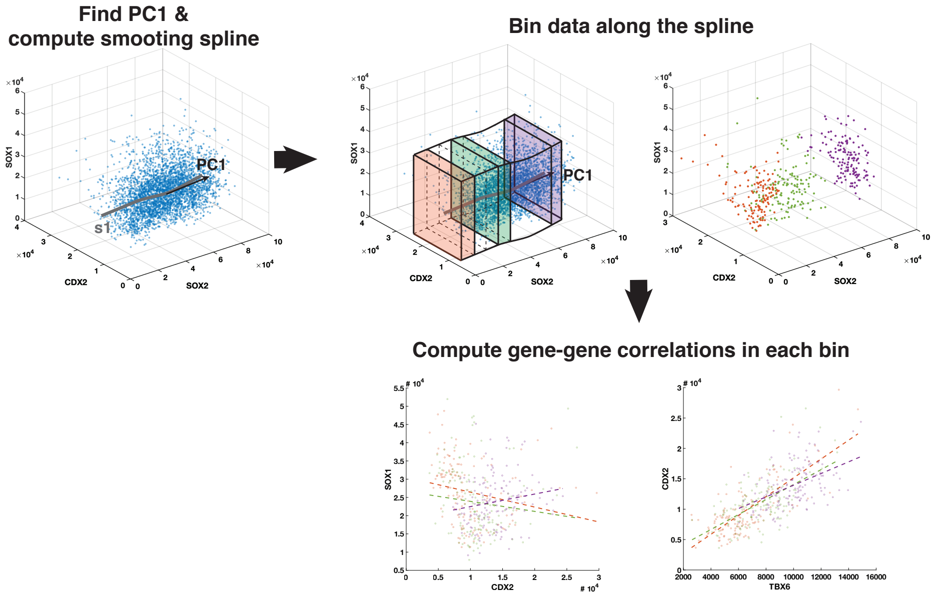

Figure SI7: Computation of local correlation. Each cluster is binned along a smoothing spline parametrised along its first principal component, which corresponds to the longest dimension of the ellipsoid defined by the points. The gene-gene correlation for the cells in each of the 25 bins is then computed. Since the clusters are normal, these correlations take similar values in all bins. For this example, we show the data corresponding to the cluster AN at D5. We have only coloured three bins out of the 25 for clarity.

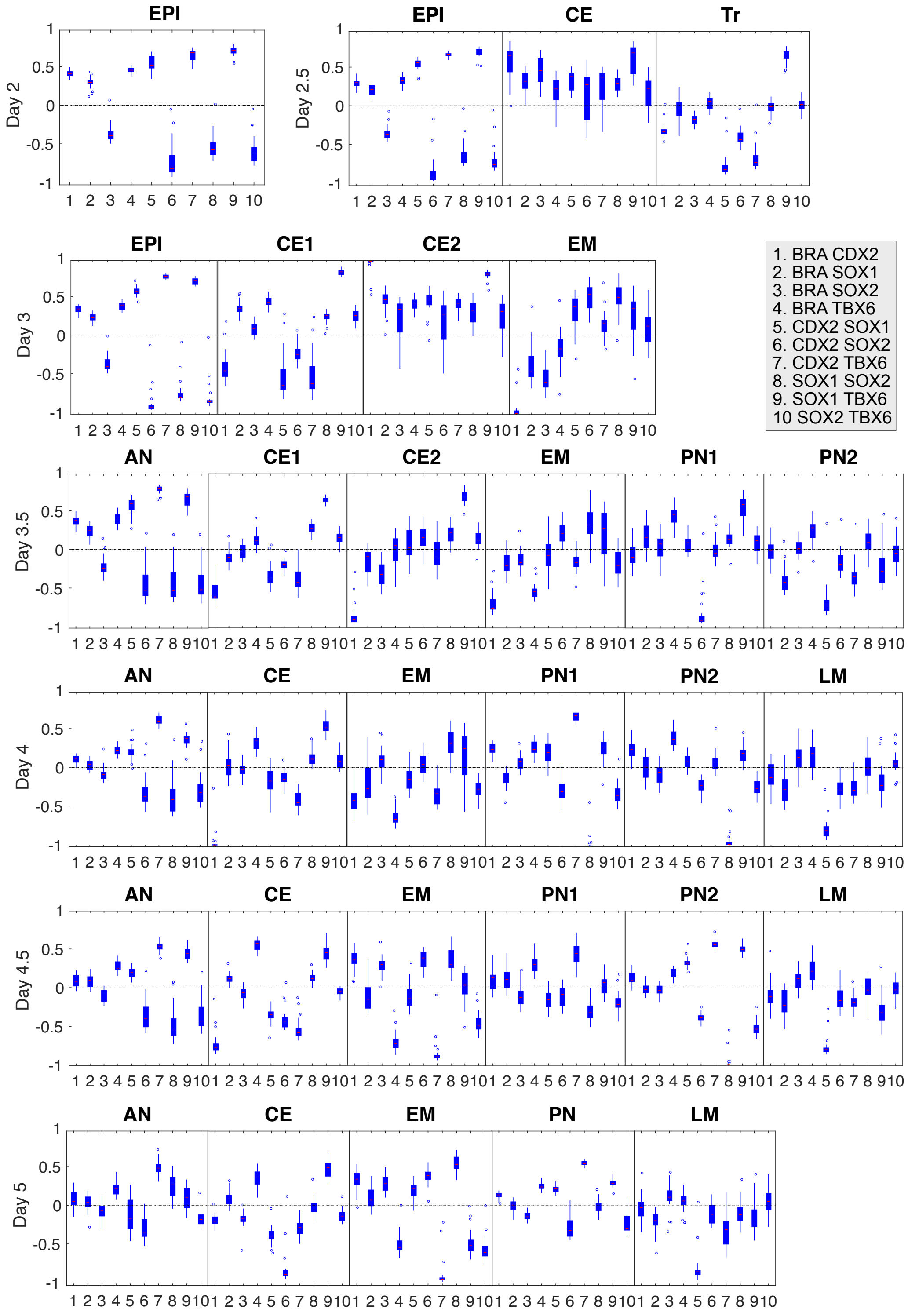

Figure SI8: Local gene-to-gene correlations along the first principal component for all clusters in the reference data set from the initial experimental series. The x-axis lists the gene pairs as detailed in the legend. The red segment is the median for the 25 gene-to-gene correlations. The blue box shows the range from the 25th to the 75th percentiles. The whiskers extend to the most extreme values that are not considered outliers. The outliers are marked with a circle.

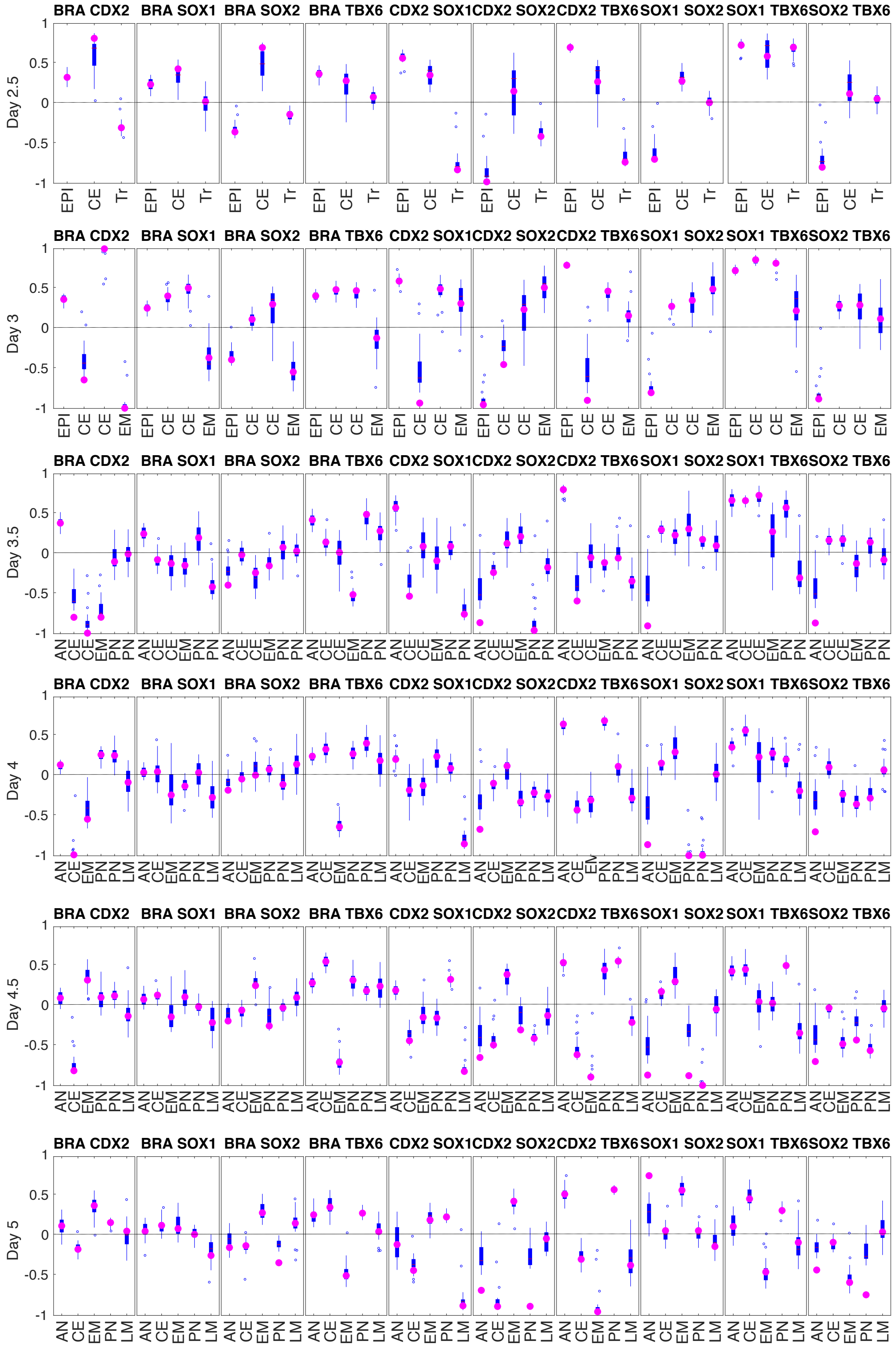

Figure SI9: Local gene-to-gene correlations along the first principal component for all clusters in the reference data set from the initial experimental series ordered by gene pair. Whisker plot as before. The pink circle denotes the value of the gene-to-gene correlation after projection on the orthogonal of the first principal component.

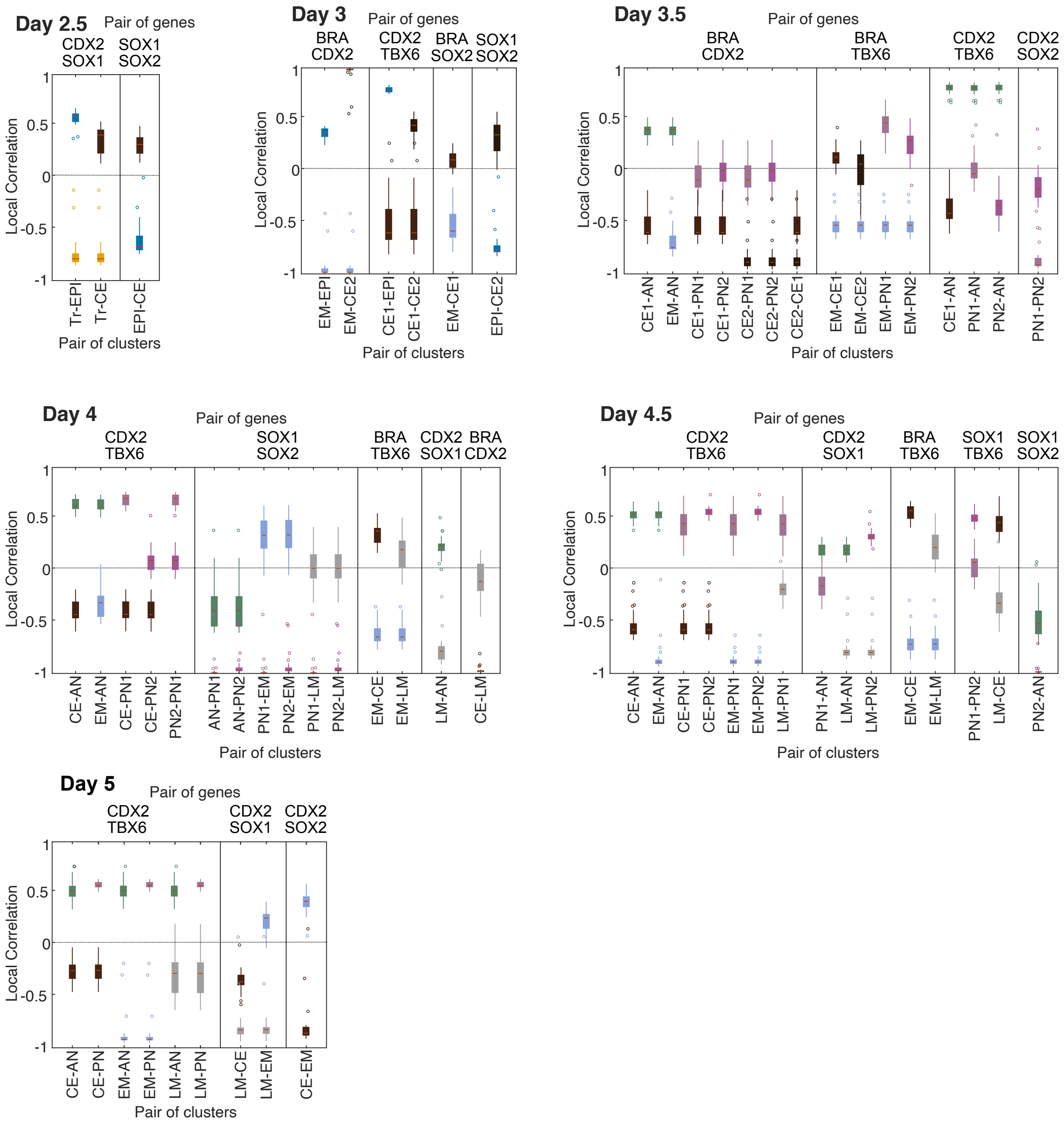

Figure SI10: Differences in the local correlations. An example of a very clear difference in local gene-to-gene correlation is shown for each pair of clusters. At day 5 there is no significant difference between the two neural clusters. The colours corresponds to cluster labels as in Fig. SI4 and the labels in the $x$ axis are ordered as the two boxes on top. Different subpanels correspond to different gene pairs as indicated on top.

### 3 Details of the mathematical model

#### 3.1 Choice of landscape

Of the elementary 3-attractor Parametrised Landscapes (PL) in [4] only the *binary choice* (Fig. SI11A) fits the observed qualitative data involving the possible attractors AN, EPI and CE. When there are three attractors, there are also two saddle points because we are considering compact systems. For all parameters, when it is present, the EPI attractor is the central one in the sense that the unstable manifolds of both saddles connect to it, whereas only one saddle connects to the other two attractors (which we therefore call peripheral). This central attractor can bifurcate by colliding with either saddle agreeing with the observation that EPI cells can transition to either AN or CE dependent on the signal. This implies the presence of the cusp point and fixes the diamond shape of the MS component (see section 1). The PL shown in Fig. SI11A is the simplest possible given these features. This cusp point is a dual cusp where two saddle points interact with an attractor that is between them (in this case EPI). The usual cusp discussed in Catastrophe Theory [5] is where two attractors interact with a saddle point.

For the transition between the possible attractors CE, M and PN we note that changing the signals causes cells to change the escape route from the initial state CE, guiding cells from CE to either the state PN or the state M. In addition, we note that we do not see any transitions between PN and M states where the saddle connecting them bifurcates with one of them. This rules out a possible PL containing a standard cusp, leaving only the possibility shown in Fig. SI11B, which we call the compactified elliptic umbilic, as the simplest candidate landscape. However, we only use part of this landscape as described below and we call this part the *binary flip*.

#### 3.2 Normal forms for the binary transitions

In order to obtain a model to perform simulations we chose normal forms for the two decisions: binary choice and binary flip. We used gradient dynamics defined by polynomial functions to define the landscape dynamics. In particular, we took subfamilies of the double cusp catastrophe ($x^4+y^4$), that contains both PLs mentioned earlier [9], to guarantee that the systems obtained were compact. That is, there were no trajectories escaping to infinity. In general, a Riemannian metric is necessary to incorporate all the dynamical properties of quasi-gradient Morse-Smale systems (see section 1) but we did not need it here, which kept the number of parameters of the model minimal.

For the first decision we used the parametrised family of potential functions

$$F_1(x, y; p) = x^4 + y^4 + y^3 - 4x^2y + y^2 - p_1\, x + p_2\, y. \tag{2}$$

The bifurcation locus for the family $F_1$ is as shown in Figure SI11A .

In this family the region with three attractors is the area inside the middle diamond shape. Here, the central attractor is always the same and it can bifurcate with any of the two saddles. For our model only the part of the family inside the central diamond and below it was relevant, because we considered the bifurcation of the middle attractor with either of the two saddles.

For the second decision we took the family of potential functions

$$F_2(x, y; p) = x^4 + y^4 + x^3 - 2xy^2 - x^2 + p_3\, x + p_4\, y. \tag{3}$$

The bifurcation locus for the family $F_2$ is shown in Figure SI11B.

In this family, the region with three attractors corresponds to the big central triangle, excluding the smaller one (blue curves with cusps) close to $(0, 0)$. Inside the small triangle a repeller in the middle of the three attractors appears as part of the landscape. The purple dotted lines denote the regions where the unstable manifold for one of the saddles flips, changing the middle attractor. For our model only the part inside the big central triangle on the left hand side and possibly further to the left is relevant because, we assume that neither PN nor M bifurcate.

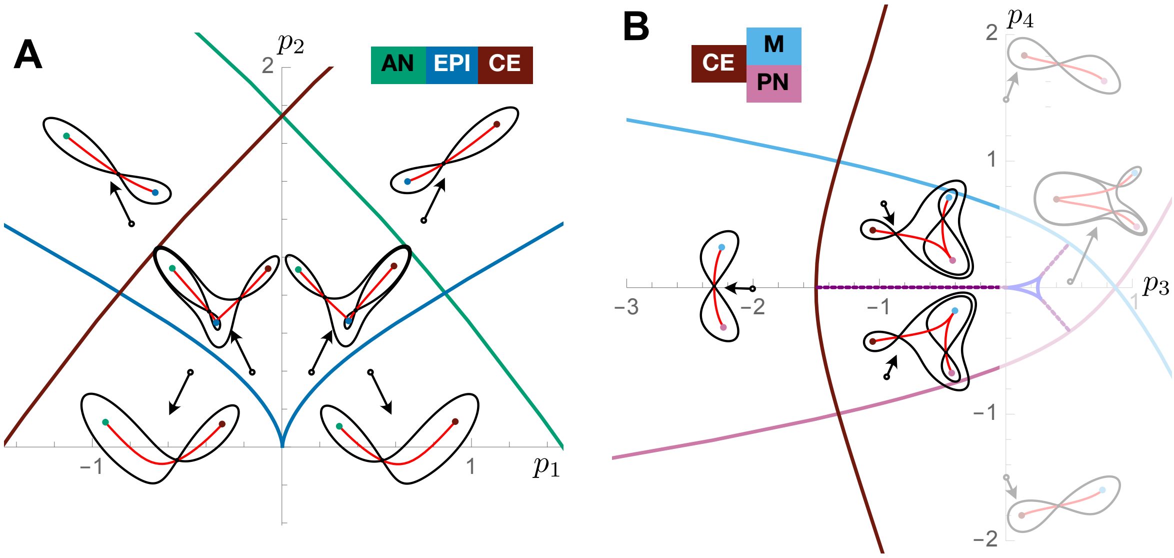

Figure SI11: A. Bifurcation curves for the family $F_1$ corresponding to the first decision. B. Bifurcation curves for the family $F_2$ corresponding to the second decision. Continuous curves correspond to bifurcation curves and they are coloured according to the attractor that bifurcates when crossing it. Purple dotted lines show the flip lines where the middle attractor changes. The landscape corresponding to a point on the purple line includes a saddle connection (the stable/ unstable manifold of one saddle contains the other one). Examples of landscapes in the different regions are shown. The black circle marks the exact parameter combination used to plot the landscape. Note that near the cusp in A the unstable manifold of the saddles (red curves) glue together smoothly at the middle attractor.

Each of the two landscapes contains an attractor assigned to CE, by which the two landscapes were "glued" together in order to build the global landscape.

#### 3.3 Defining the landscape model

The dynamical flow $L(x, y; p)$ for the global landscape was defined by

$$L(x, y, p) = -(1 - \chi(x))\, p_5\, \nabla F_1(x, y; p) - \chi(x)\, p_6\, \nabla F_2(x + 2, y + 1; p) \tag{4}$$

with

$$\begin{aligned}
F_1(x,y;p) &= x^4 + y^4 + y^3 - 4x^2y + y^2 + p_1\,x + p_2\,y && \text{binary choice landscape}\\
F_2(x,y;p) &= x^4 + y^4 - x^3 + 2xy^2 - x^2 - p_3\,x + p_4\,y && \text{binary flip landscape}\\
v &= (-2,-1) && \text{translation vector to match respective CE attractors}\\
\chi(x) &= \frac{tanh(10(x-0.5))+1}{2} && \text{gluing function.}
\end{aligned}$$

The global model depends on parameters $p_1, \ldots, p_6$ that are a function of the signals $s_1$, $s_2$.

The first step to join the two decisions was to translate one of the landscapes to place the two instances of the CE attractor at a similar position in the $(x, y)$ plane. We applied the translation $(-2, -1)$ to the second landscape $F_2$. The exact position of the attractors changes with different values of the parameters, although they remain in a fixed region for relatively small regions of the parameter space. With this translation the two instances of the common attractor land around $(1, 1)$ (Fig. SI12).

Since the global landscape should have only one attractor around $(1, 1)$, the second step was to establish a transition area so that the CE attractor from the first potential $F_1$ was lost in favour of the CE attractor from the second potential $F_2$ (in Fig. SI12 part of the level curve from $F_1$ is dashed meaning that it is deleted from the global model). That is, we glued the two landscapes using a sigmoid function to ensure a smooth transition between the two flows and we chose a vertical stripe around $x = 0.5$ for the transition to happen (vertical purple line in Fig. SI12)).

Finally, the dynamical flow $L(x, y; p)$ for the global landscape was defined by the linear combination of the two gradient systems corresponding to $F_1$ and $F_2$ (with $F_2$ translated) smoothly glued together with a sigmoid function $\chi$ as in equation (4). The parameters $p_5$ and $p_6$ control the magnitude of the gradient system for each of the two decisions and hence the velocity of the trajectories through that region.

| Parameter | Meaning |
|---|---|
| $p_1$ | topology of first decision |
| $p_2$ | topology of first decision |
| $p_3$ | topology of second decision (CE stability) |
| $p_4$ | topology of second decision (PN/M distribution) |
| $p_5$ | velocity of first decision |
| $p_6$ | velocity of second decision |

Table SI2: **Global model parameters** List of parameters for the global model. The first decision is the binary choice from EPI to either AN or CE. The second decision is the binary flip from CE to either PN or M.

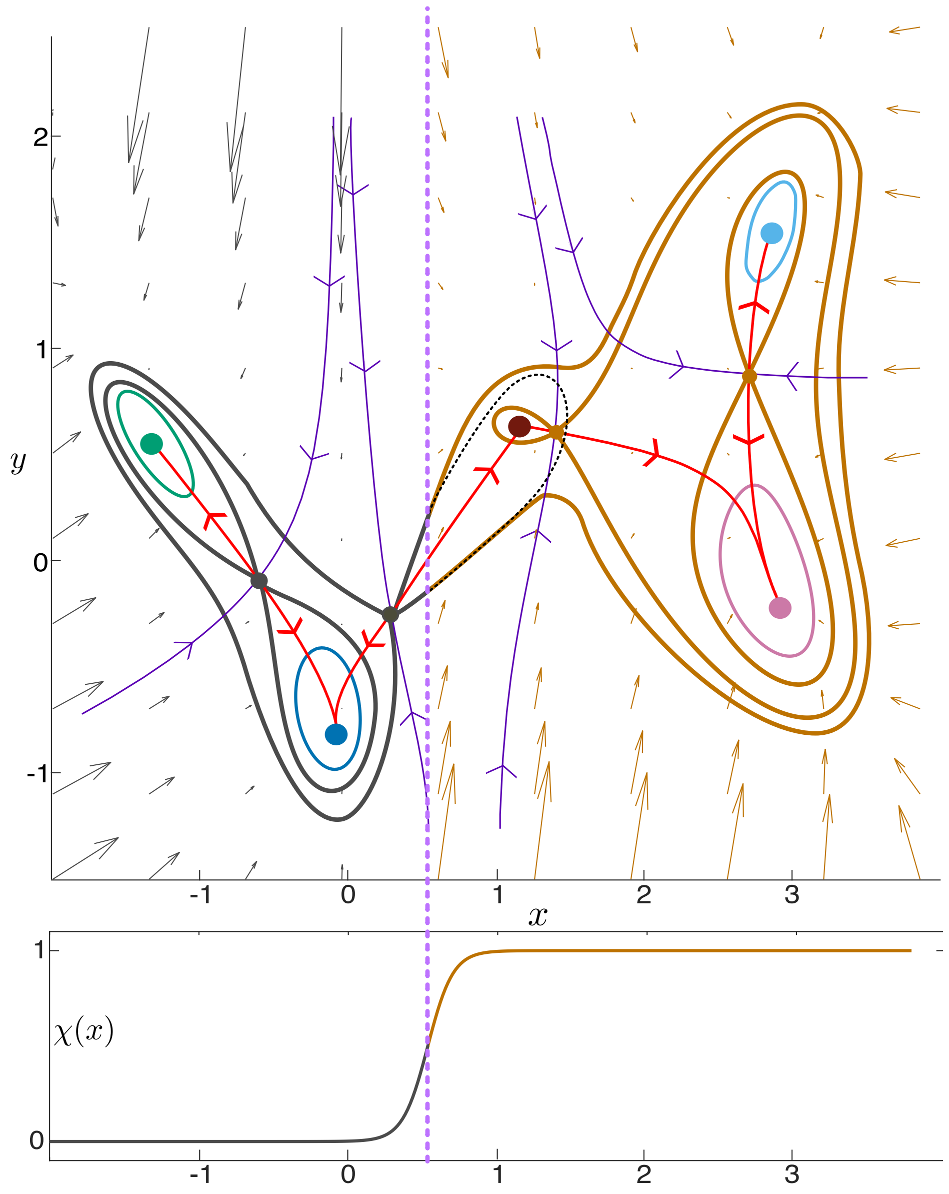

Figure SI12: Landscape flow as a result of gluing the gradient systems corresponding to the two decisions at the CE attractor.

#### 3.4 Parametrising the landscape model

The parameters $p = (p_1, \ldots p_6)$ are linear functions of the effective levels $S_1$ and $S_2$ of the two signals CHIR and FGF, respectively:

$$p(S_1, S_2) = (p_1(S_1, S_2), \ldots, p_6(S_1, S_2)) = w_0 + S_1\, w_1 + S_2\, w_2 = w_0 + (S_1\ \ S_2) \begin{pmatrix} w_1 \\ w_2 \end{pmatrix}$$

with $w_i \in \mathbb{R}^6$ for $i = 0, 1, 2$ parameter vectors to be estimated.

In the computation of the effective levels $S_1$ and $S_2$ at time $t$ is where the non-linearity comes into play. Let $(s_1(t), s_2(t))$ be the signal concentrations at time $t$.

| Parameter | Meaning |
|---|---|
| $w_0$ | Landscape parameters with No CHIR and PD |
| $w_1$ | Effect of CHIR in the landscape |
| $w_2$ | Effect of FGF in the landscape |

Table SI3: **Mapping parameters** List of parameters for the mapping from the signals (CHIR and FGF/PD) to the global model parameters.

- $s_1(t)$ is the concentration of CHIR at time $t$ where

$$s_1(t) = \begin{cases} 1 & \text{for saturated CHIR} \\ 0 & \text{for no CHIR} \end{cases}$$

- $s_2(t)$ is the concentration of FGF at time $t$ where

$$s_2(t) = \begin{cases} 1 & \text{for saturated FGF} \\ 0.9 & \text{for no FGF and no PD} \\ 0 & \text{for saturated PD} \end{cases}$$

  We define $s_2(t) = 0.9$ if there is no FGF and no PD because cells produce their own FGF and the data shows almost no difference between *CHIR 2-5* and *CHIR 2-5 FGF 2-5* conditions.

The idea is that the landscape changes instantaneously with a change of the signal level, but we observed that while after 24h of CHIR cells rapidly abandoned the CE state to become PN after CHIR removal, when the CHIR induction length was 48h they evolved almost exactly as in the case of continuous CHIR induction (CHIR 2-5). This prompted us to hypothesise that the effect of CHIR remains in the system for longer than the signal induction itself. Since for 36h of CHIR there seemed to be an intermediate delay on cells completely abandoning the CE attractor, we hypothesised that the longer the CHIR induction duration, the longer its effect remains in the system, and hence in the landscape. That is, $S_1$ changes value a certain time later than the change in $s_1$. We define a function $CT(t)$ that will quantify the amount of time of CHIR induction (5). It increases for concentrations above 0.5 and decreases for concentrations lower than 0.5. We assume that there is a threshold duration $\tau$ such that for CHIR induction shorter than $\tau$ no memory occurs and the landscape changes as we remove the signal $s_1$. For durations longer than $\tau$, the effect endures for a time proportional to the difference between $\tau$ and the CHIR induction duration. We introduced this threshold effect with a sharp sigmoid function that changes value around $\tau$. Therefore, the effective values $S_1$ and $S_2$ of the signals are computed as follows

$$CT(t) = max\left(\int_0^t 2s_1(u) - 1du, 0\right) \tag{5}$$

$$S_1(t) = min\left(1, s_1(t) + \frac{(tanh(10(CT(t) - \tau)) + 1)}{2}\right) \tag{6}$$

$$S_2(t) = s_2(t). \tag{7}$$

Figure SI13 shows the effect of the CHIR memory on the parameters for different durations of the signal.

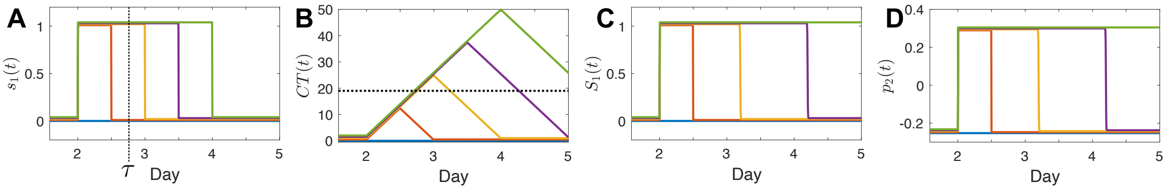

Figure SI13: A. Different profiles for $s_1(t)$ (different CHIR durations). Colours correspond to the signalling profiles throughout the panels. B. CHIR time for the different profiles in (A). C. Effective CHIR $S_1(t)$ for the different profiles in (A). D. Effect of the effective CHIR $S_1(t)$ in (C) on one parameter.

#### 3.5 Parametrised stochastic differential equation model

The form of the global landscape is determined by the effective signals, so with a change of variable we can rewrite the landscape flow as $L(x, y, S)$. Changes in the signalling regime change the particular form of $L$ in the family producing diverse differentiation patterns. The evolution of cells with time, given a signalling regime, is then modelled with a stochastic dynamical system [10]

$$(\dot{x}, \dot{y}) = L(x, y; S) + \sigma dW \tag{8}$$

where $dW$ is a two dimensional Wiener random process and $\sigma$ is a parameter controlling the amplitude of the noise perturbation. In total, the model depends on 20 parameters to be estimated:

$$(w_{0,1}, \ldots, w_{0,6}, w_{1,1}, \ldots, w_{1,6}, w_{2,1}, \ldots, w_{2,6}, \tau, \sigma). \tag{9}$$

#### 3.6 Simulation procedure

For each parameter vector as in (9) and for each experimental condition we simulated 500 trajectories on the landscape representing the transitioning of 500 cells. First, we assumed that the initial states of all the cells were equivalent and corresponded to the EPI state. It was also reasonable to incorporate some variability in the initial states, so we pre-initialised all the cells with the coordinated of the EPI attractor and let the

| Parameter | Meaning |
|---|---|
| $w_{0,1}$ | Topology of first decision with No CHIR and PD |
| $w_{0,2}$ | Topology of first decision with No CHIR and PD |
| $w_{0,3}$ | Topology of second decision with No CHIR and PD (CE stability) |
| $w_{0,4}$ | Topology of second decision with No CHIR and PD (PN/M distribution) |
| $w_{0,5}$ | Velocity of first decisions with No CHIR and PD |
| $w_{0,6}$ | Velocity of second decision with No CHIR and PD |
| $w_{1,1}$ | Effect of CHIR in the topology of the first decision |
| $w_{1,2}$ | Effect of CHIR in the topology of the first decision |
| $w_{1,3}$ | Effect of CHIR in the topology of the second decision (CE stability) |
| $w_{1,4}$ | Effect of CHIR in the topology of the second decision (PN/M distribution) |
| $w_{1,5}$ | Effect of CHIR in the velocity of the first decision |
| $w_{1,6}$ | Effect of CHIR in the velocity of the second decision |
| $w_{2,1}$ | Effect of FGF in the topology of the first decision |
| $w_{2,2}$ | Effect of FGF in the topology of the first decision |
| $w_{2,3}$ | Effect of FGF in the topology of the second decision (CE stability) |
| $w_{2,4}$ | Effect of FGF in the topology of the second decision (PN/M distribution) |
| $w_{2,5}$ | Effect of FGF in the velocity of the first decision |
| $w_{2,6}$ | Effect of FGF in the velocity of the second decision |
| $\tau$ | Memory to CHIR threshold |
| $\sigma$ | Noise amplitude |

Table SI4: **Parameters to be estimated** Complete list of parameters to be estimated by the ABC SMC method. Their meaning in the global landscape is detailed.

stochastic simulation run for 50 time steps, using a time step of $0.01h$. This allowed us to obtain an initial distribution of cells around the EPI attractor. We imposed that all the resulting points were in the EPI attractor by checking that all cells were closer to the attractor than the two saddle points separating the EPI attractor from the other attractors. If that was not the case, the parameter vector was rejected.

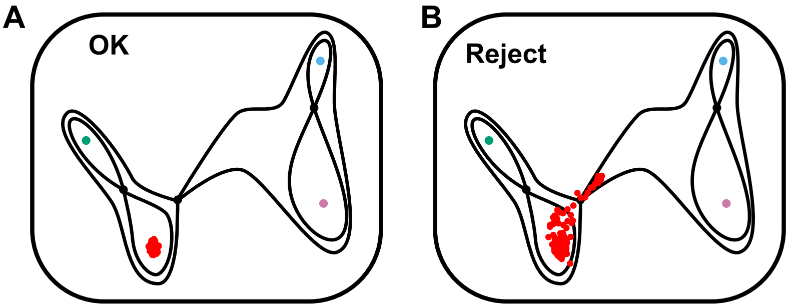

Figure SI14: Example of initial conditions. A. Acceptable initial set of points (in red). B. Initial set of points that would be rejected because there are points outside the EPI basin. For the particular set of parameters used in this example the CE attractor is bifurcated under the initial signal combination No CHIR with FGF.

By using the resulting points as initial conditions, we solved the system in Eq. 8 by using Euler-Maruyama [11, 12], obtaining 500 random walks on the landscape from $t = 0h$ to $t = 72h$, corresponding to days 2 to 5, using a time step $dt = 0.01h$ for each experimental condition.

We saved the coordinates for all simulated experimental conditions at the time points corresponding to the measured experimental points: from day 2 to day 5, every 12 hours, i.e. $t = 0, 12, \dots, 72$. To determine the corresponding cell identity we clustered the cells using a GMM in a similar fashion to the experimental data. We took all saved points, pooled them into a single dataset, and fitted a Gaussian model with 7 components corresponding to the seven cell identities in the experimental data (EPI, Tr, AN, CE, PN, M and UT). We computed the probability that a simulated cell could come from each component of the GMM and we assigned it to the most probable one. We initialised the mean values for the GMM fitting in points belonging to the approximate regions corresponding to the attractors in the model (or the transition it corresponded to). Since all the landscapes that were being considered belonged to the parametrised family of landscapes defined in (2) and (3), the specific position of each fixed point was restricted to a particular region of the $(x, y)$ plane. It is crucial here that the coefficients of $x^2$ and $y^2$ are constrained since otherwise the fixed points would have much more irrelevant flexibility. The specific values we used are listed in (10).

$$\begin{array}{llll} \text{EPI:}(0,-0.5) & \text{Tr:}(0.5,0) & \text{AN:}(-1.2,1) & \\ \text{CE:}(1,1) & \text{PN:}(3,0) & \text{M:}(3,2) & \text{UT:}(2,1) \end{array} \tag{10}$$

We let the matlab function `fitgmdist` compute the GMM. Matlab returns the GMM as an ordered list of Gaussian components with associated weights. If the distance between the computed means and the starting means was larger than a threshold, then the parameter vector was rejected. In this way we ensured that the order of the components of the computed GMM corresponded to the order we fixed and that there were points around all attractors. We included a distribution for UT because otherwise the cells transitioning from CE to PN or M were classified as belonging to one of the three attractors randomly. An example of the result of such clustering is shown in Fig. SI15. The method did not always work exactly as expected, as discussed in section 5 (see figures SI24 and SI25).

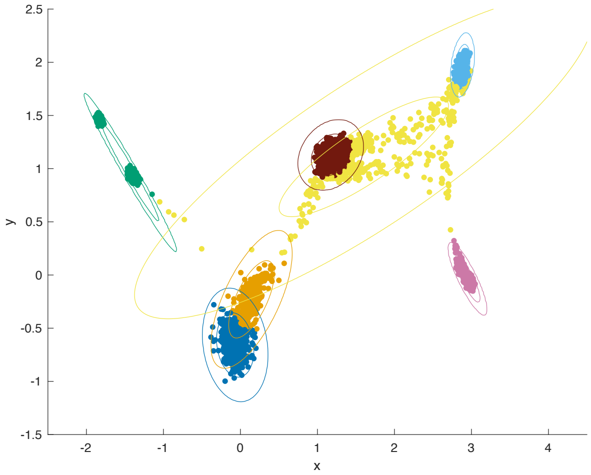

Figure SI15: Example of simulated cells with assigned fates using a GMM clustering procedure. The cells are pooled from all time points and experimental conditions and correspond to one specific parameter vector. The cluster corresponding to AN is defined by the two groups of green points, so the corresponding normal component is long in the direction that expands them. There are two groups of points corresponding to AN because the AN attractor significantly moves depending on the signalling conditions. The normal components corresponding to EPI (dark blue) and Tr (orange) are superposed because the Tr cluster corresponds to cells abandoning the EPI attractor. This defines an interface between the two clusters corresponding to cells with equal probability to come from either component. A similar interface could be defined by two close attractors if the noise amplitude is large enough. The component corresponding to UT defines a large distribution that captures all cells that are not clearly assigned to any of the other clusters.

### 4 Fitting algorithm

As described above, the landscape model proposed depends on a total of 20 parameters. The underlying model depends upon 6 parameters: 2 time constants used as velocities through the landscape and 4 that determine the shape of the landscape, each of which is a linear function of the signals. Of the 20 parameters, 18 are used to parametrise the dependence of the model on the signals while 2 are used to specify the time threshold for the CHIR memory and the noise amplitude. In order to fit this model and validate the results, we used a training data set which contained 7 experimental conditions (a total of 343 independent proportions), and we used a validation set which contained 4 experimental conditions (a total of 196 independent proportions), which can be found in Table T2.

In order to find a set of parameters with which our model could reproduce the experimental training data set, we followed the approach developed in [13] and took advantage of the approximate Bayesian computation (ABC) framework. In this approach, due to the high dimensional nature of the parameter space of the model and data, the sequential Monte Carlo ABC (ABC SMC) algorithm of [19] is implemented, with a multivariate normal perturbation kernel with optimal covariance matrix (OLCM) [20], which allowed us to parallelise the fitting while efficiently exploring the parameter space.

Starting from some proposed prior distribution for the parameters, the ABC SMC sampler methodology approximates a posterior distribution by sequentially generating a set of probability distributions

$$\{\pi_s\}_{0\leq s\leq T} = \{\pi(\boldsymbol{\theta} \mid d(\boldsymbol{X}(\boldsymbol{\theta}), \boldsymbol{X}_0) \leq \varepsilon_s)\}_{0\leq s\leq T},$$

where $\boldsymbol{\theta}$ is a parameter vector, $\boldsymbol{X}(\boldsymbol{\theta})$ is the simulated data with the corresponding parameter vector, $\boldsymbol{X}_0$ is the training data, $d$ is a distance between the simulated and experimental data, that we will define below, and $\{\varepsilon_s\}_{\{0\leq s\leq T\}}$ is a decreasing sequence of positive numbers, also to be defined. To produce these probability distributions, the algorithm starts by sampling $N$ parameter values $\boldsymbol{\theta}^*$ from a prior distribution, $\pi(\cdot)$, and which satisfy $d(\boldsymbol{X}(\boldsymbol{\theta}^*), \boldsymbol{X}_0) \leq \varepsilon_1$, in this context called particles. To each of the initial accepted particles $\{\boldsymbol{\theta}_1^{(i)}\}_{i=1}^N$, it assigns a set of equal weights $\{\omega_1^{(i)} = 1/N\}_{i=1}^N$, approximating the first probability distribution $\pi_1$.

Then, the algorithm proceeds in a sequential manner. At each step $s$, a set of $N$ particles $\{\boldsymbol{\theta}_s^{(i)}\}_{i=1}^N$ is generated by sampling particles,$\boldsymbol{\theta}^{**}$, from the discrete distribution with support the $N$ particles generated at the previous step, $\{\boldsymbol{\theta}_{s-1}^{(i)}\}_{i=1}^N$, and probabilities given by the weights $\{\omega_{s-1}^{(i)}\}_{i=1}^N$. Then, the sample is perturbed using the Markov kernel function $K_s(\boldsymbol{\theta}^*, \boldsymbol{\theta}^{**})$. These steps are repeated until $N$ particles $\boldsymbol{\theta}^*$ that satisfy $\pi(\boldsymbol{\theta}^*) > 0$ and $d(\boldsymbol{X}(\boldsymbol{\theta}^*), \boldsymbol{X}_0) \leq \varepsilon_s$ are found.

Therefore, for the implementation of this algorithm we needed to:

1. Determine the range in which the parameters are defined and define the prior distributions for the parameters to be fitted.

2. Study constraints on the parameters imposed by the data.

3. Choose a suitable number of particles $N$ that will approximate the distribution $\{\pi_s\}_{0\leq s\leq T}$ and each step $s$ of the algorithm.

4. Define a distance function $d$ to compare the simulations and the data and define a decreasing sequence of thresholds $\{\varepsilon_s\}_{\{0\leq s\leq T\}}$ that will determine the maximum distance between the experimental data and the simulated data at each step $s$ of the algorithm.

5. Define a perturbation kernel $\{K_s(\cdot,\cdot)\}_{s=1}^{T}$ that will set the region in the parameter space to be explored at each step. As mentioned before, we use a multivariate normal perturbation kernel with optimal covariance matrix (OLCM) [20, 13].

#### 4.1 Priors for the parameters

In order to implement this algorithm, we chose uniform uninformative parameters prior distributions for the time constants ($w_{i,5}, w_{i,6}$), memory threshold ($\tau$) and noise amplitude ($\sigma$) parameters. The data that we had informed the shape of the landscape for certain signal combinations, that is, it informed the parameters $p_i(S_1, S_2)$ for $i = 1, \ldots 4$. We define

$$p_i^{jk} = p_i(S_1, S_2) = w_{0,i} + S_1 \cdot w_{1,i} + S_2 \cdot w_{2,i} \text{ for } i = 1, \ldots 4 \text{ with } j = S_1 \text{ and } k = S_2.$$

We chose the priors for the landscape parameters corresponding to 3 experimental conditions (No Chir + FGF, Chir + FGF, Chir + PD), that is, $p_i^{01}, p_i^{11}, p_i^{10}$. Since our signals define an affine space of dimension 2, these three experimental conditions were sufficient to define a basis of the landscape space. From these 3 vectors one can compute the $w_i$'s and therefore any element from the family (see Fig. 4A in main text). We used catastrophe theory (Figures SI11) to choose adequate priors. We show the priors together with the relevant bifurcation locus on figures SI33 to SI35. For example, when considering the second decision we always assumed $p_3^{ij} > 0$ because we wanted the CE attractor to be the highest one in the potential, and we chose the range for $p_4^{ij}$ depending on the PN:M balance. The no CHIR case favours PN, so we assumed $p_4^{01} > 0$, the CHIR + FGF regime favours M, so we chose $p_4^{11} < 0$ and the CHIR + PD seems to distribute fates among both of them, so we chose a prior centred around 0. The priors can be found in table SI5.

When sampling the time constants we imposed that all $p_5^{j,k}$ and $p_6^{j,k}$ took positive values.

#### 4.2 Choice of distance function

Given a particle, we generated the simulated data as explained in Section 3.6 for the experimental conditions included in the training data ($TD$). We obtained a proportion $p_{\{t,e,f\}}^{sim}(\boldsymbol{\theta})$ of simulated cells in each of the seven states $f$ (EPI, Tr, AN, CE, PN, M and UT) for each timepoint $t$ and experimental condition $e$, obtaining the $7 \times 7 \times 7$ matrix of simulated proportions $\boldsymbol{X}^{TD}(\boldsymbol{\theta}) = (p_{\{e,f,t\}}^{sim}(\boldsymbol{\theta}))$, that we then compared to the experimental data, $\boldsymbol{X}_0^{TD} = (p_{\{e,f,t\}}(\boldsymbol{\theta}))$ in the following way:

$$d(\boldsymbol{X}^{TD}(\boldsymbol{\theta}), \boldsymbol{X}_0^{TD}) = \sum_{e\in TD} \sum_{f=1}^{7} \sum_{t=1}^{7} \left| p_{e,f,t}^{sim}(\boldsymbol{\theta}) - p_{e,f,t} \right| .$$

In running the algorithm we considered the proportions as values over a total of 1 instead of percentages over 100. The thresholds were chosen accordingly.

Table SI5: Tables of priors for the parameters.

| Parameter | Prior | Parameters | Prior |
|---|---|---|---|
| $p_1^{01}$ | $\mathcal{U}([-0.5, 0])$ | $w_{1,5}$ | $\mathcal{U}([-1, 1])$ |
| $p_2^{01}$ | $\mathcal{U}([0.5, 1.5])$ | $w_{1,6}$ | $\mathcal{U}([-1, 1])$ |
| $p_3^{01}$ | $\mathcal{U}([-3, -0.5])$ | $p_1^{10}$ | $\mathcal{U}([-6, 0])$ |
| $p_4^{01}$ | $\mathcal{U}([0, 1])$ | $p_2^{10}$ | $\mathcal{U}([-12, 0])$ |
| $w_{0,5}$ | $\mathcal{U}([0, 1])$ | $p_3^{10}$ | $-\Gamma(9.8, 0.14)$ |
| $w_{0,6}$ | $\mathcal{U}([0, 1])$ | $p_4^{10}$ | $N(0, 0.2)$ |
| $p_1^{11}$ | $\mathcal{U}([0, 1])$ | $w_{2,5}$ | $\mathcal{U}([-1, 1])$ |
| $p_2^{11}$ | $\mathcal{U}([0, 1])$ | $w_{2,6}$ | $\mathcal{U}([-1, 1])$ |
| $p_3^{11}$ | $\mathcal{U}([-2, 0])$ | $\tau$ | $\mathcal{U}([12, 36])$ |
| $p_4^{11}$ | $\mathcal{U}([-1, 0])$ | $\sigma$ | $\mathcal{U}([0, 0.1])$ |

##### 4.3 Choice of thresholds

For each step $s = 1, 2, \ldots$ of the algorithm, a threshold $\varepsilon_s$ needs to be defined, which should be smaller than the previous threshold and which determines the accepted distance between the simulated data and the experimental data.

For $s = 1$, we chose an $\varepsilon_s$ which was sufficiently challenging to ensure a relatively small set of accepted parameter vectors, but which was not so small that the initial finding of acceptable parameter vectors took an unreasonable amount of computational time. After the initial step, following a common approach [21, 18], the threshold at time $s$ was obtained as the $\alpha$-quantile of the distances obtained at the previous step $s-1$. In our case, we chose $\alpha = 0.15$. The resulting thresholds are shown in Fig. SI16, SI36 and SI51.

### 5 Fitting results

Our goal was to find parameter vectors for our model that were able to reproduce the experimental data within a reasonable accuracy with respect to the expected experimental variation. We present the results of the fitting algorithm in this section (and the following ones) without regarding the weights obtained by the algorithm. We were not seeking a posterior distribution for the real parameters but only to prove the existence of good enough parameter vectors.

We ran the ABC SMC algorithm for 13 iterations changing the acceptance threshold at each iteration as indicated in figure SI16. We stopped the algorithm at this point because the 14th threshold would have been very close to the 13th one and the parameter distributions looked similar in iterations 12 and 13 (Fig. SI31).

We present the results showing first the distributions of cell fate proportions (for the 7 fates and 6 time points) for the 10000 parameter vectors accepted in the last

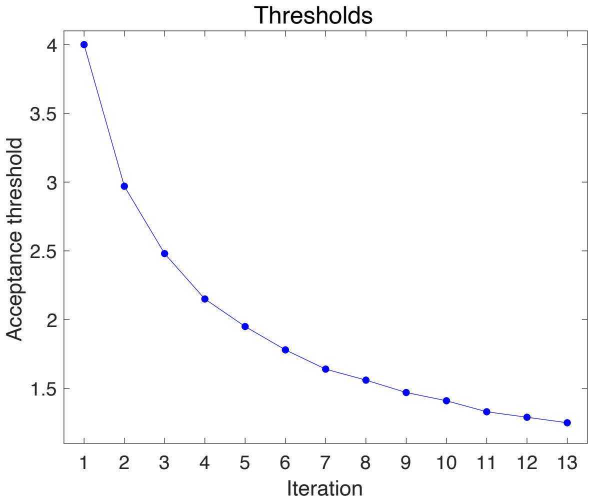

Figure SI16: Evolution of acceptance threshold in the 13 iterations of the initial fitting algorithm.

iteration of the algorithm. We did not include day 2 since the distribution of fates corresponds to the initial condition with all cells in the EPI state. We show first the results for the experimental regimes used as training and then the results for the validation regimes. The fate proportion arising from the experimental data used as training data is indicated with a vertical dotted line. Note that they are mostly normal distributions. We show the corresponding variation coefficients in tables SI6 and SI7.

#### 5.1 Fate distributions

##### 5.1.1 Training Data

In figures SI17 to SI23 we show the distributions of the proportions of simulated cells assigned to each fate for the all the parameter vectors accepted in the last iteration of the initial fitting algorithm. Most of these distributions are unimodal except for the distributions of M and UT in the CHIR 2-3 regime and the distribution of AN in the CH 2-5 FGF 0-3 regime. These fates correspond to the largest coefficients of variation in table SI6.

The bimodality of the fates M and UT under CHIR 2-3 conditions is due to a technical issue with the Gaussian mixture model fitted to the simulated cells. For some of the parameter vectors, the cells around the M attractor under No CHIR and No FGF regime were classified as transitioning cells, while the M cluster laid above, corresponding to the position of the attractor under CHIR induction conditions as shown in figure SI24D. The distribution of the sum of both M and UT proportions show a unimodal distribution with mean close to the experimental proportion from the initial experimental series.

The bimodality of the fates AN and UT under CHIR 2-5 FGF 0-3 conditions is due to a technical issue with the Gaussian mixture model fitted to the simulated cells. For some of the parameter vectors, the cells around the AN attractor under CHIR and PD regime were classified as transitioning cells, while the AN cluster laid below, corresponding to the position of the attractor under No CHIR induction conditions as shown figure SI25C. The distribution of the sum of both AN and UT proportions

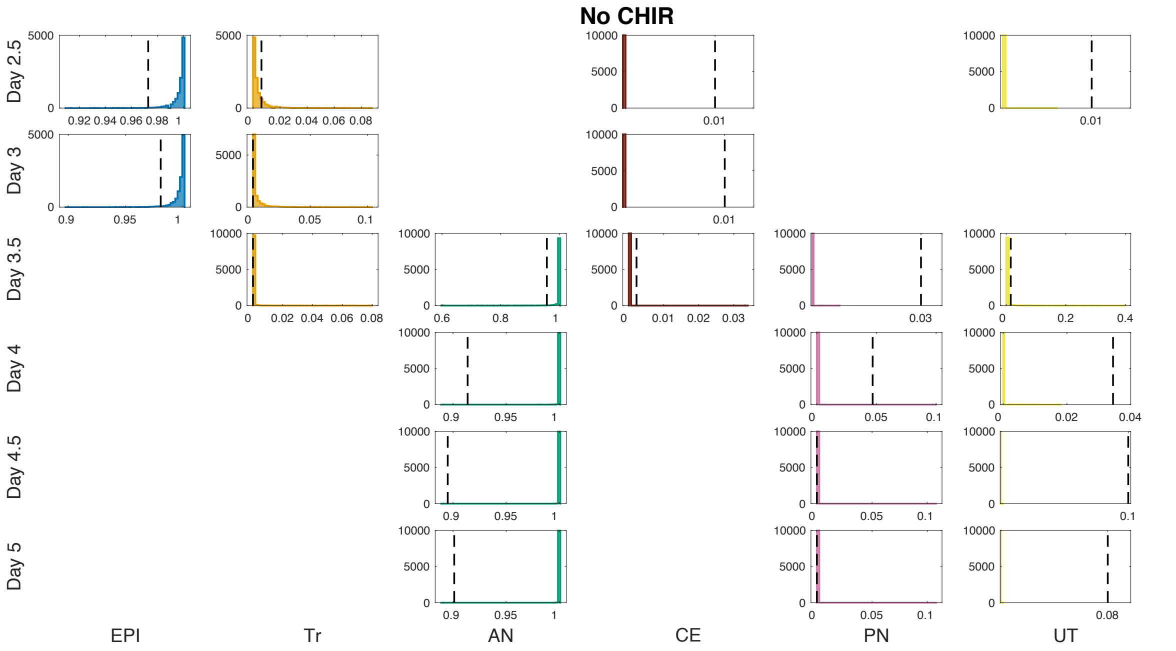

Figure SI17: Proportion distributions for No CHIR regime using the 10000 parameter vectors accepted in the last iteration of the initial fitting algorithm. The vertical dotted lines indicate the proportions from the initial experimental series used as training data. The missing panels correspond to populations with no assigned simulated cells and experimental proportion equal to zero. Note that each panel has different vertical and horizontal ranges.

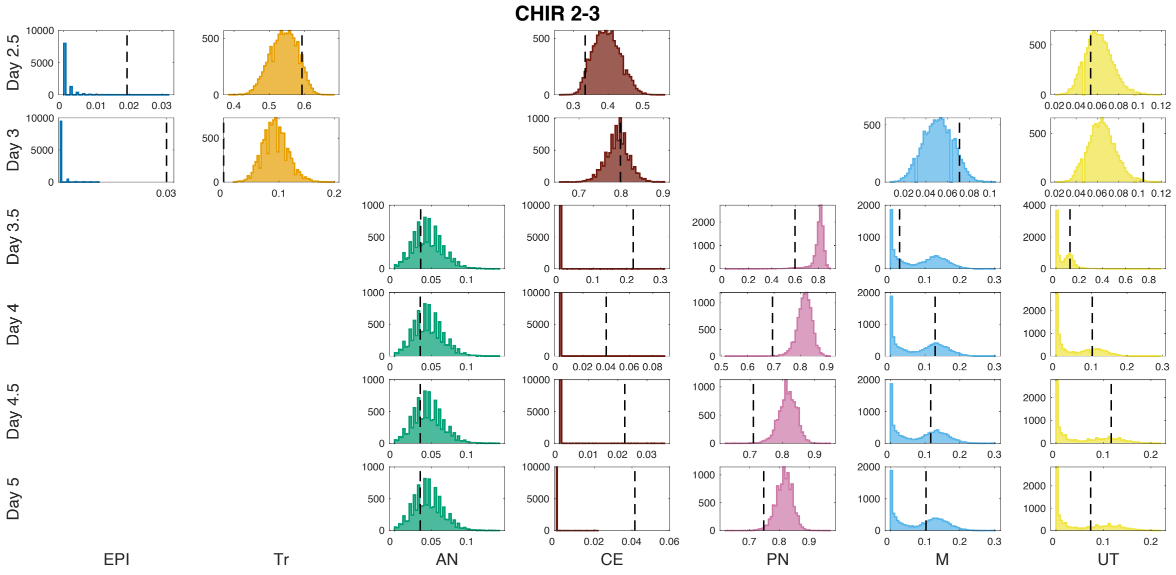

Figure SI18: Proportion distributions for CHIR 2-3 regime simulations in the last iteration of the initial fitting algorithm used as training data. The vertical dotted lines indicate the proportion from the initial experimental series.

show a unimodal distribution.

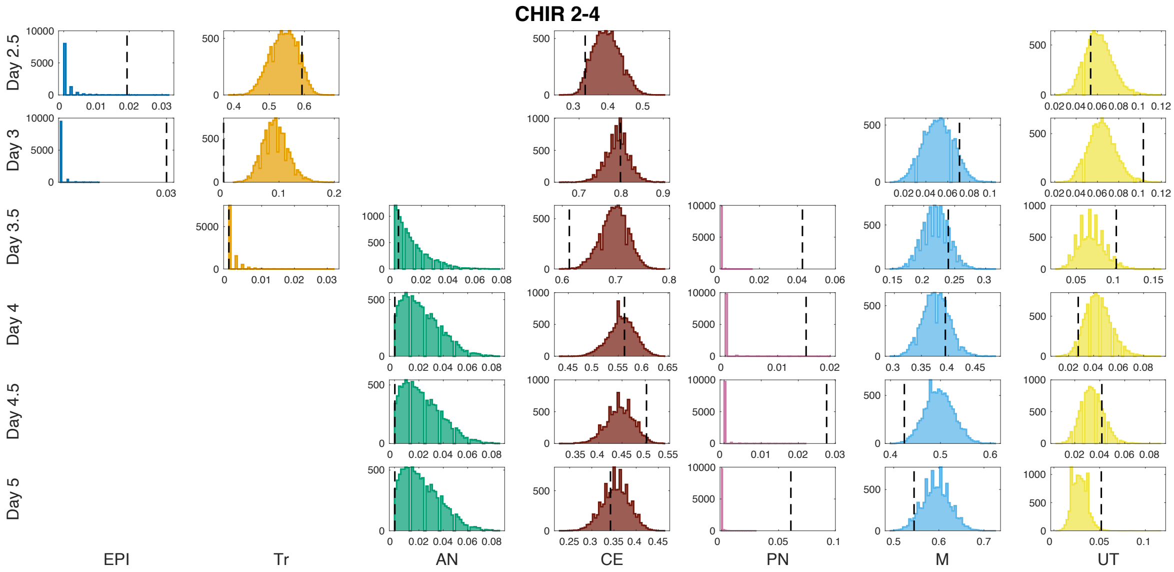

Figure SI19: Proportion distributions for CHIR 2-4 regime simulations in the last iteration of the initial fitting algorithm used as training data. The vertical dotted lines indicate the proportion from the initial experimental series.

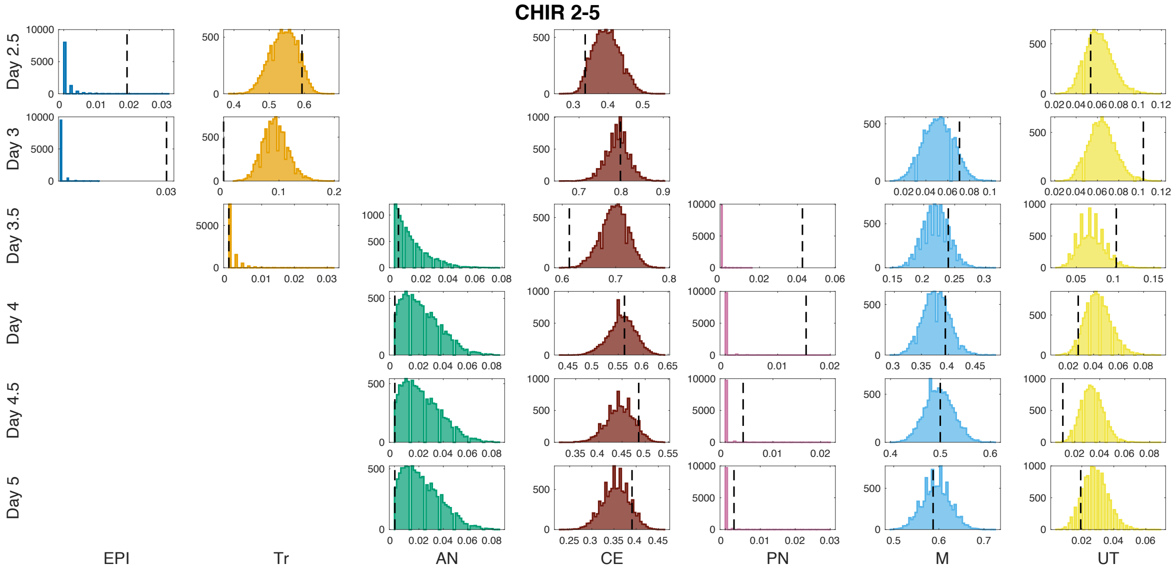

Figure SI20: Proportion distributions for CHIR 2-5 regime simulations in the last iteration of the initial fitting algorithm used as training data. The vertical dotted lines indicate the proportion from the initial experimental series.

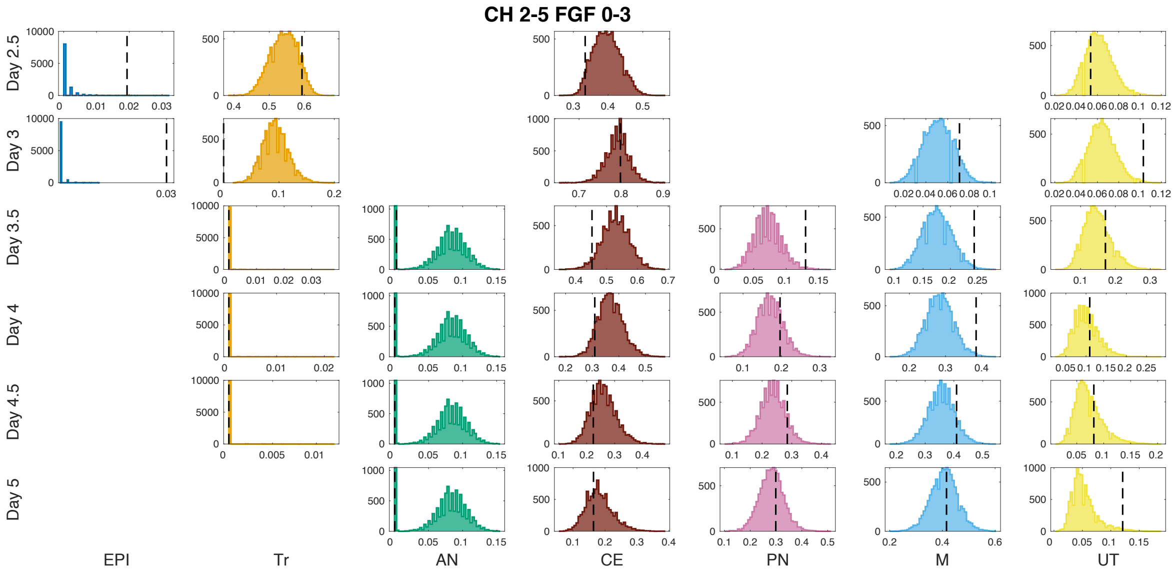

Figure SI21: Proportion distributions for CHIR 2-5 FGF 0-3 regime simulations in the last iteration of the initial fitting algorithm used as training data. The vertical dotted lines indicate the proportion from the initial experimental series.

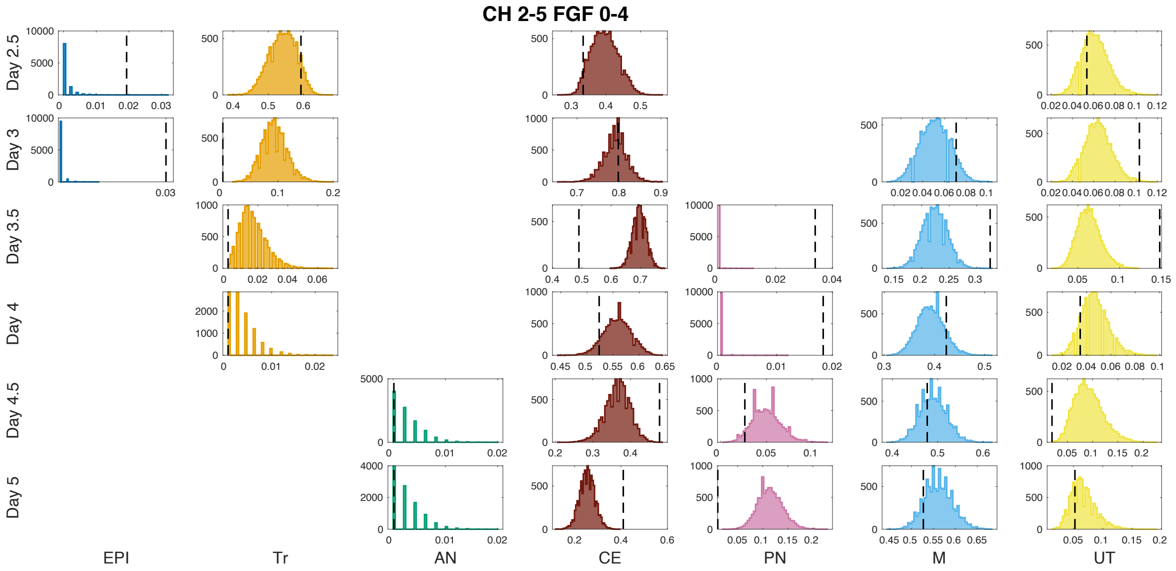

Figure SI22: Proportion distributions for CHIR 2-5 FGF 0-4 regime simulations in the last iteration of the initial fitting algorithm used as training data. The vertical dotted lines indicate the proportion from the initial experimental series.

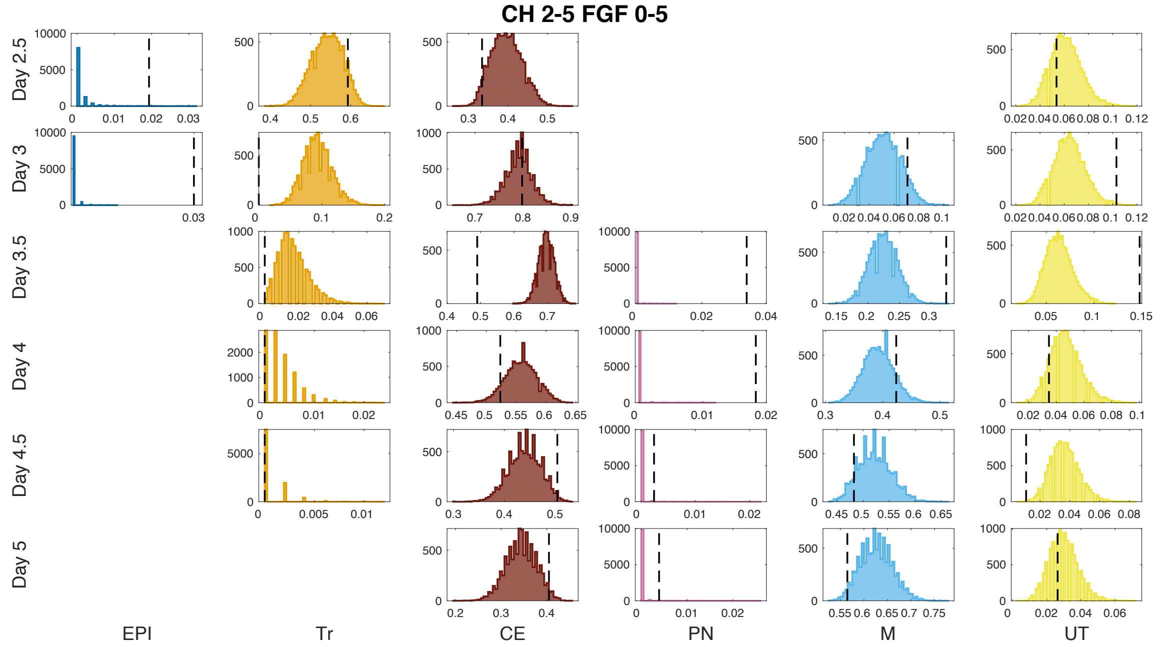

Figure SI23: Proportion distributions for CHIR 2-5 FGF 0-5 regime simulations in the last iteration of the initial fitting algorithm used as training data. The vertical dotted lines indicate the proportion from the initial experimental series.

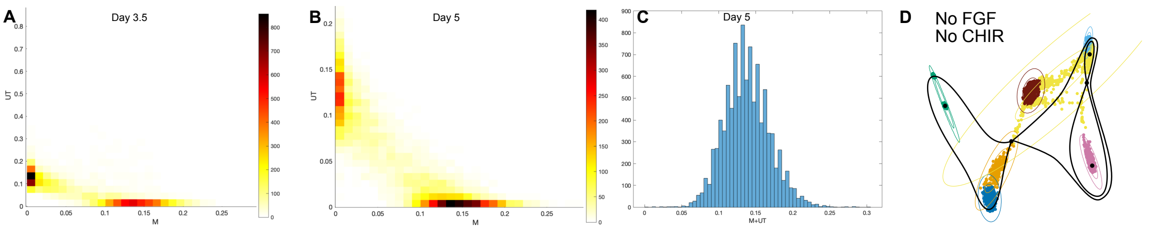

Figure SI24: Analysis of fate bimodality for M and UT under CHIR 2-3 conditions. A. Relative proportion distributions for M and UT under CHIR 2-3 regime simulations at day 3.5. B. Relative proportion distributions for M and UT under CHIR 2-3 regime simulations at day 5. C. Distribution of the sum of M and UT at day 5. D. Landscape for No FGF and No CHIR conditions defined by a parameter vector giving a large UT proportion at day 5 under CHIR 2-3 regime. The coloured level curves show the different components of the Gaussian mixture model fitted to the simulated data corresponding to the same parameter vector. Coloured points correspond to simulated cells for all training experimental conditions. Colours correspond to cell identities as in Figure 3B.

| Condition | Day | EPI | Tr | AN | CE | PN | M | UT |
|---|---|---|---|---|---|---|---|---|
| No CHIR | 2.5 | 0.01 | 0 | 0 | 0 | 0 | 0 | 0 |
| | 3 | 0.01 | 0 | 0 | 0 | 0 | 0 | 0 |
| | 3.5 | 0 | 0 | 0.02 | 0 | 0 | 0 | 0 |
| | 4 | 0 | 0 | 0 | 0 | 0 | 0 | 0 |
| | 4.5 | 0 | 0 | 0 | 0 | 0 | 0 | 0 |
| | 5 | 0 | 0 | 0 | 0 | 0 | 0 | 0 |
| CHIR 2-3 | 2.5 | 0 | 0.07 | 0 | 0.1 | 0 | 0 | 0.21 |
| | 3 | 0 | 0.25 | 0 | 0.03 | 0 | 0.27 | 0.2 |
| | 3.5 | 0 | 0 | 0 | 0 | 0.08 | 0.78 | 1.1 |
| | 4 | 0 | 0 | 0 | 0 | 0.04 | 0.78 | 1 |
| | 4.5 | 0 | 0 | 0 | 0 | 0.03 | 0.78 | 1 |
| | 5 | 0 | 0 | 0 | 0 | 0.03 | 0.78 | 1.01 |
| CHIR 2-4 | 2.5 | 0 | 0.07 | 0 | 0.1 | 0 | 0 | 0.21 |
| | 3 | 0 | 0.25 | 0 | 0.03 | 0 | 0.27 | 0.2 |
| | 3.5 | 0 | 0 | 0 | 0.04 | 0 | 0.1 | 0.23 |
| | 4 | 0 | 0 | 0 | 0.05 | 0 | 0.06 | 0 |
| | 4.5 | 0 | 0 | 0 | 0.07 | 0 | 0.06 | 0 |
| | 5 | 0 | 0 | 0 | 0.09 | 0 | 0.05 | 0 |
| CHIR2-5 | 2.5 | 0 | 0.07 | 0 | 0.1 | 0 | 0 | 0.21 |
| | 3 | 0 | 0.25 | 0 | 0.03 | 0 | 0.27 | 0.2 |
| | 3.5 | 0 | 0 | 0 | 0.04 | 0 | 0.1 | 0.23 |
| | 4 | 0 | 0 | 0 | 0.05 | 0 | 0.06 | 0 |
| | 4.5 | 0 | 0 | 0 | 0.07 | 0 | 0.06 | 0 |
| | 5 | 0 | 0 | 0 | 0.08 | 0 | 0.05 | 0 |
| CH 2-5 FGF 2-3 | 2.5 | 0 | 0.07 | 0 | 0.1 | 0 | 0 | 0.21 |
| | 3 | 0 | 0.25 | 0 | 0.03 | 0 | 0.27 | 0.2 |
| | 3.5 | 0 | 0 | 0.43 | 0.08 | 0.29 | 0.15 | 0.26 |
| | 4 | 0 | 0 | 0.43 | 0.13 | 0.2 | 0.13 | 0.3 |
| | 4.5 | 0 | 0 | 0.43 | 0.18 | 0.18 | 0.13 | 0.35 |
| | 5 | 0 | 0 | 0.43 | 0.23 | 0.17 | 0.13 | 0.43 |
| CH 2-5 FGF 2-4 | 2.5 | 0 | 0.07 | 0 | 0.1 | 0 | 0 | 0.21 |
| | 3 | 0 | 0.25 | 0 | 0.03 | 0 | 0.27 | 0.2 |
| | 3.5 | 0 | 0 | 0 | 0.03 | 0 | 0.1 | 0.22 |
| | 4 | 0 | 0 | 0 | 0.05 | 0 | 0.07 | 0 |
| | 4.5 | 0 | 0 | 0 | 0.09 | 0 | 0.06 | 0.31 |
| | 5 | 0 | 0 | 0 | 0.13 | 0.22 | 0.06 | 0.32 |
| CH 2-5 FGF 2-5 | 2.5 | 0 | 0.07 | 0 | 0.1 | 0 | 0 | 0.21 |
| | 3 | 0 | 0.25 | 0 | 0.03 | 0 | 0.27 | 0.2 |
| | 3.5 | 0 | 0 | 0 | 0.03 | 0 | 0.1 | 0.22 |
| | 4 | 0 | 0 | 0 | 0.05 | 0 | 0.07 | 0 |
| | 4.5 | 0 | 0 | 0 | 0.07 | 0 | 0.06 | 0 |
| | 5 | 0 | 0 | 0 | 0.1 | 0 | 0.05 | 0 |

Table SI6: **Coefficients of variation** for the simulations computed from the 10000 parameter vectors accepted in the last iteration of the initial fitting algorithm. Only the fates with a mean of at least 0.05 are shown.

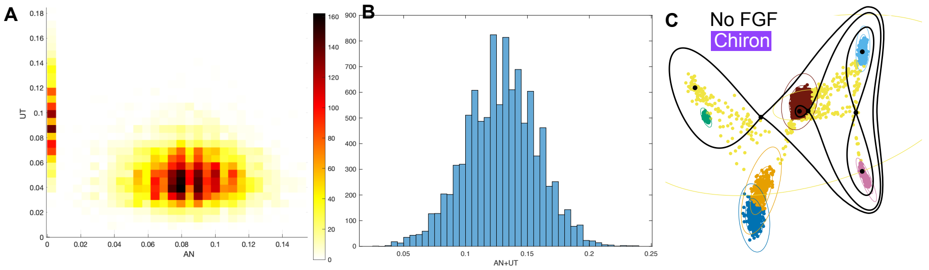

Figure SI25: Analysis of fate bimodality for AN and UT under CHIR 2-5 FGF 0-3 conditions. A. Relative proportion distributions for AN and UT under CHIR 2-5 FGF 0-3 regime simulations at day 5. B. Distribution of the sum of AN and UT at day 5. C. Landscape for PD and CHIR conditions defined by a parameter vector giving no AN cells at day 5 under CHIR 2-5 FGF 0-3 regime. The coloured level curves show the different components of the Gaussian mixture model fitted to the simulated data corresponding to the same parameter vector. Coloured points correspond to simulated cells for all training experimental conditions. Colours correspond to cell identities as in Figure 3B.

##### 5.1.2 Validation Data

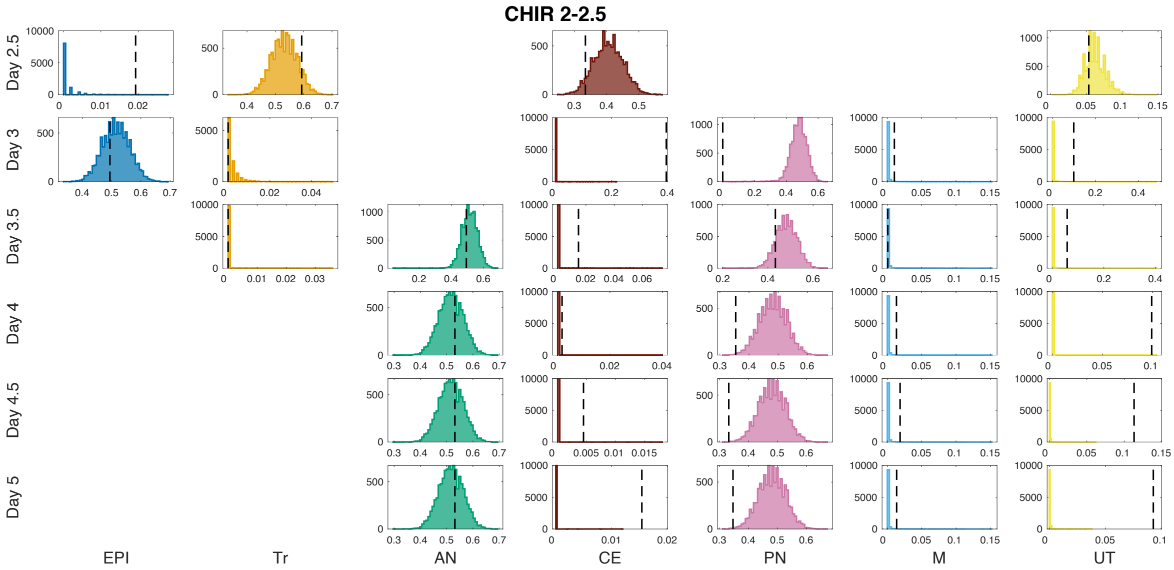

Figure SI26: Proportion distributions for CHIR 2-2.5 regime simulations using the 10000 parameter vectors accepted in the last round of the initial fitting algorithm. The vertical dotted lines indicate the proportion from the initial experimental series used as validation.

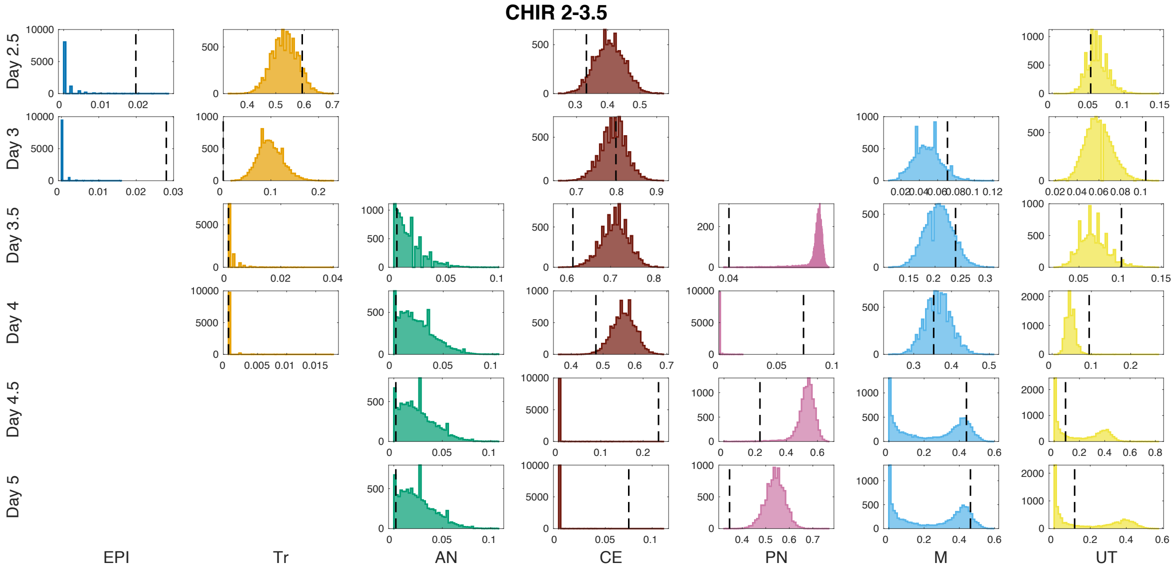

Figure SI27: Proportion distributions for CHIR 2-3.5 regime simulations using the 10000 parameter vectors accepted in the last iteration of the initial fitting algorithm. The vertical dotted lines indicate the proportion from the initial experimental series used as validation.

In figures SI26 to SI29 we see that most of the distributions are unimodal. That is not the case for the distributions of M and UT in the CHIR 2-3.5 regime. These fates correspond to the largest coefficients of variation in table SI7. This was caused by the misclassification of M cells as UT by the GMM model as explained above for

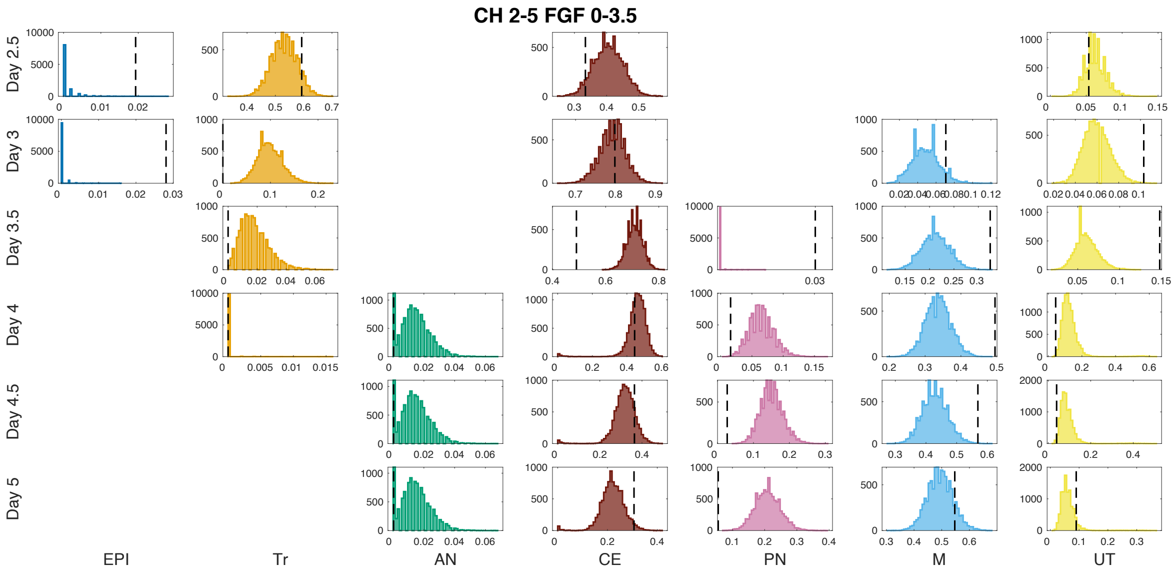

Figure SI28: Proportion distributions for CHIR 2-5 FGF 0-3.5 regime simulations using the 10000 parameter vectors accepted in the last iteration of the initial fitting algorithm. The vertical dotted lines indicate the proportion from the initial experimental series used as validation.

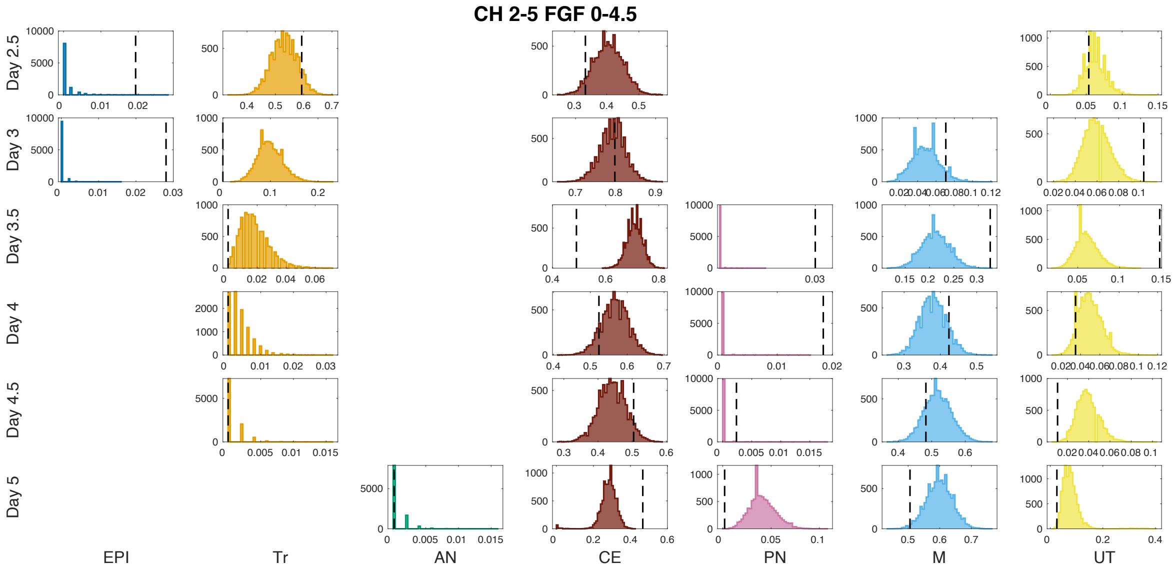

Figure SI29: Proportion distributions for CHIR 2-5 FGF 0-4.5 regime simulations using the 10000 parameter vectors accepted in the last iteration of the initial fitting algorithm. The vertical dotted lines indicate the proportion from the initial experimental series used as validation.

the CHIR 2-3 case. Also the AN distribution for CH 2-5 FGF 0-3.5 is bimodal. The coefficient of variation is not computed for this fate since almost all proportions are below 0.05 and the coefficient of variation is not informative in that case. Nevertheless, the corresponding variation coefficient for UT is quite large in this case. This is caused by the misclassification of AN and UT cells under CHIR + PD conditions as discussed above.

| Condition | Day | EPI | Tr | AN | CE | PN | M | UT |
|---|---|---|---|---|---|---|---|---|
| CHIR 2-2.5 | 2.5 | 0 | 0.09 | 0 | 0.11 | 0 | 0 | 0.23 |
| | 3 | 0.09 | 0 | 0 | 0 | 0.11 | 0 | 0 |
| | 3.5 | 0 | 0 | 0.09 | 0 | 0.1 | 0 | 0 |
| | 4 | 0 | 0 | 0.09 | 0 | 0.1 | 0 | 0 |
| | 4.5 | 0 | 0 | 0.09 | 0 | 0.1 | 0 | 0 |
| | 5 | 0 | 0 | 0.09 | 0 | 0.1 | 0 | 0 |
| CHIR 2-3.5 | 2.5 | 0 | 0.09 | 0 | 0.11 | 0 | 0 | 0.23 |
| | 3 | 0 | 0.27 | 0 | 0.04 | 0 | 0 | 0.21 |
| | 3.5 | 0 | 0 | 0 | 0.04 | 0 | 0.13 | 0.25 |
| | 4 | 0 | 0 | 0 | 0.06 | 0 | 0.09 | 0 |
| | 4.5 | 0 | 0 | 0 | 0 | 0.11 | 0.73 | 0.9 |
| | 5 | 0 | 0 | 0 | 0 | 0.08 | 0.73 | 0.94 |
| CH 2-5 FGF 2-3.5 | 2.5 | 0 | 0.09 | 0 | 0.11 | 0 | 0 | 0.23 |
| | 3 | 0 | 0.27 | 0 | 0.04 | 0 | 0 | 0.21 |
| | 3.5 | 0 | 0 | 0 | 0.04 | 0 | 0.13 | 0.22 |
| | 4 | 0 | 0 | 0 | 0.14 | 0.30 | 0.11 | 0.45 |
| | 4.5 | 0 | 0 | 0 | 0.17 | 0.21 | 0.1 | 0.46 |
| | 5 | 0 | 0 | 0 | 0.21 | 0.19 | 0.09 | 0.48 |
| CH 2-5 FGF 2-4.5 | 2.5 | 0 | 0.09 | 0 | 0.11 | 0 | 0 | 0.23 |
| | 3 | 0 | 0.27 | 0 | 0.04 | 0 | 0 | 0.21 |
| | 3.5 | 0 | 0 | 0 | 0.04 | 0 | 0.13 | 0.22 |
| | 4 | 0 | 0 | 0 | 0.06 | 0 | 0.1 | 0 |
| | 4.5 | 0 | 0 | 0 | 0.09 | 0 | 0.08 | 0 |
| | 5 | 0 | 0 | 0 | 0.16 | 0 | 0.07 | 0.47 |

Table SI7: **Coefficients of variation** for the simulations computed from the 10000 parameter vectors accepted in the last iteration of the initial fitting algorithm. Only the fates with a mean of at least 0.05 are shown.

#### 5.2 Parameter distributions

We present here the empirical distributions of the accepted particles in the last iteration of the fitting algorithm. As mentioned before, we disregarded the weights computed by the fitting algorithm and focused only on the values of the accepted parameters. For the time constants, we show the values of $p_5^{jk}$ and $p_6^{jk}$ instead of $w_{i,5}$ and $w_{i,6}$ because they are more informative on the cells behaviour.

We observe that the amplitude noise parameter in the accepted parameter vectors took small values, of the order $10^{-3} \cdot 10^2 \ell^2/h = 0.1\ \ell^2/h$ where $\ell$ is the arbitrary length unit used in the model, such that the distance between M and PN attractors is approximately $2\ell$. We conjecture that this magnitude was caused by the transitioning populations included in the data. In order to be able to capture the transitioning population Tr at day 2.5, the model required a small noise amplitude to ensure the cells remained in the correct area of the landscape. As a consequence of the small value of the noise amplitude, the CE attractor is very small under CHIR induction conditions, since this small amount of noise needs to be able to drive cells out of the attractor and into the PN and M region.

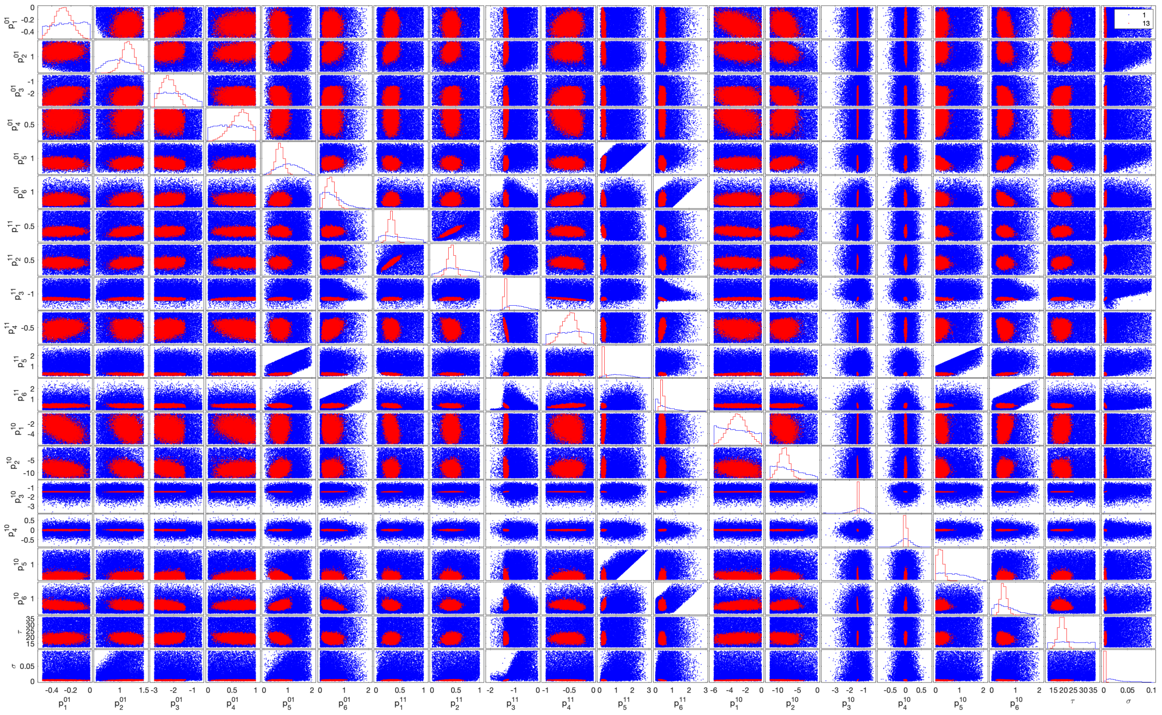

Figure SI30: Distributions of accepted parameters in the first iteration (blue) compared to accepted parameters in the last iteration (red) of the initial fitting algorithm.

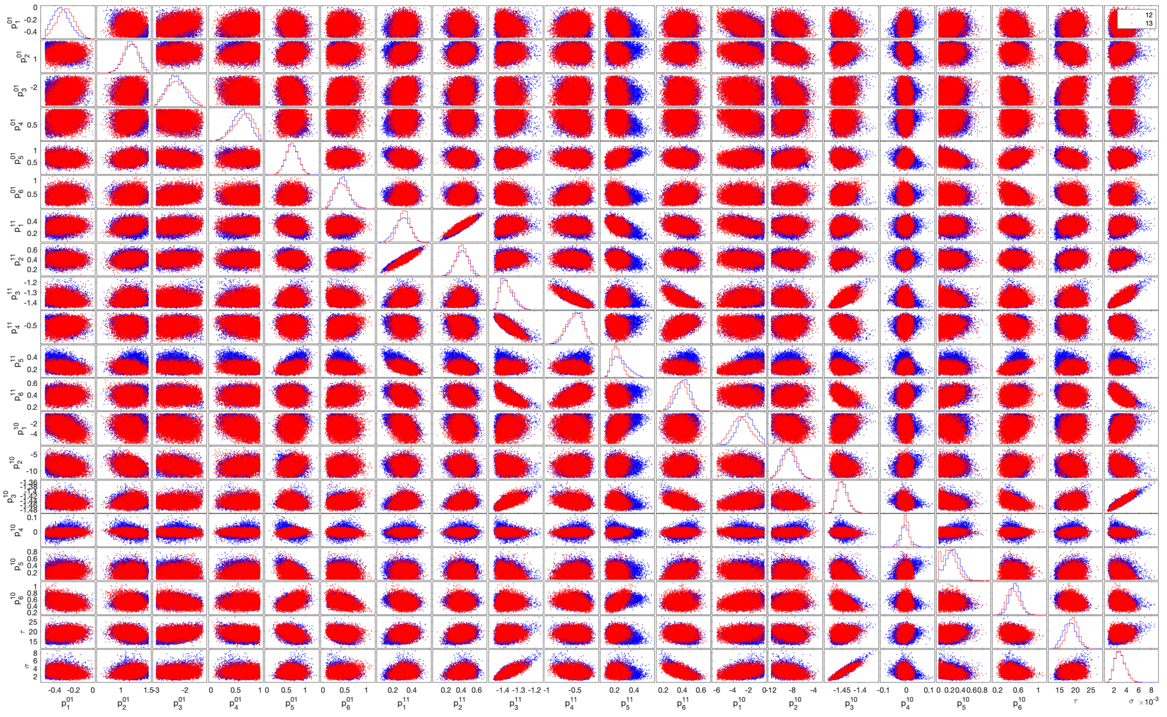

Figure SI31: Distributions of accepted parameters in the previous from last iteration (blue) compared to accepted parameters in the last iteration (red) of the initial fitting algorithm.

The memory parameter $\tau$ took values around 18 (hours), implying that the memory effect was required for the model to perform as accurately as it did. If the memory was not necessary, this parameter would also take much larger values. The larger the value is, the more CHIR is required for the memory to have an effect, and therefore the less effect it finally has in the simulation results.

Most of the parameters show little correlation with others, supporting the minimality in the number of parameters used (Fig. SI32). We observe a high correlation between $\sigma$, the noise amplitude parameter, and $p_3$ under CHIR induction conditions. This is because the size of the CE attractor and the noise amplitude need to be tightly controlled so that cells are able to escape the CE attractor because of the noise effect, as mentioned before. There is also a very high correlation between $p_1^{11}$ and $p_2^{11}$, most likely because to capture the transitioning population accurately we need a very specific form for the landscape of the first decision. In SI33 we see how they align with the bifurcation curve. Similarly, for $p_3^{11}$ and $p_4^{11}$, in order to obtain the correct escape rate from CE, in figure SI34 we see how they align with the bifurcation curve. Furthermore, the range of values that these parameters take is much narrower than for the others. We also note that the velocities for this condition (CHIR=1, FGF=1) are very close to zero with very narrow distributions. This is because in order to maintain the transitioning population for long enough, and the slow escaping from CE, a slow flux is beneficial.

| | $p_1^{01}$ | $p_2^{01}$ | $p_3^{01}$ | $p_4^{01}$ | $p_5^{01}$ | $p_6^{01}$ | $p_1^{11}$ | $p_2^{11}$ | $p_3^{11}$ | $p_4^{11}$ | $p_5^{11}$ | $p_6^{11}$ | $p_1^{10}$ | $p_2^{10}$ | $p_3^{10}$ | $p_4^{10}$ | $p_5^{10}$ | $p_6^{10}$ | $\tau$ | $\sigma$ |
|---|---|---|---|---|---|---|---|---|---|---|---|---|---|---|---|---|---|---|---|---|
| $p_1^{01}$ | | 0.12 | 0.41 | 0.09 | -0.14 | 0.05 | 0.10 | -0.06 | -0.27 | 0.23 | -0.28 | 0.07 | -0.47 | -0.27 | -0.19 | 0.04 | 0.18 | -0.27 | 0.12 | -0.22 |
| $p_2^{01}$ | 0.12 | | 0.15 | 0.26 | 0.20 | 0.20 | 0.29 | 0.22 | 0.11 | -0.01 | -0.24 | -0.09 | -0.24 | -0.21 | 0.23 | -0.23 | 0.06 | -0.15 | -0.24 | 0.25 |
| $p_3^{01}$ | 0.41 | 0.15 | | -0.06 | -0.30 | 0.21 | 0.16 | 0.11 | -0.12 | 0.18 | -0.20 | 0.08 | -0.20 | -0.36 | 0.04 | -0.16 | 0.15 | -0.34 | 0.23 | 0.07 |
| $p_4^{01}$ | 0.09 | 0.26 | -0.06 | | 0.18 | -0.09 | | -0.05 | 0.21 | -0.33 | -0.11 | -0.02 | -0.33 | 0.18 | | -0.02 | 0.16 | 0.07 | -0.14 | -0.01 |
| $p_5^{01}$ | -0.14 | 0.20 | -0.30 | 0.18 | | 0.08 | -0.14 | -0.07 | 0.07 | -0.09 | 0.35 | -0.06 | 0.01 | 0.31 | -0.12 | 0.15 | -0.26 | 0.46 | -0.40 | 0.01 |
| $p_6^{01}$ | 0.05 | 0.20 | 0.21 | -0.09 | 0.08 | | 0.10 | 0.13 | -0.15 | 0.34 | -0.25 | 0.23 | -0.03 | 0.04 | 0.28 | -0.15 | -0.03 | -0.40 | -0.21 | 0.19 |
| $p_1^{11}$ | 0.10 | 0.29 | 0.16 | | -0.14 | 0.10 | | 0.89 | 0.19 | -0.14 | -0.40 | -0.30 | -0.34 | -0.03 | 0.17 | -0.03 | | -0.17 | 0.09 | 0.18 |
| $p_2^{11}$ | -0.06 | 0.22 | 0.11 | -0.05 | -0.07 | 0.13 | 0.89 | | 0.23 | -0.19 | -0.10 | -0.23 | -0.08 | -0.07 | 0.22 | -0.05 | -0.09 | -0.03 | 0.19 | 0.23 |
| $p_3^{11}$ | -0.27 | 0.11 | -0.12 | 0.21 | 0.07 | -0.15 | 0.19 | 0.23 | | -0.85 | 0.13 | -0.73 | 0.30 | -0.19 | 0.65 | -0.14 | -0.34 | 0.08 | 0.04 | 0.74 |
| $p_4^{11}$ | 0.23 | -0.01 | 0.18 | -0.33 | -0.09 | 0.34 | -0.14 | -0.19 | -0.85 | | -0.22 | 0.51 | -0.18 | 0.04 | -0.24 | 0.05 | 0.17 | -0.29 | -0.10 | -0.30 |
| $p_5^{11}$ | -0.28 | -0.24 | -0.20 | -0.11 | 0.35 | -0.25 | -0.40 | -0.10 | 0.13 | -0.22 | | -0.18 | 0.40 | 0.10 | -0.16 | 0.04 | -0.18 | 0.54 | 0.09 | -0.04 |
| $p_6^{11}$ | 0.07 | -0.09 | 0.08 | -0.02 | -0.06 | 0.23 | -0.30 | -0.23 | -0.73 | 0.51 | -0.18 | | -0.13 | 0.12 | -0.46 | 0.09 | 0.39 | -0.07 | -0.05 | -0.61 |
| $p_1^{10}$ | -0.47 | -0.24 | -0.20 | -0.33 | 0.01 | -0.03 | -0.34 | -0.08 | 0.30 | -0.18 | 0.40 | -0.13 | | -0.25 | 0.36 | -0.11 | -0.31 | 0.24 | 0.15 | 0.39 |
| $p_2^{10}$ | -0.27 | -0.21 | -0.36 | 0.18 | 0.31 | 0.04 | -0.03 | -0.07 | -0.19 | 0.04 | 0.10 | 0.12 | -0.25 | | -0.29 | 0.06 | -0.06 | 0.20 | -0.26 | -0.35 |
| $p_3^{10}$ | -0.19 | 0.23 | 0.04 | | -0.12 | 0.28 | 0.17 | 0.22 | 0.65 | -0.24 | -0.16 | -0.46 | 0.36 | -0.29 | | -0.17 | -0.37 | -0.43 | 0.07 | 0.91 |
| $p_4^{10}$ | 0.04 | -0.23 | -0.16 | -0.02 | 0.15 | -0.15 | -0.03 | -0.05 | -0.14 | 0.05 | 0.04 | 0.09 | -0.11 | 0.06 | -0.17 | | 0.09 | 0.02 | 0.16 | -0.20 |
| $p_5^{10}$ | 0.18 | 0.06 | 0.15 | 0.16 | -0.26 | -0.03 | | -0.09 | -0.34 | 0.17 | -0.18 | 0.39 | -0.31 | -0.06 | -0.37 | 0.09 | | -0.06 | 0.11 | -0.39 |
| $p_6^{10}$ | -0.27 | -0.15 | -0.34 | 0.07 | 0.46 | -0.40 | -0.17 | -0.03 | 0.08 | -0.29 | 0.54 | -0.07 | 0.24 | 0.20 | -0.43 | 0.02 | -0.06 | | -0.17 | -0.21 |
| $\tau$ | 0.12 | -0.24 | 0.23 | -0.14 | -0.40 | -0.21 | 0.09 | 0.19 | 0.04 | -0.10 | 0.09 | -0.05 | 0.15 | -0.26 | 0.07 | 0.16 | 0.11 | -0.17 | | |
| $\sigma$ | -0.22 | 0.25 | 0.07 | -0.01 | 0.01 | 0.19 | 0.18 | 0.23 | 0.74 | -0.30 | -0.04 | -0.61 | 0.39 | -0.35 | 0.91 | -0.20 | -0.39 | -0.21 | | |

Figure SI32: Correlations between parameters of the accepted particles in the last iteration of the initial fitting.

Some of the parameter distributions in Figures SI33, SI34 and SI35 are very tight, especially around the bifurcation curves. For example, in the CHIR with PD regime, the joint posterior distribution for the parameters $p_3$ and $p_4$, which correspond to the

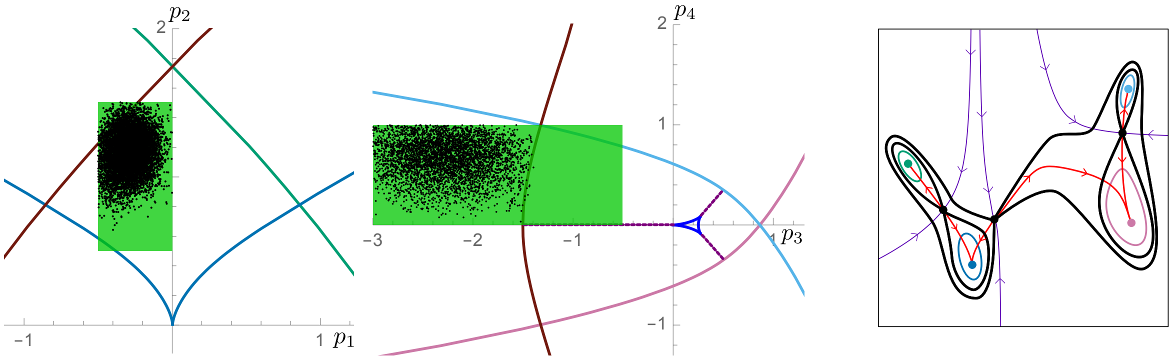

Figure SI33: Left: Priors (green) and empiric distribution after the initial fitting algorithm for the landscape parameters under conditions No CHIR with FGF together with the corresponding bifurcation curves (coloured as bifurcating attractor). Right: average landscape from all accepted parameters under conditions No CHIR with FGF.

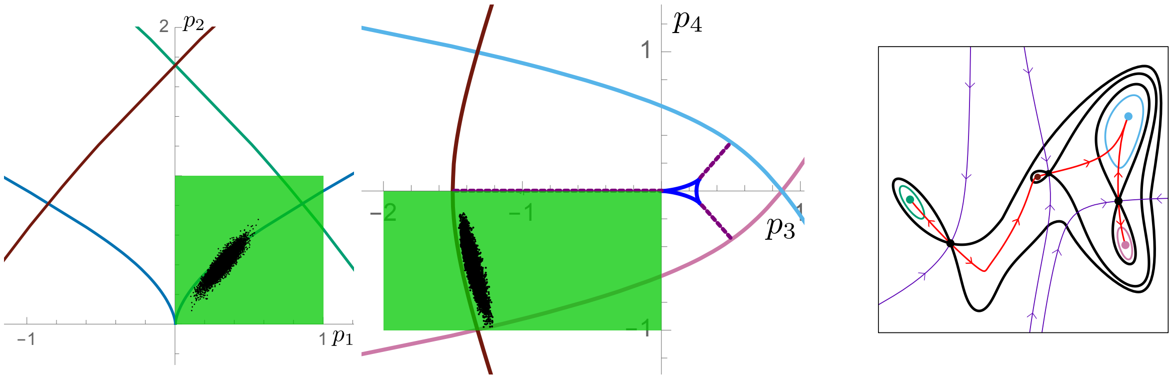

Figure SI34: Left: Priors (green) and empiric distribution after the initial fitting algorithm for the landscape parameters under conditions CHIR with FGF together with the corresponding bifurcation curves (coloured as bifurcating attractor). Right: average landscape from all accepted parameters under conditions CHIR with FGF.

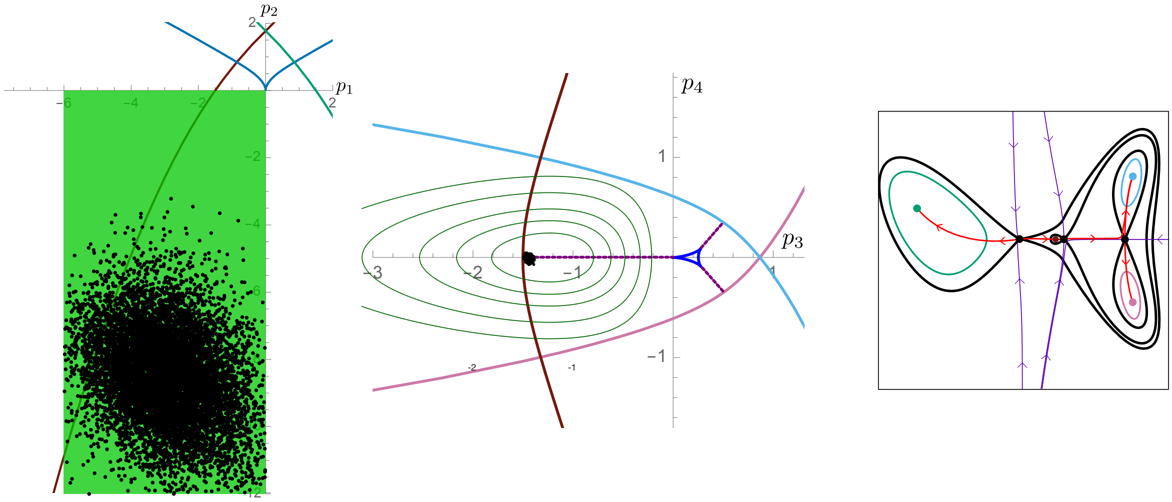

Figure SI35: Left: Priors (green) and empiric distribution after the initial fitting algorithm for the landscape parameters under conditions CHIR with PD together with the corresponding bifurcation curves (coloured as bifurcating attractor). The very tight distributions for $p_3$ and $p_4$ correspond to very tightly regulated features of the landscape: CE is close to bifurcation and the distribution between PN and M fates is almost equal. Contrastingly, the distribution for $p_1$ and $p_2$ is very loose. This is because there is no training data specifically informing these parameters. They are a consequence of the landscape for CHIR with No FGF. The very negative values of $p_2$ causes the AN attractor to significantly move with respect to other signalling profiles. Right: average landscape from all accepted parameters under conditions CHIR with PD.

second decision, is very concentrated. This is because the CE attractor needs to be close to bifurcation, which restricts $p_3$, and the unstable manifold of the saddle from CE to PN/M needs to be very close to the saddle separating M and PN which restricts $p_4$.

### 6 Refining the model

#### 6.1 Refitting for the pulsing experiments

We took as training data the three reference conditions from the Test experimental series (No CHIR, CHIR 2-3, CHIR 2-5) and the two pulsing conditions. We adjusted the priors of the parameters to take into account the information obtained by the first round of fitting. For those parameters that had converged to very tight distributions we adopted that distribution as prior. For parameters with wider distributions we kept the original prior to give sufficient flexibility. The priors that were modified are detailed in table SI8, the rest were kept as in table SI5.

Table SI8: Table of parameter priors modified for the refitting algorithm.

| Parameter | Initial Prior | New Prior |
|---|---|---|
| $p_1^{11}$ | $\mathcal{U}([0,1])$ | $\mathcal{U}([0,0.9])$ |
| $p_2^{11}$ | $\mathcal{U}([0,1])$ | $\mathcal{U}([0.1,0.9])$ |
| $p_3^{11}$ | $\mathcal{U}([-2,0])$ | $N(-1.35,0.06)$ |
| $p_4^{11}$ | $\mathcal{U}([-1,0])$ | $\mathcal{U}([-1,0])$ |
| $p_3^{10}$ | $-\Gamma(9.8,0.14)$ | $N(-1.4,0.03)$ |
| $p_4^{10}$ | $N(0,0.2)$ | $N(0,0.03)$ |
| $\tau$ | $\mathcal{U}([12,36])$ | $N(18,2.5)$ |
| $\sigma$ | $\mathcal{U}([0,0.1])$ | $\mathcal{U}([10^{-5},1.2\cdot 10^{-2}])$ |

We ran the ABC SMC algorithm (refitting) for 12 iterations, changing the acceptance threshold at each round as indicated in figure SI36. We stopped the algorithm at this point because the 13th threshold would have been very close to the 12th and the parameter distributions looked similar in iterations 11 and 12 (Fig. SI45).

We present the results showing first the distributions of cell fate proportions (for the 7 fates and 6 time points) for the 10000 parameter vectors accepted in the last iteration of the algorithm. We do not include day 2 since the distribution of fates corresponds to the initial condition with all cells in the EPI state. We show first the results for the experimental regimes used as training and then the results for the validation regimes. The vertical dotted line indicates the fate proportion coming from the experimental data (test experimental series). Note that they are mostly normal distributions. We show the corresponding variation coefficients in tables SI9 and SI10.

##### 6.1.1 Training Data

In figures SI37 to SI41 we see that most of the distributions are unimodal except for AN and T in the No CHIR regime and M in the CHIR 2-3 regime. These fates together with CE at day 4.5 for CHIR 2-3 and Tr at day 3.5 in the 2 Pulses regime correspond to the largest coefficients of variation in table SI6. The last two coefficients are large because they correspond to fates that were very populated in previous time

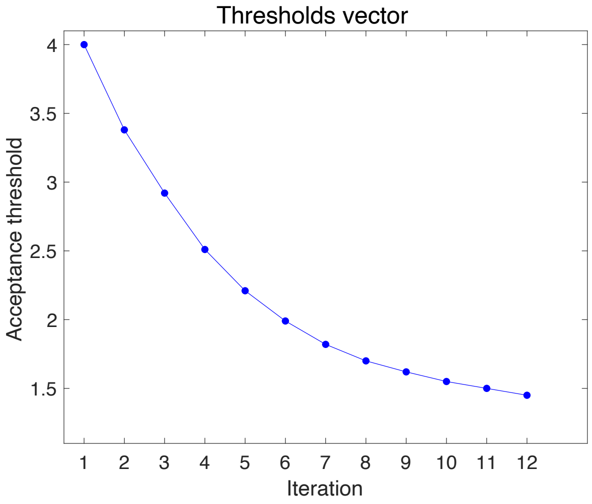

Figure SI36: Evolution of acceptance threshold in the 12 iterations of the refitting algorithm.

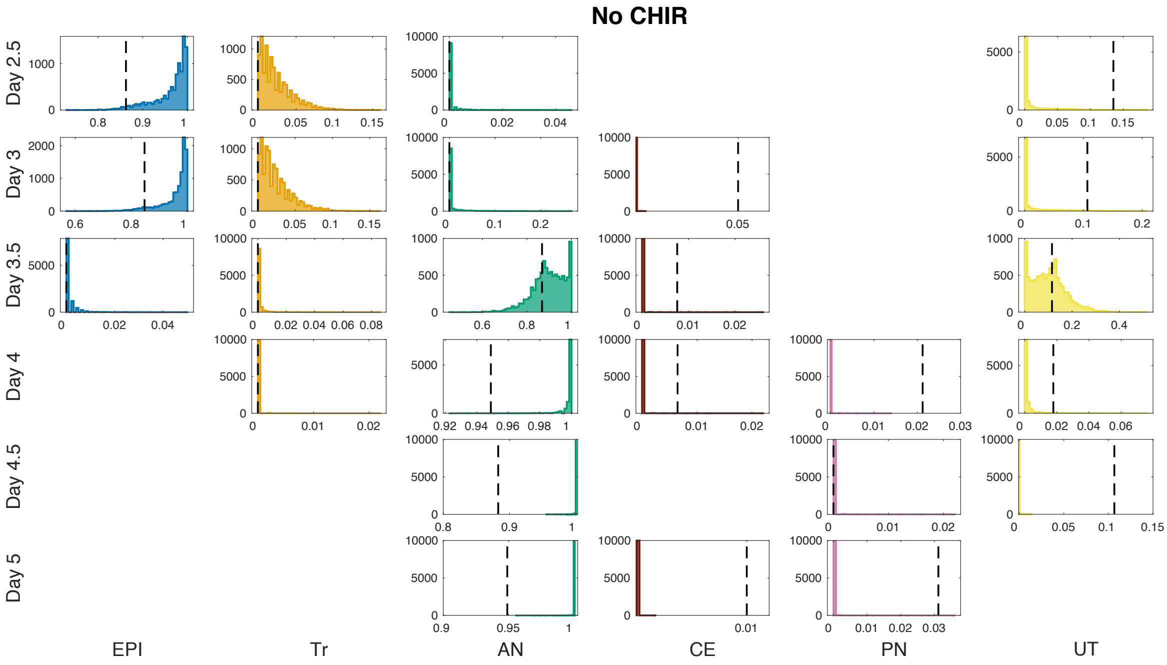

Figure SI37: Proportion distributions for No CHIR regime using the 10000 parameter vectors accepted in the last iteration of the refitting algorithm. The vertical dotted lines indicate the proportion from the test experimental series used as training data. The missing panels correspond to populations with no assigned simulated cells and experimental proportion equal to zero. Note that each panel has different vertical and horizontal ranges.

points and are loosing the cells in favour of other states. That makes the proportion variable, exhibiting a very large variation for a small mean value. The same explains the bimodality in the No CHIR case because for some of the simulations cells took longer to transition from EPI into AN. Regarding the bistability of M in the CHIR 2-3 regime, we find that it is related to the distribution of $p_4^{01}$. We can see in figure

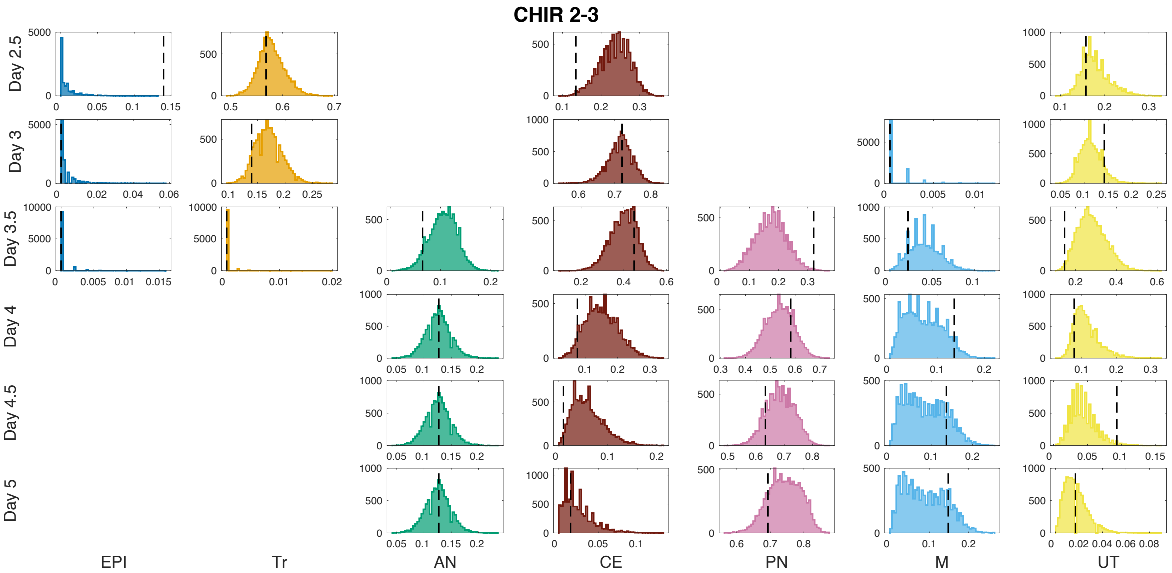

Figure SI38: Proportion distributions for CHIR 2-3 regime simulations in the last iteration of the refitting algorithm. The vertical dotted lines indicate the proportion from the test experimental series used as training.

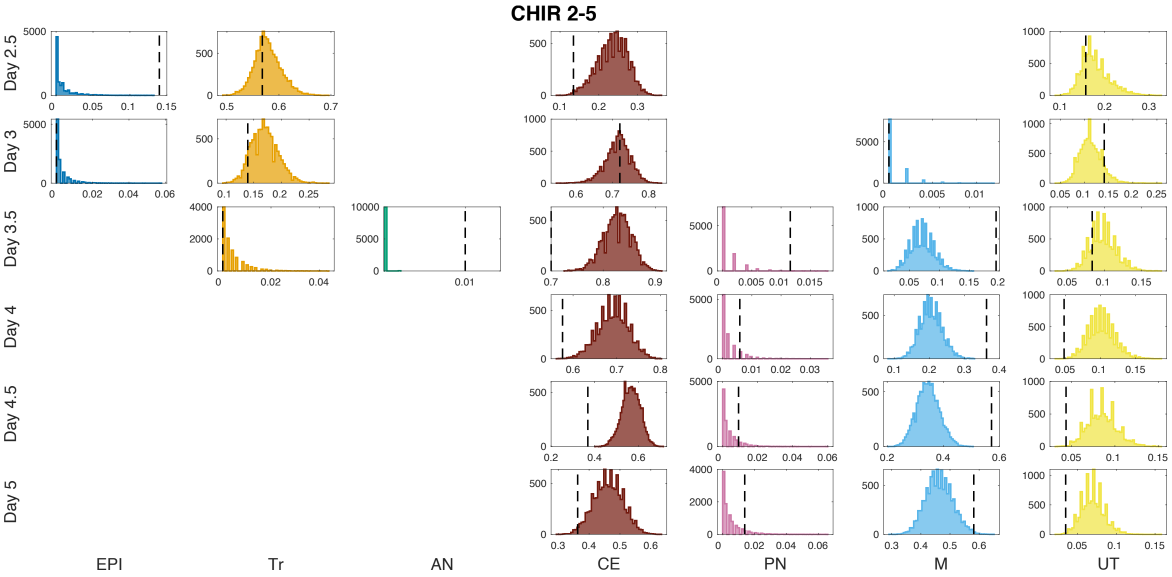

Figure SI39: Proportion distributions for CHIR 2-5 regime simulations in the last iteration of the refitting algorithm. The vertical dotted lines indicate the proportion from the test experimental series used as training.

SI45 that it has a heavy tail. The tail in the distribution gives larger proportions of PN and smaller of M, while values closer to 0 give larger proportions of M as shown in figure SI42. That, together with the different stabilities of CE give rise to a bistable M distribution that corresponds to the wide PN distribution obtained at day 5.

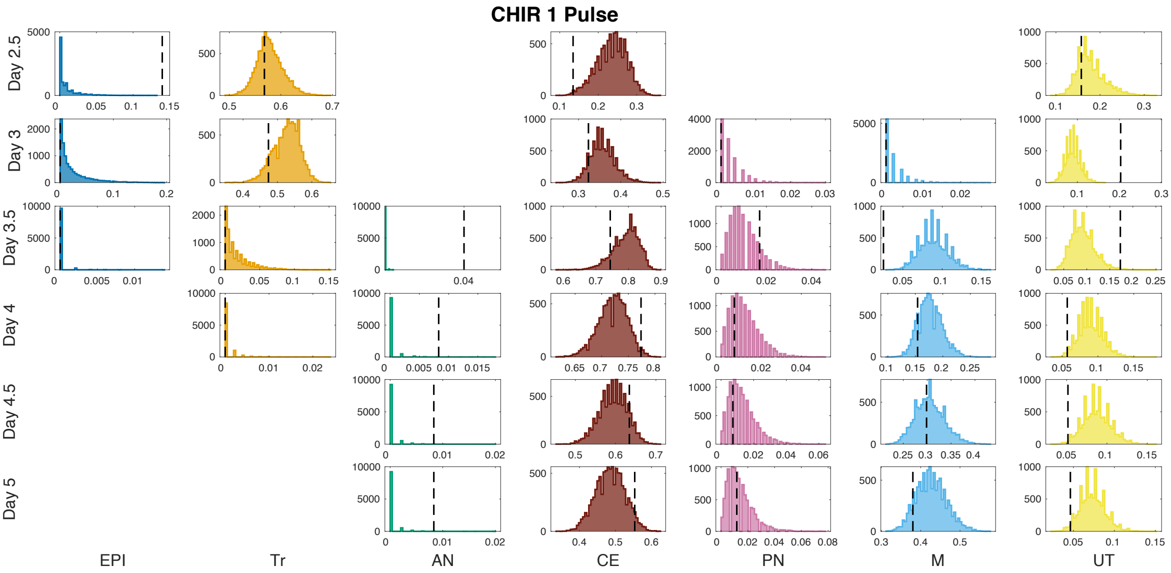

Figure SI40: Proportion distributions for CHIR 1 Pulse regime simulations in the last iteration of the refitting algorithm. The vertical dotted lines indicate the proportion from the test experimental series used as training.

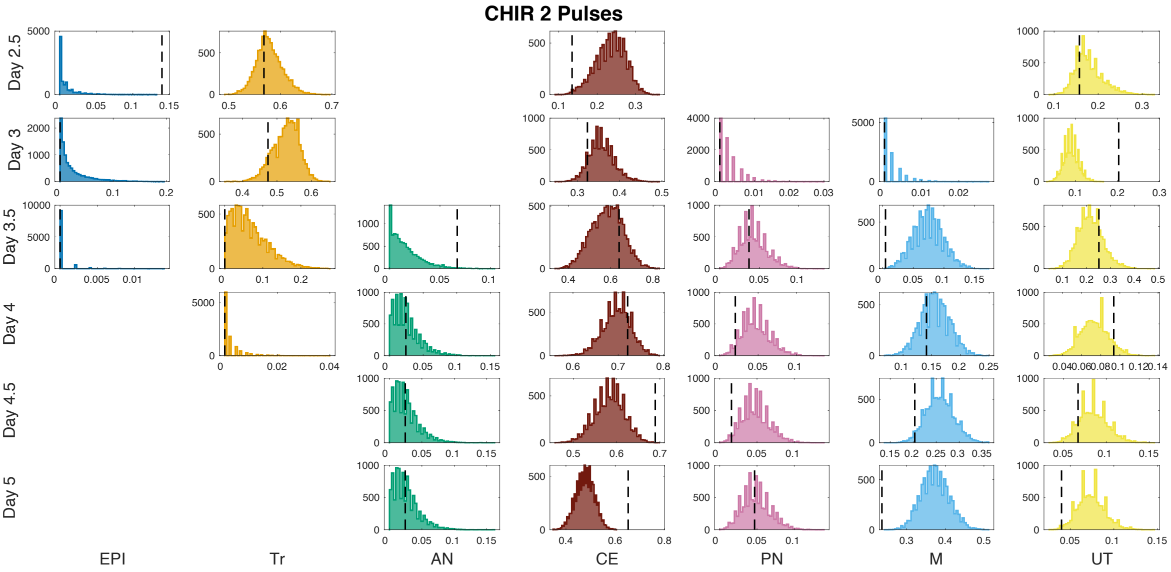

Figure SI41: Proportion distributions for CHIR 2 Pulses regime simulations in the last iteration of the refitting algorithm. The vertical dotted lines indicate the proportion from the test experimental series used as training.

##### 6.1.2 Validation Data

We used the regime CHIR 2-2.5 as validation for the refitting results. Again the model reproduces the recapturing of Tr by the EPI attractor after removal of CHIR, although in this instance the total amount of anterior neural was slightly underestimated by the simulations. We observe that there is a larger amount of M than shown in the data.

In figure SI43 we note that the distribution of M has a thick tail. The tail in the M distribution is caused by the larger values in the $p_4^{01}$, as discussed above. The largest

| Condition | Day | EPI | Tr | AN | CE | PN | M | UT |
|---|---|---|---|---|---|---|---|---|
| No CHIR | 2.5 | 0.04 | 0 | 0 | 0 | 0 | 0 | 0 |
| | 3 | 0.06 | 0 | 0 | 0 | 0 | 0 | 0 |
| | 3.5 | 0 | 0 | 0.08 | 0 | 0 | 0 | 0.69 |
| | 4 | 0 | 0 | 0 | 0 | 0 | 0 | 0 |
| | 4.5 | 0 | 0 | 0 | 0 | 0 | 0 | 0 |
| | 5 | 0 | 0 | 0 | 0 | 0 | 0 | 0 |
| CHIR 2-3 | 2.5 | 0 | 0.04 | 0 | 0.16 | 0 | 0 | 0.17 |
| | 3 | 0 | 0.14 | 0 | 0.05 | 0 | 0 | 0.2 |
| | 3.5 | 0 | 0 | 0.26 | 0.16 | 0.31 | 0 | 0.25 |
| | 4 | 0 | 0 | 0.18 | 0.34 | 0.11 | 0.54 | 0.32 |
| | 4.5 | 0 | 0 | 0.18 | 0.54 | 0.07 | 0.57 | 0 |
| | 5 | 0 | 0 | 0.18 | 0 | 0.07 | 0.58 | 0 |
| CHIR 2-5 | 2.5 | 0 | 0.04 | 0 | 0.16 | 0 | 0 | 0.17 |
| | 3 | 0 | 0.14 | 0 | 0.05 | 0 | 0 | 0.2 |
| | 3.5 | 0 | 0 | 0 | 0.03 | 0 | 0.28 | 0.18 |
| | 4 | 0 | 0 | 0 | 0.05 | 0 | 0.16 | 0.19 |
| | 4.5 | 0 | 0 | 0 | 0.07 | 0 | 0.12 | 0.2 |
| | 5 | 0 | 0 | 0 | 0.1 | 0 | 0.1 | 0.21 |
| CHIR 1 Pulse | 2.5 | 0 | 0.04 | 0 | 0.16 | 0 | 0 | 0.17 |
| | 3 | 0 | 0.07 | 0 | 0.08 | 0 | 0 | 0.2 |
| | 3.5 | 0 | 0 | 0 | 0.05 | 0 | 0.2 | 0.28 |
| | 4 | 0 | 0 | 0 | 0.04 | 0 | 0.12 | 0.18 |
| | 4.5 | 0 | 0 | 0 | 0.06 | 0 | 0.1 | 0.19 |
| | 5 | 0 | 0 | 0 | 0.08 | 0 | 0.09 | 0.2 |
| CHIR 2 Pulses | 2.5 | 0 | 0.04 | 0 | 0.16 | 0 | 0 | 0.17 |
| | 3 | 0 | 0.07 | 0 | 0.08 | 0 | 0 | 0.2 |
| | 3.5 | 0 | 0.71 | 0 | 0.13 | 0 | 0.33 | 0.24 |
| | 4 | 0 | 0 | 0 | 0.04 | 0 | 0.16 | 0.2 |
| | 4.5 | 0 | 0 | 0 | 0.06 | 0 | 0.12 | 0.2 |
| | 5 | 0 | 0 | 0 | 0.08 | 0 | 0.1 | 0.21 |

Table SI9: **Coefficients of variation** for the simulations computed from the 10000 parameter vectors accepted in the last iteration of the refitting algorithm. Only the fates with a mean of at least 0.05 are shown.

coefficients of variation correspond to transitioning populations. We note that the coefficient of variation for M is not computed since the mean value of M is very small and the resulting value is not informative.

##### 6.1.3 Parameter distributions

We present here the empirical distributions of the particles accepted in the last iteration of the refitting algorithm. As mentioned before, we disregarded the weights computed by the fitting algorithm and focused only on the values of the accepted parameters. The obtained parameter distributions are shown in figures SI44 and SI45.

The results together with the priors are shown in figures SI48, SI49 and SI50.

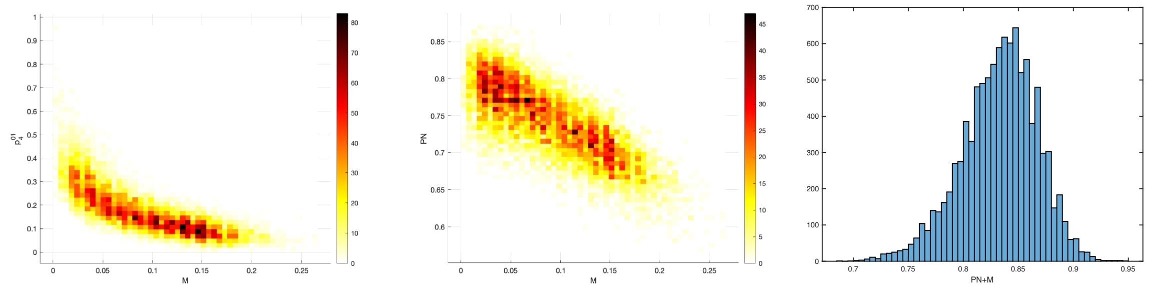

Figure SI42: Analysis of fate bimodality for M under CHIR 2-3 conditions with the parameters obtained in the refitting algorithm. A. Relative distribution of M fate at day 5 with respect to the value of $p_4^{01}$. B. Relative fate distributions for M and PN at day 5. C. Distribution of the sum of PN and M at day 5.

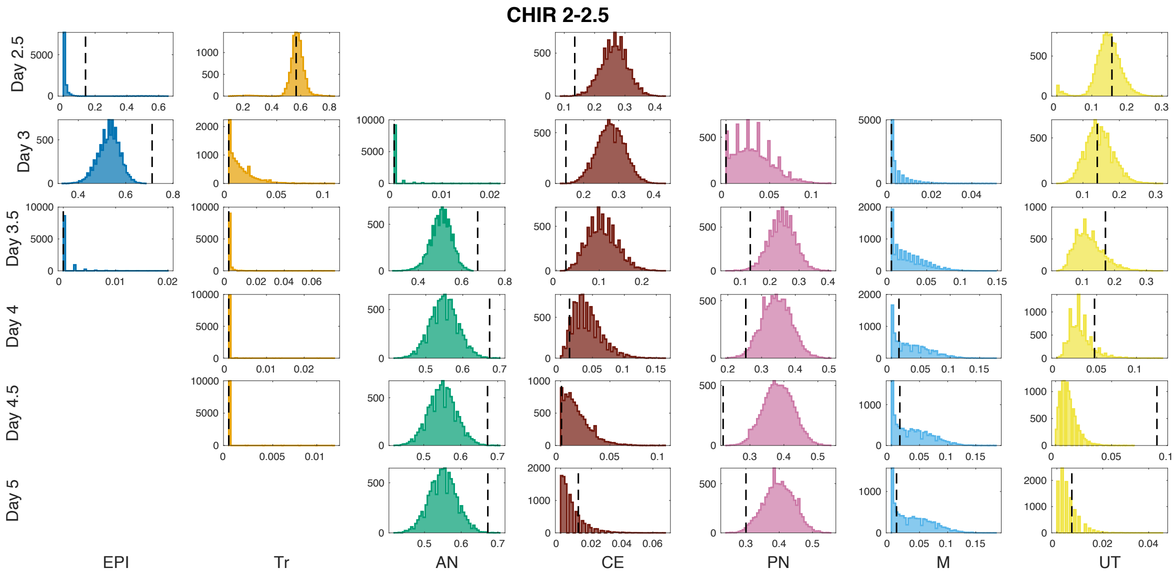

Figure SI43: Proportion distributions for CHIR 2-2.5 regime simulations using the 10000 parameter vectors obtained in the last iteration of the refitting. The vertical dotted lines indicate the proportion from the test experimental series used as validation.

| Condition | Day | EPI | Tr | AN | CE | PN | M | UT |
|---|---|---|---|---|---|---|---|---|
| CHIR 2-2.5 | 2.5 | 0 | 0.10 | 0 | 0.16 | 0 | 0 | 0.26 |
| | 3 | 0.09 | 0 | 0 | 0.14 | 0 | 0 | 0.26 |
| | 3.5 | 0 | 0 | 0.09 | 0.32 | 0.18 | 0 | 0.36 |
| | 4 | 0 | 0 | 0.07 | 0 | 0.13 | 0 | 0 |
| | 4.5 | 0 | 0 | 0.07 | 0 | 0.11 | 0 | 0 |
| | 5 | 0 | 0 | 0.07 | 0 | 0.11 | 0 | 0 |

Table SI10: **Coefficients of variation** for the simulations computed from the 10000 parameter vectors accepted in the last iteration of the refitting algorithm. Only the fates with a mean of at least 0.05 are shown.

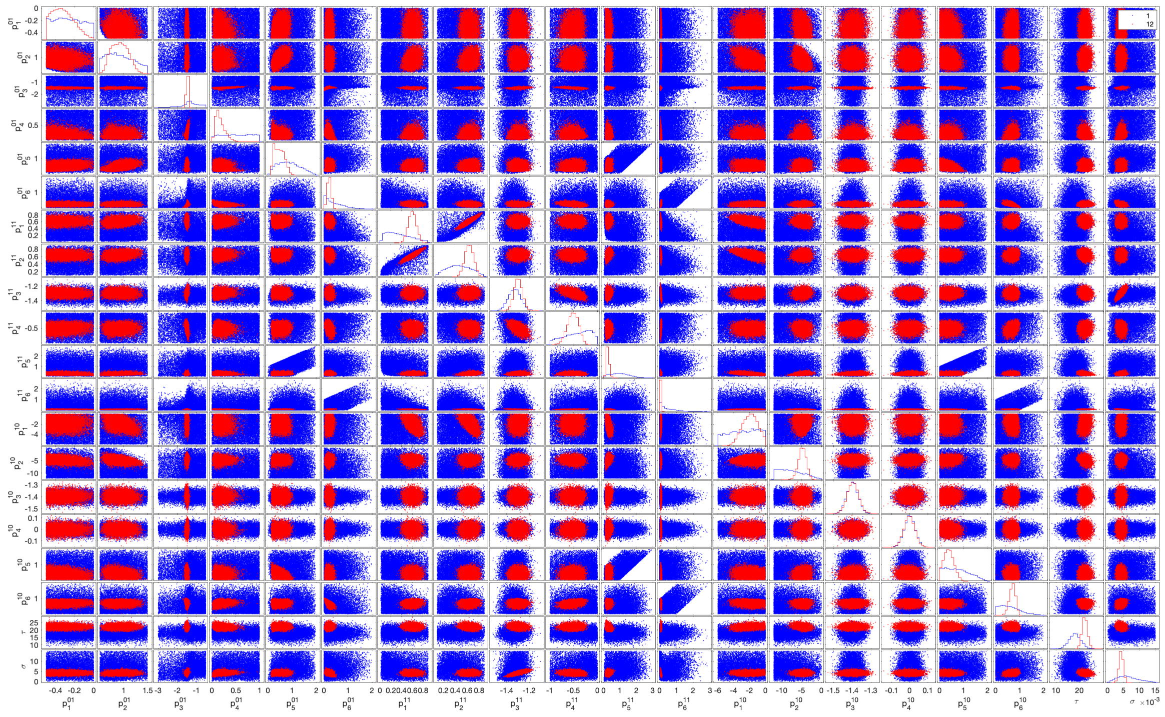

Figure SI44: Distributions of accepted parameters in the first iteration (blue) compared to accepted parameters in the last iteration (red) for the refitting algorithm.

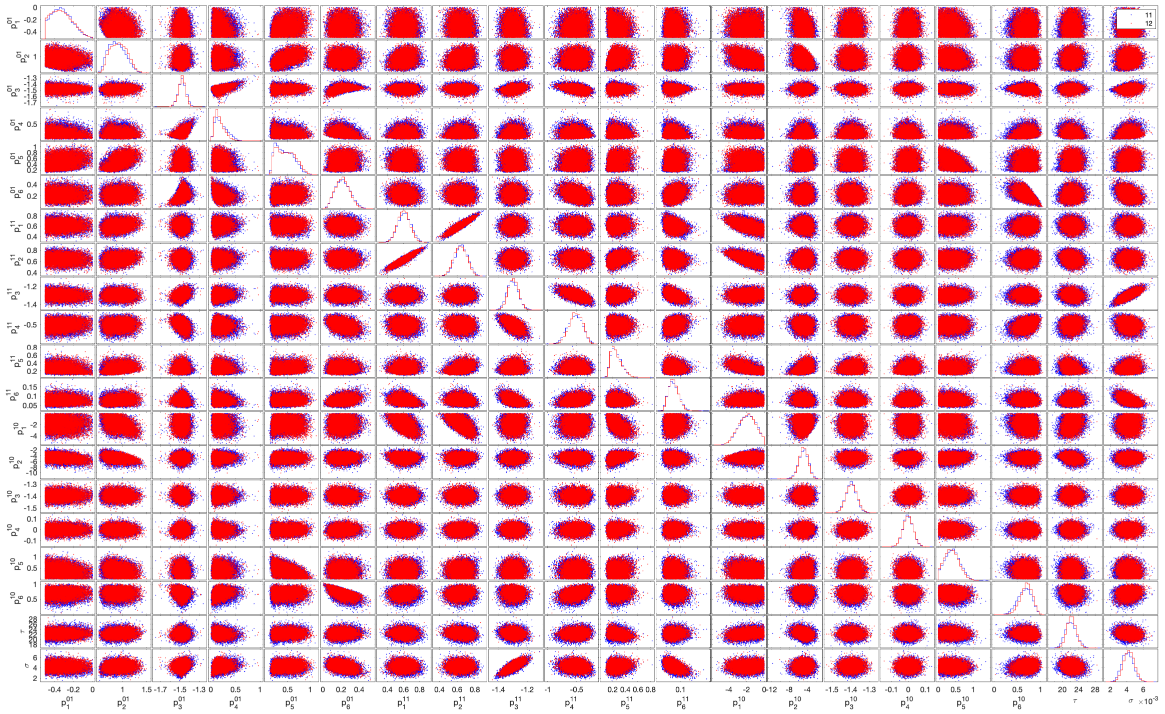

Figure SI45: Distributions of accepted parameters in the previous from last iteration (blue) compared to accepted parameters in the last iteration (red) for the refitting algorithm.

We stopped the fitting algorithm at this point even though the threshold is not as good as for the initial fitting because the parameter distributions are almost identical to the ones in the previous iteration.

| | $p_1^{01}$ | $p_2^{01}$ | $p_3^{01}$ | $p_4^{01}$ | $p_5^{01}$ | $p_6^{01}$ | $p_1^{11}$ | $p_2^{11}$ | $p_3^{11}$ | $p_4^{11}$ | $p_5^{11}$ | $p_6^{11}$ | $p_1^{10}$ | $p_2^{10}$ | $p_3^{10}$ | $p_4^{10}$ | $p_5^{10}$ | $p_6^{10}$ | $\tau$ | $\sigma$ |
|---|---|---|---|---|---|---|---|---|---|---|---|---|---|---|---|---|---|---|---|---|
| $p_1^{01}$ | | -0.03 | -0.03 | -0.15 | 0.34 | 0.02 | 0.02 | -0.06 | -0.07 | 0.10 | -0.12 | -0.05 | -0.01 | -0.26 | 0.05 | 0.02 | -0.19 | 0.05 | 0.01 | |
| $p_2^{01}$ | -0.03 | | 0.05 | -0.02 | 0.63 | 0.05 | 0.23 | 0.21 | -0.04 | 0.05 | 0.23 | 0.16 | -0.46 | -0.35 | -0.05 | -0.03 | -0.23 | | 0.11 | 0.08 |
| $p_3^{01}$ | -0.03 | 0.05 | | 0.42 | 0.07 | 0.48 | -0.05 | -0.03 | 0.27 | -0.41 | -0.02 | -0.01 | -0.01 | 0.04 | -0.16 | 0.07 | -0.12 | -0.18 | 0.05 | 0.15 |
| $p_4^{01}$ | -0.15 | -0.02 | 0.42 | | -0.15 | -0.30 | 0.09 | 0.11 | 0.10 | -0.07 | 0.01 | -0.09 | -0.15 | | 0.05 | -0.08 | -0.01 | 0.19 | -0.10 | 0.19 |
| $p_5^{01}$ | 0.34 | 0.63 | 0.07 | -0.15 | | 0.13 | 0.03 | 0.05 | -0.04 | 0.06 | 0.20 | 0.13 | -0.06 | 0.05 | -0.04 | 0.07 | -0.48 | 0.01 | 0.06 | 0.12 |
| $p_6^{01}$ | 0.02 | 0.05 | 0.48 | -0.30 | 0.13 | | -0.12 | -0.08 | -0.04 | -0.37 | 0.04 | 0.30 | 0.05 | 0.06 | -0.09 | 0.14 | -0.04 | -0.58 | 0.05 | -0.34 |
| $p_1^{11}$ | 0.02 | 0.23 | -0.05 | 0.09 | 0.03 | -0.12 | | 0.92 | 0.01 | -0.04 | 0.15 | -0.29 | -0.54 | -0.05 | -0.03 | | -0.02 | 0.11 | -0.08 | -0.03 |
| $p_2^{11}$ | -0.06 | 0.21 | -0.03 | 0.11 | 0.05 | -0.08 | 0.92 | | 0.06 | -0.04 | 0.42 | -0.15 | -0.63 | 0.11 | -0.01 | 0.05 | -0.01 | 0.11 | -0.21 | 0.02 |
| $p_3^{11}$ | -0.07 | -0.04 | 0.27 | 0.10 | -0.04 | -0.04 | 0.01 | 0.06 | | -0.59 | 0.10 | -0.46 | 0.06 | 0.08 | -0.01 | -0.04 | 0.02 | -0.03 | -0.14 | 0.76 |
| $p_4^{11}$ | 0.10 | 0.05 | -0.41 | -0.07 | 0.06 | -0.37 | -0.04 | -0.04 | -0.59 | | -0.04 | 0.24 | 0.01 | -0.13 | | 0.05 | | 0.37 | 0.08 | -0.16 |
| $p_5^{11}$ | -0.12 | 0.23 | -0.02 | 0.01 | 0.20 | 0.04 | 0.15 | 0.42 | 0.10 | -0.04 | | -0.03 | -0.60 | 0.38 | 0.12 | 0.15 | 0.06 | 0.01 | -0.25 | 0.11 |
| $p_6^{11}$ | -0.05 | 0.16 | -0.01 | -0.09 | 0.13 | 0.30 | -0.29 | -0.15 | -0.46 | 0.24 | -0.03 | | 0.03 | -0.10 | -0.03 | 0.04 | -0.06 | -0.08 | 0.05 | -0.51 |
| $p_1^{10}$ | -0.01 | -0.46 | -0.01 | -0.15 | -0.06 | 0.05 | -0.54 | -0.63 | 0.06 | 0.01 | -0.60 | 0.03 | | 0.03 | | -0.01 | -0.06 | -0.12 | 0.22 | 0.06 |
| $p_2^{10}$ | -0.26 | -0.35 | 0.04 | | 0.05 | 0.06 | -0.05 | 0.11 | 0.08 | -0.13 | 0.38 | -0.10 | 0.03 | | 0.01 | 0.09 | 0.14 | -0.01 | -0.26 | 0.06 |
| $p_3^{10}$ | 0.05 | -0.05 | -0.16 | 0.05 | -0.04 | -0.09 | -0.03 | -0.01 | -0.01 | | 0.12 | -0.03 | | 0.01 | | 0.04 | 0.07 | -0.02 | -0.02 | 0.04 |
| $p_4^{10}$ | 0.02 | -0.03 | 0.07 | -0.08 | 0.07 | 0.14 | | 0.05 | -0.04 | 0.05 | 0.15 | 0.04 | -0.01 | 0.09 | 0.04 | | -0.10 | | -0.08 | -0.01 |
| $p_5^{10}$ | -0.19 | -0.23 | -0.12 | -0.01 | -0.48 | -0.04 | -0.02 | -0.01 | 0.02 | | 0.06 | -0.06 | -0.06 | 0.14 | 0.07 | -0.10 | | -0.07 | -0.01 | -0.09 |
| $p_6^{10}$ | 0.05 | | -0.18 | 0.19 | 0.01 | -0.58 | 0.11 | 0.11 | -0.03 | 0.37 | 0.01 | -0.08 | -0.12 | -0.01 | -0.02 | | -0.07 | | -0.09 | 0.20 |
| $\tau$ | 0.01 | 0.11 | 0.05 | -0.10 | 0.06 | 0.05 | -0.08 | -0.21 | -0.14 | 0.08 | -0.25 | 0.05 | 0.22 | -0.26 | -0.02 | -0.08 | -0.01 | -0.09 | | -0.17 |
| $\sigma$ | | 0.08 | 0.15 | 0.19 | 0.12 | -0.34 | -0.03 | 0.02 | 0.76 | -0.16 | 0.11 | -0.51 | 0.06 | 0.06 | 0.04 | -0.01 | -0.09 | 0.20 | -0.17 | |

Figure SI46: Correlations between accepted parameters in the last iteration of the refitting algorithm.

The correlations of the parameters are very similar to those obtained in the initial fitting. We note that the large correlation between $p_3^{10}$ has disappeared because there was no data with CHIR + PD in the test dataset, and hence the algorithm could not capture this correlation.

Next, we compare the parameter distributions obtained in the two fitting instances. The distributions are shown in figure SI47. The results together with the priors are shown in figures SI48, SI49 and SI50. We see that the $p_3^{01}$ after the refitting was positioned along the bifurcation curve, giving a drastically different result from the initial fitting with a very tight distribution. Although a bifurcated CE attractor was able to reproduce the initial datasets, a more stable point was required to reproduce the pulsing experiments (in the No CHIR + FGF regime). Also the value of $p_4^{01}$ was smaller after the refitting, benefiting the M attractor. In general, the time constants $p_5^{ij}, p_6^{ij}$ took smaller values, making the transitions between attractors slower. The most noticeable change is in $p_6^{11}$. That increased the stability of CE under CHIR + FGF conditions. In general, we see that the most relevant conclusion from the refitting is that CE needed to be a more stable state to be able to reproduce the pulsing experiments. We can see in the Refitted landscapes (Fig. 8) that CE did not bifurcate under No CHIR + FGF conditions and that even though it bifurcated under No CHIR + No FGF conditions it retained a certain stability (elbow shape of the unstable manifold).

The parameters $p_1^{11}$ and $p_2^{11}$ that control the balance of anterior and posterior fates, mainly in the CHIR 2-2.5 condition, had shifted their position in the bifurcation space and they were positioned more tightly along the bifurcation locus after the refitting, allowing for a larger transitioning population at day 2.5 under CHIR + FGF conditions and producing a slightly larger AN population in the CHIR 2-2.5 condition.

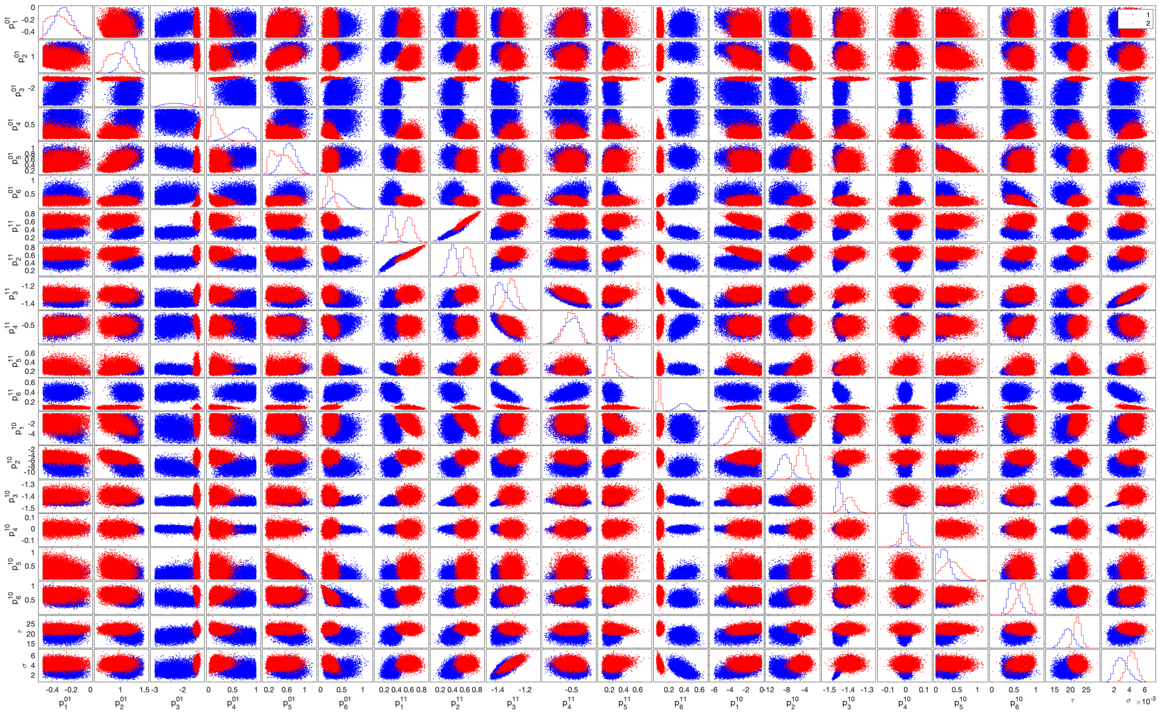

Figure SI47: Distributions of accepted parameters in the last iteration of the initial fitting (blue) compared to accepted parameters in the refitting algorithm (red).

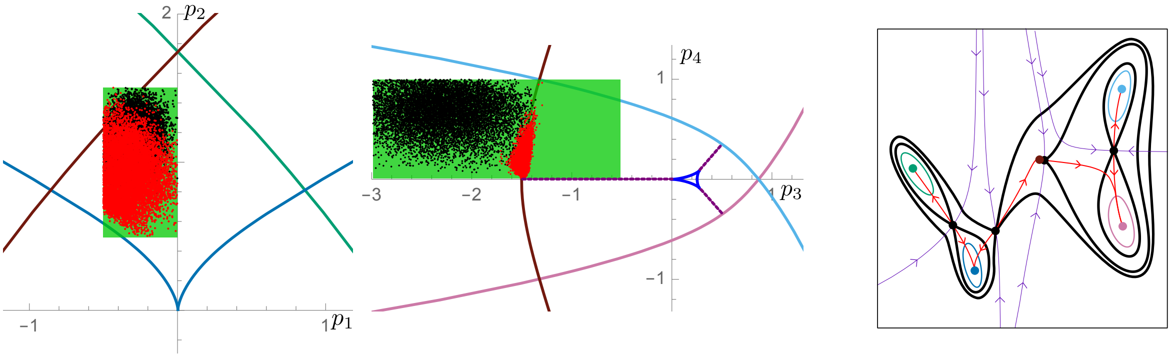

Figure SI48: Left: Priors for the refitting (green) and empiric distributions after initial fitting (black) and refitting (red) algorithms for the landscape parameters under conditions No CHIR with FGF together with the corresponding bifurcation curves (coloured as bifurcating attractor). The marked change in the distribution of $p_3$ corresponds to the fact that the CE attractor is predicted to be clearly more stable after the refitting. Right: average landscape from all accepted parameters under conditions No CHIR with FGF.

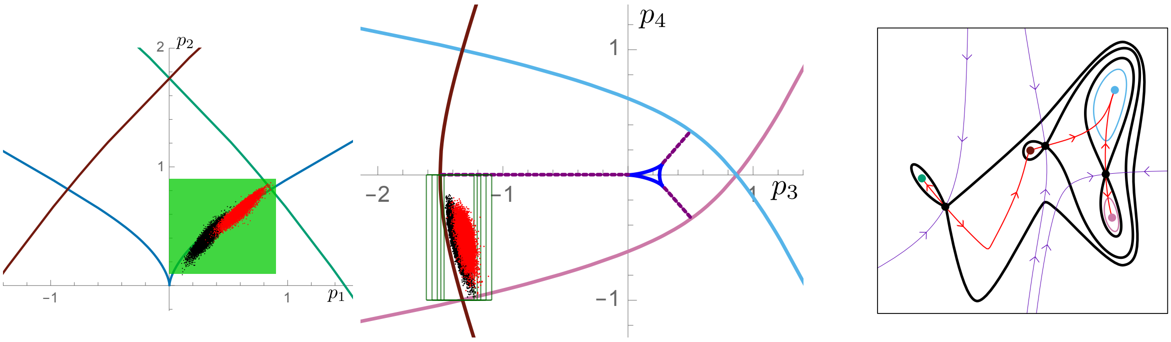

Figure SI49: Left: Priors for the refitting (green) and empiric distribution after initial fitting (black) and refitting (red) algorithms for the landscape parameters under conditions CHIR with FGF together with the corresponding bifurcation curves (coloured as bifurcating attractor). Right: average landscape from all accepted parameters under conditions CHIR with FGF.

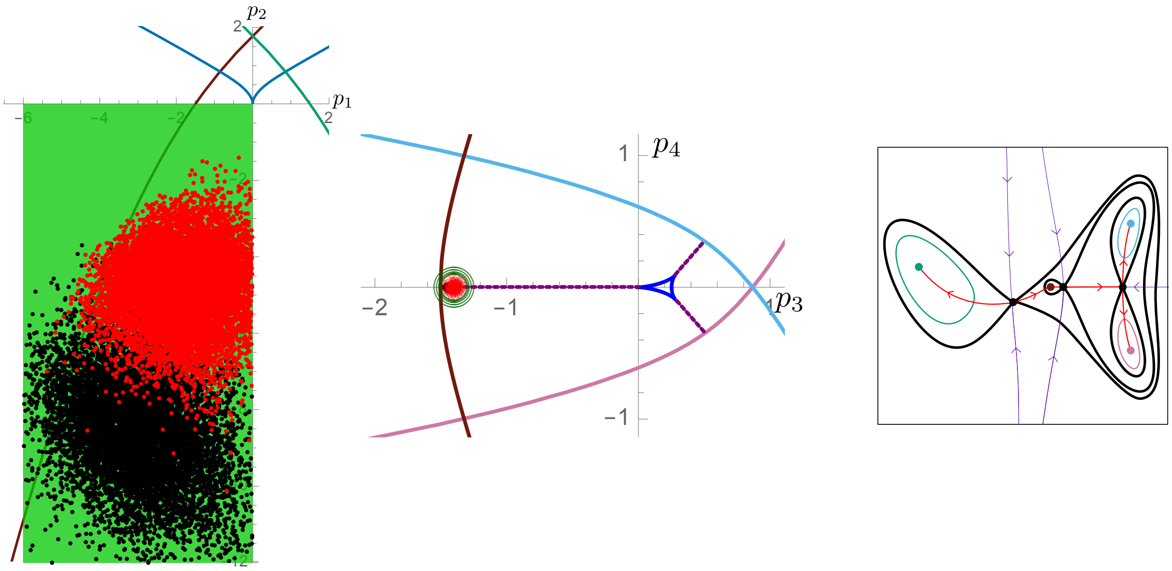

Figure SI50: Left: Priors for the refitting (green) and empiric distribution after initial fitting (black) and refitting (red) algorithms for the landscape parameters under conditions CHIR with PD together with the corresponding bifurcation curves (coloured as bifurcating attractor). Right: average landscape from all accepted parameters under conditions CHIR with PD.

#### 6.2 CHIR dose-response curve

We included a dose-response curve for CHIR concentration in our model to improve the agreement with the experimental data. We decided to use a sigmoid function depending on two parameters $\lambda, \mu$ to model the value of $S_1$ (effective level) as a function of the signal $s_1(t)$ (signal concentrations) as follows:

$$S_1(t) = \frac{tanh(\lambda(s_1(t) + \mu)) + 1}{2}.$$

To perform the fitting of these two parameters while keeping the distributions of the other 20 parameters we ran an ABC SMC algorithm consisting on taking one of the 10000 accepted 20-dim parameter vectors from the refitting and trying to find a pair $(\lambda, \mu)$, sampling from the priors, that would give a good enough approximation of the experimental proportions. We ran the algorithm until we obtained 10000 accepted 22-dimensional parameter vectors. We then changed the threshold, and tried to improve the parameters $(\lambda, \mu)$ while keeping the other 20 dimensions equal.

We ran the algorithm until the distribution of $(\lambda, \mu)$ was almost invariant from one iteration to the next, while the distribution of the other 20 parameters being accepted was changing, that is, many of the accepted parameter vectors by the refitting algorithm were being discarded. We performed 4 rounds of the fitting algorithm changing the acceptance threshold at each round as indicated in figure SI51.

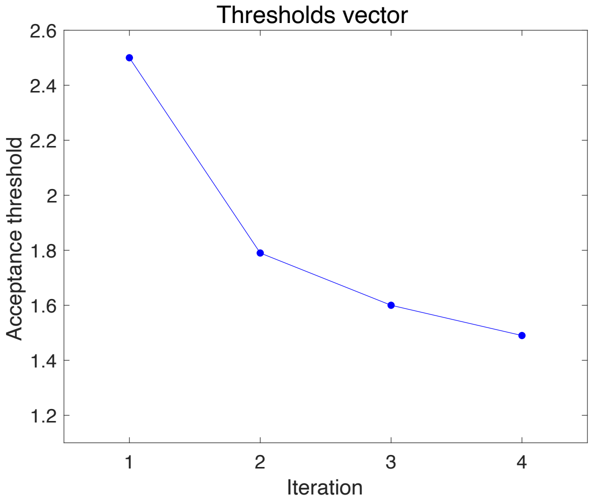

Figure SI51: Evolution of acceptance threshold in the 4 iterations of the Dose-Response fitting algorithm.

##### 6.2.1 Training Data

The distributions are mainly unimodal for all fates. Notably the distribution of M in the CHIR 2-3 condition is unimodal because the DR fitting has concentrated on the smaller values of $p_4^{01}$ disregarding the tail of the distribution discussed above. Under regime CHIR 2-3 0.3 the distributions of M and PN show thick tails. Under CHIR 2-3 0.5 regime the distributions of CE and PN are clearly bimodal. That is a direct consequence of the bistability of the $\mu$ parameter (see figures SI60 and SI58).

| Condition | Day | EPI | Tr | AN | CE | PN | M | UT |
|---|---|---|---|---|---|---|---|---|
| No CHIR | 2.5 | 0.03 | 0 | 0 | 0 | 0 | 0 | 0 |
| | 3 | 0.04 | 0 | 0 | 0 | 0 | 0 | 0 |
| | 3.5 | 0 | 0 | 0.07 | 0 | 0 | 0 | 0.54 |
| | 4 | 0 | 0 | 0 | 0 | 0 | 0 | 0 |
| | 4.5 | 0 | 0 | 0 | 0 | 0 | 0 | 0 |
| | 5 | 0 | 0 | 0 | 0 | 0 | 0 | 0 |
| CHIR 2-3 | 2.5 | 0 | 0.06 | 0 | 0.14 | 0 | 0 | 0.17 |
| | 3 | 0 | 0.13 | 0 | 0.04 | 0 | 0 | 0.17 |
| | 3.5 | 0 | 0 | 0.23 | 0.10 | 0.22 | 0.23 | 0.18 |
| | 4 | 0 | 0 | 0.17 | 0.22 | 0.07 | 0.18 | 0.23 |
| | 4.5 | 0 | 0 | 0.17 | 0.36 | 0.05 | 0.18 | 0 |
| | 5 | 0 | 0 | 0.17 | 0 | 0.04 | 0.17 | 0 |
| CHIR 2-5 | 2.5 | 0 | 0.06 | 0 | 0.14 | 0 | 0 | 0.17 |
| | 3 | 0 | 0.13 | 0 | 0.04 | 0 | 0 | 0.17 |
| | 3.5 | 0 | 0 | 0 | 0.03 | 0 | 0.27 | 0.17 |
| | 4 | 0 | 0 | 0 | 0.05 | 0 | 0.15 | 0.19 |
| | 4.5 | 0 | 0 | 0 | 0.08 | 0 | 0.10 | 0.21 |
| | 5 | 0 | 0 | 0 | 0.10 | 0 | 0.08 | 0.22 |
| CHIR 2-3 0.1 | 2.5 | 0 | 0.06 | 0 | 0.14 | 0 | 0 | 0.17 |
| | 3 | 0 | 0.13 | 0 | 0.04 | 0 | 0 | 0.17 |
| | 3.5 | 0 | 0 | 0.23 | 0.10 | 0.23 | 0.22 | 0.18 |
| | 4 | 0 | 0 | 0.17 | 0.21 | 0.07 | 0.18 | 0.23 |
| | 4.5 | 0 | 0 | 0.17 | 0.35 | 0.05 | 0.17 | 0 |
| | 5 | 0 | 0 | 0.17 | 0 | 0.04 | 0.17 | 0 |
| CHIR 2-3 0.3 | 2.5 | 0 | 0.06 | 0 | 0.14 | 0 | 0 | 0.17 |
| | 3 | 0 | 0.13 | 0 | 0.04 | 0 | 0 | 0.17 |
| | 3.5 | 0 | 0 | 0.47 | 0.11 | 0.42 | 0.24 | 0.17 |
| | 4 | 0 | 0 | 0.27 | 0.24 | 0.23 | 0.26 | 0.28 |
| | 4.5 | 0 | 0 | 0.26 | 0.38 | 0.18 | 0.29 | 0 |
| | 5 | 0 | 0 | 0.26 | 0 | 0.16 | 0.31 | 0 |
| CHIR 2-3 0.5 | 2.5 | 0 | 0.06 | 0 | 0.14 | 0 | 0 | 0.17 |
| | 3 | 0 | 0.13 | 0 | 0.04 | 0 | 0 | 0.17 |
| | 3.5 | 0 | 0 | 0 | 0.08 | 0 | 0.22 | 0.30 |
| | 4 | 0 | 0 | 0 | 0.18 | 1.05 | 0.13 | 0.23 |
| | 4.5 | 0 | 0 | 0 | 0.28 | 0.95 | 0.11 | 0.23 |
| | 5 | 0 | 0 | 0 | 0.39 | 0.92 | 0.11 | 0.28 |

Table SI11: **Coefficients of variation** for the simulations computed from the 10000 parameter vectors accepted in the last iteration of the dose-response fitting algorithm. Only the fates with a mean of at least 0.05 are shown.

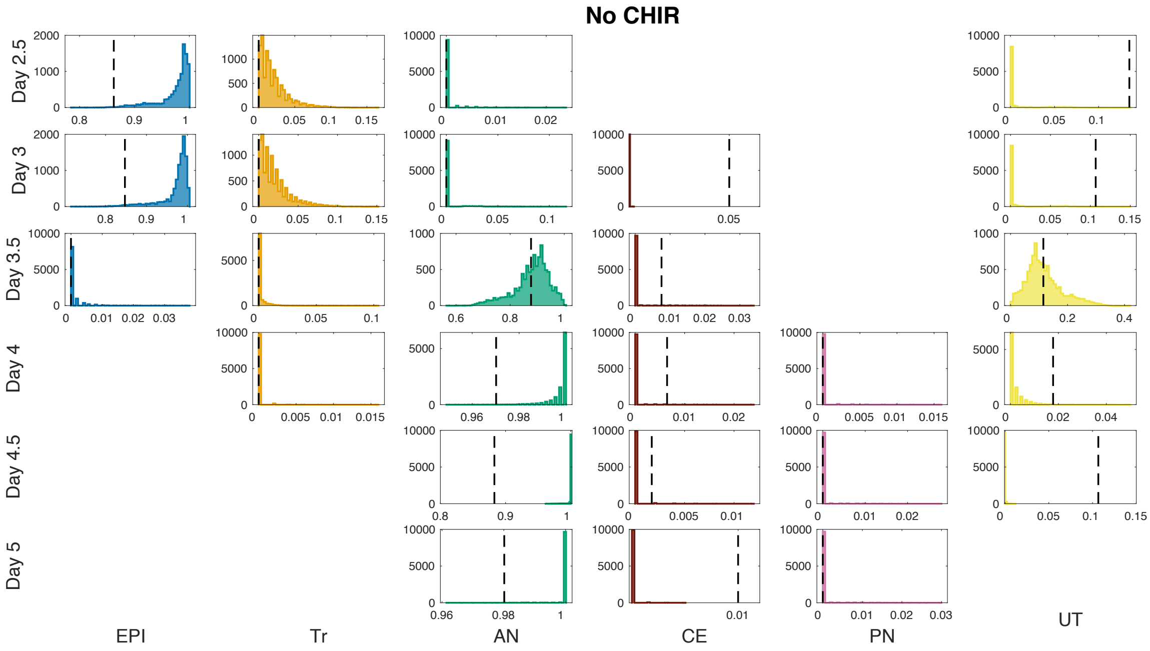

Figure SI52: Proportion distributions for No CHIR regime using the 10000 parameter vectors accepted in the last iteration of the dose-response fitting algorithm. The vertical dotted lines indicate the proportion from the test experimental series. The missing panels correspond to populations with no assigned simulated cells and experimental proportion equal to zero. Note that each panel has different vertical and horizontal ranges.

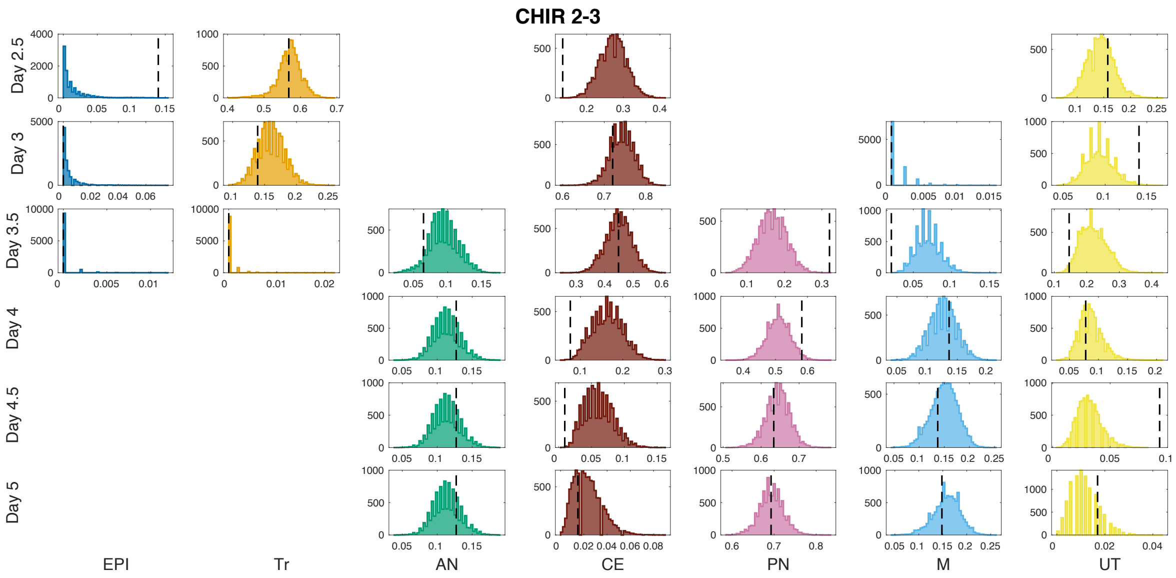

Figure SI53: Proportion distributions for CHIR 2-3 regime simulations in the last iteration of the dose-response fitting algorithm. The vertical dotted lines indicate the proportion from the test experimental series used as training.

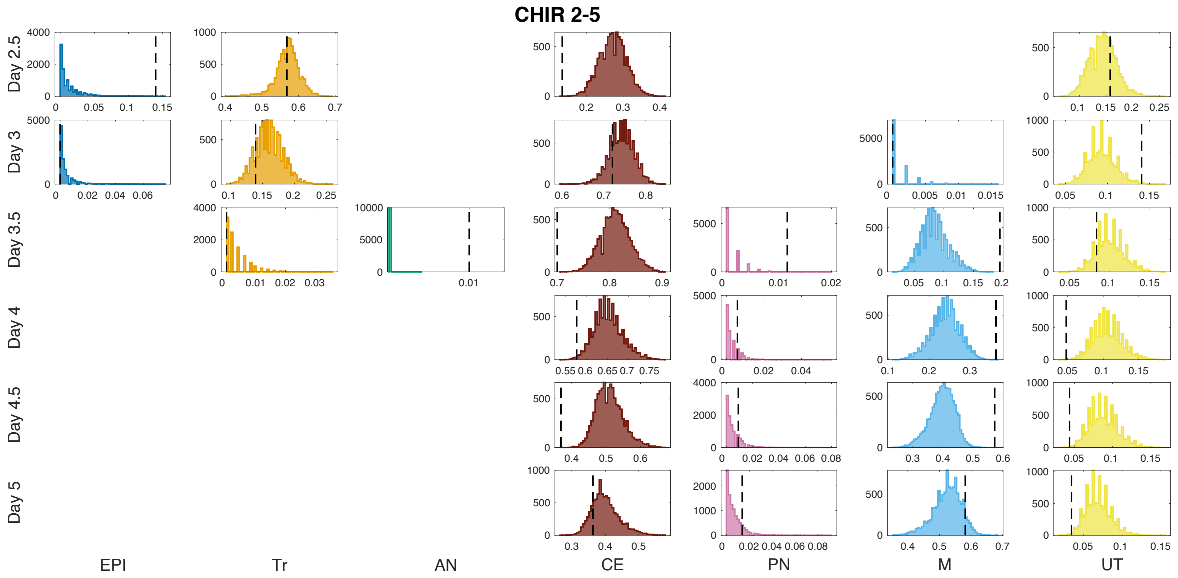

Figure SI54: Proportion distributions for CHIR 2-5 regime simulations in the last iteration of the dose-response fitting algorithm. The vertical dotted lines indicate the proportion from the test experimental series used as training.

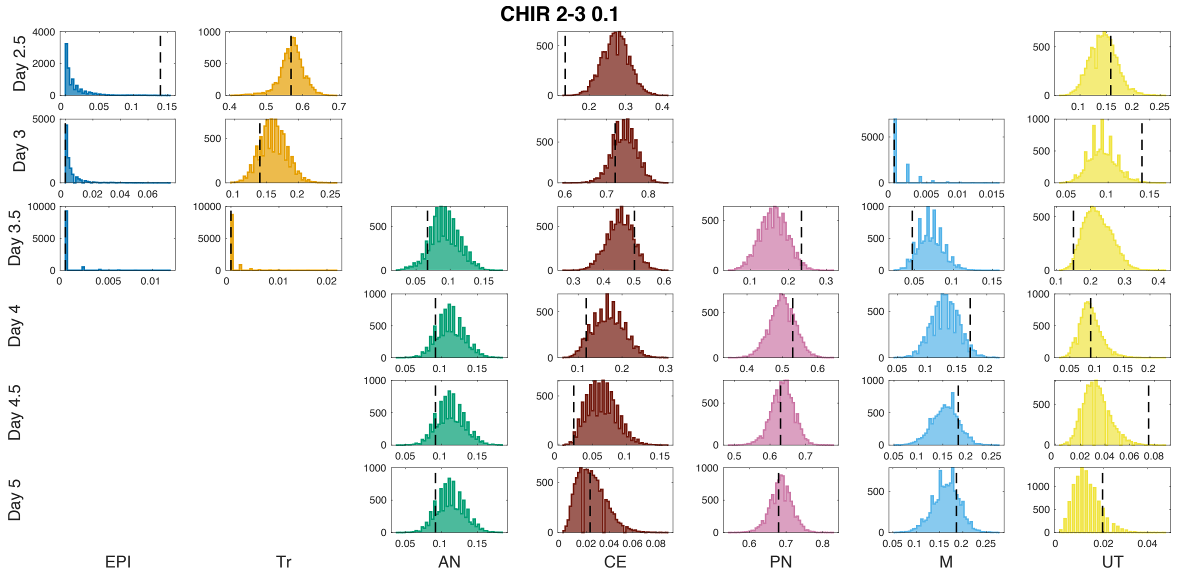

Figure SI55: Proportion distributions for CHIR 2-3 0.1 regime simulations in the last iteration of the dose-response fitting algorithm. The vertical dotted lines indicate the proportion from the test experimental series used as training.

##### 6.2.2 Parameter distributions

We stopped the dose-response fitting algorithm because the distribution of the parameters $\lambda$ and $\mu$ did not change in the last iteration compared to the parameter vectors $(p_1^{01}, \ldots, \sigma)$ discarded in the last iteration. The distributions obtained for the parameters $\lambda$ and $\mu$ were not ideal, which indicates that more points in the dose-response curve would be necessary. As mentioned above the distributions of $p_3^{01}$ and $p_4^{01}$ became tighter in this last iteration compared to the distribution obtained by the refitting

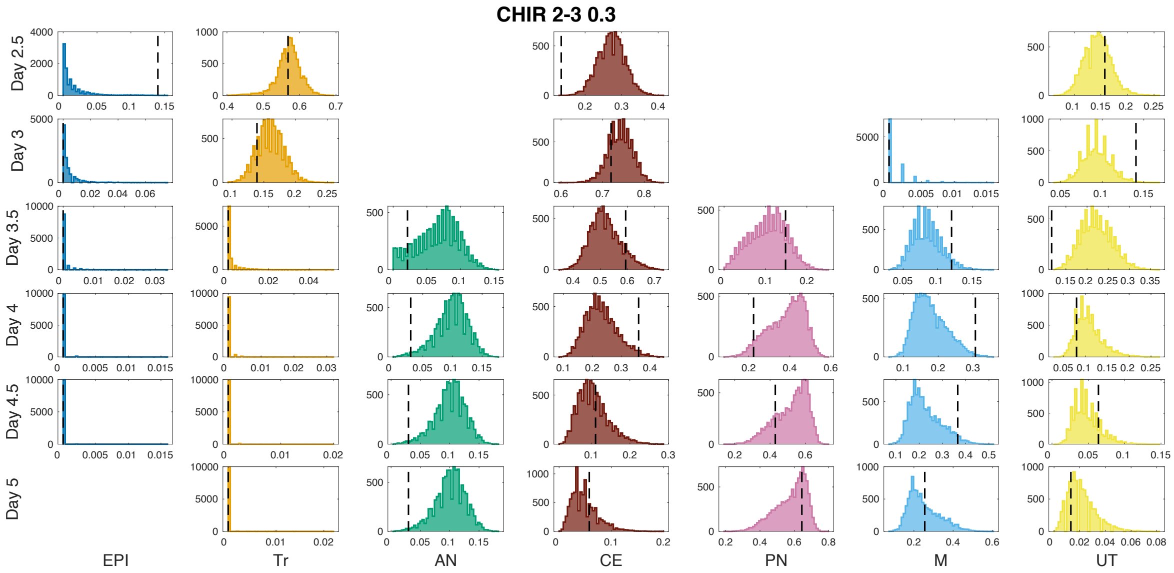

Figure SI56: Proportion distributions for CHIR 2-3 0.3 regime simulations in the last iteration of the dose-response fitting algorithm. The vertical dotted lines indicate the proportion from the test experimental series used as training.

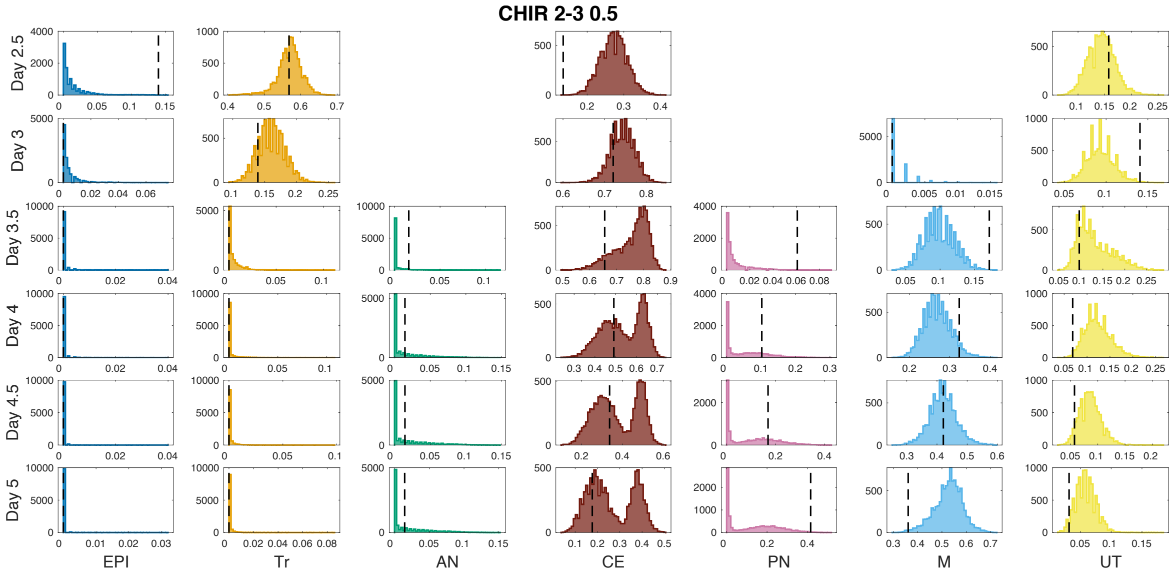

Figure SI57: Proportion distributions for CHIR 2-3 0.5 regime simulations in the last iteration of the dose-response fitting algorithm. The vertical dotted lines indicate the proportion from the test experimental series used as training.

algorithm.

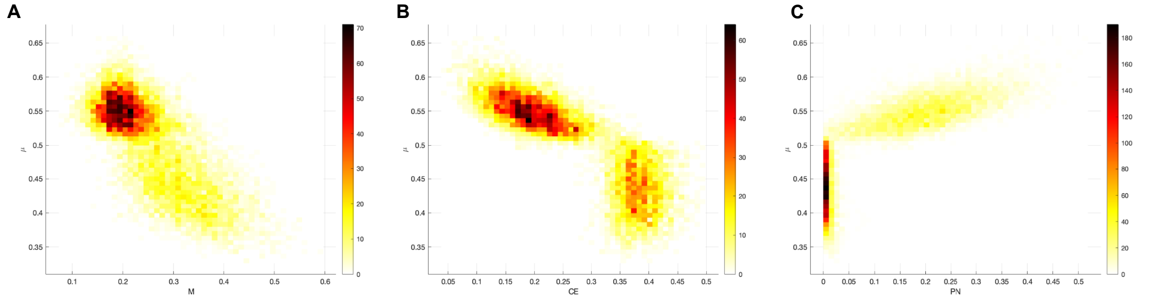

Figure SI58: Analysis of fate bimodality for M, CE and PN in relation to the bimodality of parameter $\mu$. A. Distribution of M at day 5 under CHIR 2-3 0.3 regime in relation to $\mu$. B. Distribution of CE at day 5 under CHIR 2-3 0.5 regime in relation to $\mu$. C. Distribution of PN at day 5 under CHIR 2-3 0.5 regime in relation to $\mu$.

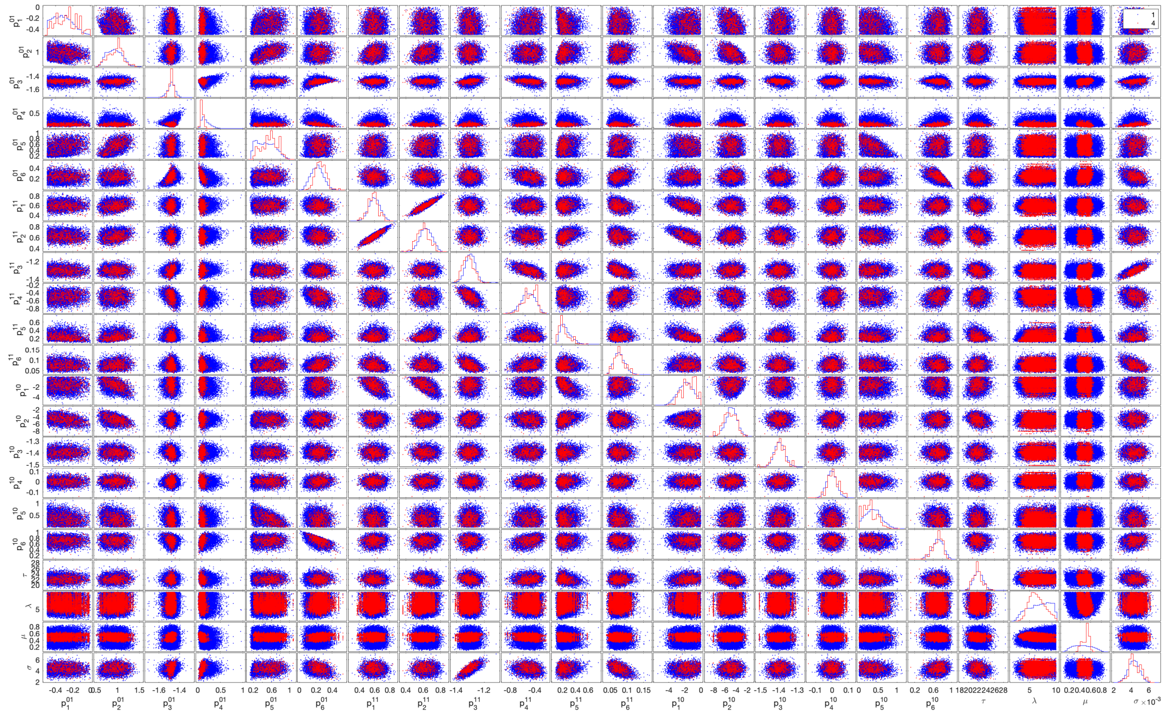

Figure SI59: Distributions of accepted parameters in the first iteration (blue) compared to accepted parameters in the last iteration (red) in the dose-response fitting algorithm.

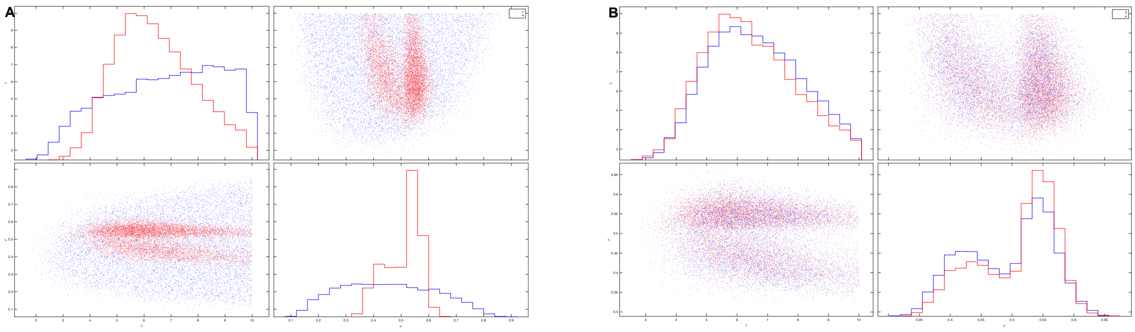

Figure SI60: Distribution for the new parameters ($\lambda$ and $\mu$) added to the model parametrising the dose-response curve. A. Distributions of accepted parameters in the first iteration (blue) compared to accepted parameters in the last iteration (red) in the dose-response fitting algorithm. B. Distributions of accepted parameters in the previous from the last iteration (blue) compared to accepted parameters in the last iteration (red) in the dose-response fitting algorithm.

| | $p_1^{01}$ | $p_2^{01}$ | $p_3^{01}$ | $p_4^{01}$ | $p_5^{01}$ | $p_6^{01}$ | $p_1^{11}$ | $p_2^{11}$ | $p_3^{11}$ | $p_4^{11}$ | $p_5^{11}$ | $p_6^{11}$ | $p_1^{10}$ | $p_2^{10}$ | $p_3^{10}$ | $p_4^{10}$ | $p_5^{10}$ | $p_6^{10}$ | $\tau$ | $\lambda$ | $\mu$ | $\sigma$ |
|---|---|---|---|---|---|---|---|---|---|---|---|---|---|---|---|---|---|---|---|---|---|---|
| $p_1^{01}$ | | -0.25 | -0.08 | 0.02 | 0.21 | -0.15 | -0.12 | -0.26 | -0.16 | 0.31 | -0.35 | 0.07 | 0.20 | -0.46 | 0.15 | | -0.13 | 0.21 | 0.26 | -0.12 | 0.02 | -0.06 |
| $p_2^{01}$ | -0.25 | | 0.12 | 0.03 | 0.53 | 0.11 | 0.29 | 0.23 | -0.07 | 0.05 | 0.25 | 0.15 | -0.55 | -0.35 | -0.08 | -0.02 | -0.13 | -0.26 | 0.12 | 0.07 | -0.09 | 0.03 |
| $p_3^{01}$ | -0.08 | 0.12 | | -0.02 | 0.11 | 0.39 | -0.03 | 0.08 | 0.47 | -0.35 | 0.16 | -0.18 | -0.13 | 0.08 | -0.28 | -0.09 | -0.14 | -0.26 | -0.02 | 0.09 | 0.23 | 0.36 |
| $p_4^{01}$ | 0.02 | 0.03 | -0.02 | | 0.06 | -0.62 | -0.09 | 0.02 | 0.17 | 0.15 | -0.07 | 0.02 | 0.07 | -0.07 | 0.06 | -0.27 | -0.13 | 0.27 | -0.18 | -0.08 | -0.23 | 0.48 |
| $p_5^{01}$ | 0.21 | 0.53 | 0.11 | 0.06 | | 0.09 | 0.11 | 0.11 | -0.11 | 0.11 | 0.20 | 0.11 | -0.13 | -0.05 | 0.01 | | -0.44 | -0.13 | 0.22 | 0.01 | -0.09 | 0.03 |
| $p_6^{01}$ | -0.15 | 0.11 | 0.39 | -0.62 | 0.09 | | 0.04 | 0.08 | -0.11 | -0.27 | 0.26 | 0.12 | -0.19 | 0.20 | -0.05 | 0.12 | 0.13 | -0.62 | | 0.05 | 0.18 | -0.39 |
| $p_1^{11}$ | -0.12 | 0.29 | -0.03 | -0.09 | 0.11 | 0.04 | | 0.87 | -0.05 | -0.10 | 0.09 | -0.26 | -0.48 | 0.10 | -0.12 | 0.01 | -0.08 | -0.20 | -0.10 | | 0.03 | -0.09 |
| $p_2^{11}$ | -0.26 | 0.23 | 0.08 | 0.02 | 0.11 | 0.08 | 0.87 | | 0.12 | -0.26 | 0.45 | -0.27 | -0.61 | 0.34 | -0.19 | 0.01 | -0.05 | -0.25 | -0.30 | 0.06 | 0.02 | 0.04 |
| $p_3^{11}$ | -0.16 | -0.07 | 0.47 | 0.17 | -0.11 | -0.11 | -0.05 | 0.12 | | -0.65 | 0.24 | -0.56 | -0.02 | 0.12 | -0.14 | -0.26 | 0.02 | -0.04 | -0.31 | 0.03 | 0.16 | 0.82 |
| $p_4^{11}$ | 0.31 | 0.05 | -0.35 | 0.15 | 0.11 | -0.27 | -0.10 | -0.26 | -0.65 | | -0.38 | 0.40 | 0.23 | -0.39 | 0.05 | 0.15 | -0.08 | 0.34 | 0.36 | -0.05 | -0.16 | -0.28 |
| $p_5^{11}$ | -0.35 | 0.25 | 0.16 | -0.07 | 0.20 | 0.26 | 0.09 | 0.45 | 0.24 | -0.38 | | -0.19 | -0.58 | 0.44 | -0.10 | 0.18 | 0.08 | -0.24 | -0.33 | 0.12 | | 0.11 |
| $p_6^{11}$ | 0.07 | 0.15 | -0.18 | 0.02 | 0.11 | 0.12 | -0.26 | -0.27 | -0.56 | 0.40 | -0.19 | | 0.20 | -0.27 | 0.22 | 0.16 | -0.08 | 0.05 | 0.13 | 0.02 | -0.12 | -0.46 |
| $p_1^{10}$ | 0.20 | -0.55 | -0.13 | 0.07 | -0.13 | -0.19 | -0.48 | -0.61 | -0.02 | 0.23 | -0.58 | 0.20 | | -0.14 | 0.27 | 0.01 | -0.16 | 0.27 | 0.19 | -0.06 | -0.01 | 0.10 |
| $p_2^{10}$ | -0.46 | -0.35 | 0.08 | -0.07 | -0.05 | 0.20 | 0.10 | 0.34 | 0.12 | -0.39 | 0.44 | -0.27 | -0.14 | | -0.21 | 0.03 | 0.12 | -0.12 | -0.35 | 0.05 | 0.05 | -0.07 |
| $p_3^{10}$ | 0.15 | -0.08 | -0.28 | 0.06 | 0.01 | -0.05 | -0.12 | -0.19 | -0.14 | 0.05 | -0.10 | 0.22 | 0.27 | -0.21 | | | 0.01 | 0.02 | 0.01 | -0.04 | -0.03 | -0.01 |
| $p_4^{10}$ | | -0.02 | -0.09 | -0.27 | | 0.12 | 0.01 | 0.01 | -0.26 | 0.15 | 0.18 | 0.16 | 0.01 | 0.03 | | | -0.07 | 0.09 | 0.05 | 0.04 | -0.02 | -0.27 |
| $p_5^{10}$ | -0.13 | -0.13 | -0.14 | -0.13 | -0.44 | 0.13 | -0.08 | -0.05 | 0.02 | -0.08 | 0.08 | -0.08 | -0.16 | 0.12 | 0.01 | -0.07 | | -0.06 | -0.12 | -0.02 | 0.08 | -0.18 |
| $p_6^{10}$ | 0.21 | -0.26 | -0.26 | 0.27 | -0.13 | -0.62 | -0.20 | -0.25 | -0.04 | 0.34 | -0.24 | 0.05 | 0.27 | -0.12 | 0.02 | 0.09 | -0.06 | | 0.09 | -0.03 | -0.12 | 0.12 |
| $\tau$ | 0.26 | 0.12 | -0.02 | -0.18 | 0.22 | | -0.10 | -0.30 | -0.31 | 0.36 | -0.33 | 0.13 | 0.19 | -0.35 | 0.01 | 0.05 | -0.12 | 0.09 | | -0.03 | -0.14 | -0.22 |
| $\lambda$ | -0.12 | 0.07 | 0.09 | -0.08 | 0.01 | 0.05 | | 0.06 | 0.03 | -0.05 | 0.12 | 0.02 | -0.06 | 0.05 | -0.04 | 0.04 | -0.02 | -0.03 | -0.03 | | -0.21 | 0.04 |
| $\mu$ | 0.02 | -0.09 | 0.23 | -0.23 | -0.09 | 0.18 | 0.03 | 0.02 | 0.16 | -0.16 | | -0.12 | -0.01 | 0.05 | -0.03 | -0.02 | 0.08 | -0.12 | -0.14 | -0.21 | | -0.03 |
| $\sigma$ | -0.06 | 0.03 | 0.36 | 0.48 | 0.03 | -0.39 | -0.09 | 0.04 | 0.82 | -0.28 | 0.11 | -0.46 | 0.10 | -0.07 | -0.01 | -0.27 | -0.18 | 0.12 | -0.22 | 0.04 | -0.03 | |

Figure SI61: Correlations between parameters in the last iteration of the dose-response fitting

### 7 Predictions

Once the model was fitted and validated, it could be used to formulate predictions. In particular we used it to formulate predictions on the behaviour of the system under experimental conditions that were not used in creating the model. The model is parametrised as a continuous function on the signals CHIR and FGF. The model offers predictions for concentrations not tested in the data as part of the family.

#### 7.1 Anterior commitment

The model maintained a saddle separating the anterior and posterior fates under all considered conditions. In particular, under CHIR induction conditions, a posteriorising condition, the saddle was not destroyed. As a consequence, the model predicted (Fig. SI62) that once cells had acquired an AN identity the addition of CHIR was not able to force them to become posterior.

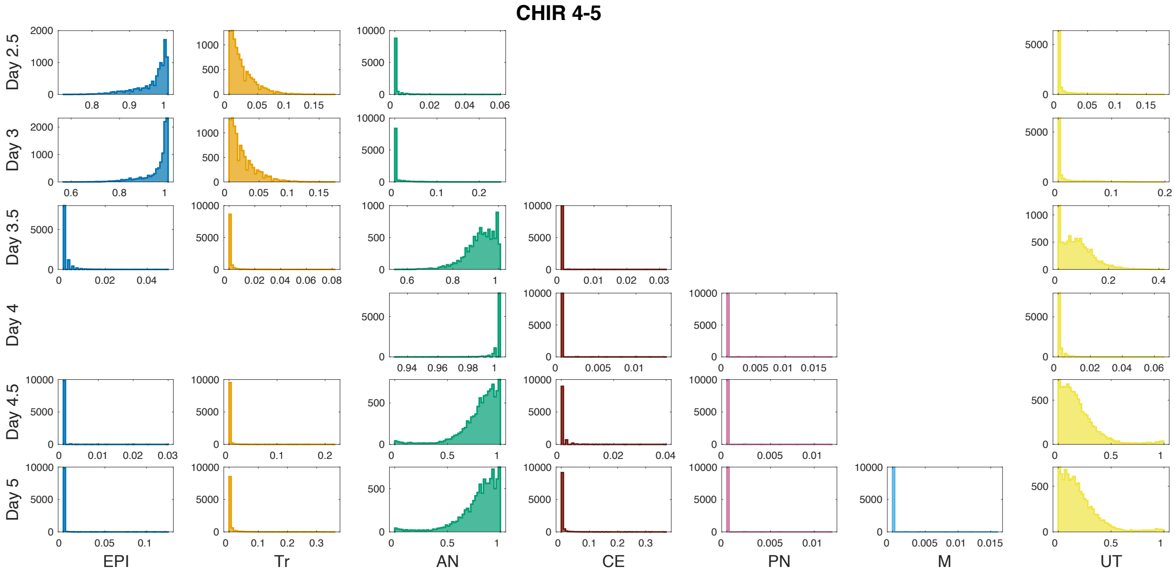

Figure SI62: Proportion distributions for CHIR 4-5 regime simulations using the 10000 parameter vectors obtained in the last iteration of the refitting.

To test this hypothesis we performed an experiment where cells were kept in No CHIR conditions until day 4 and then CHIR and FGF were added to the medium until day 5 (Fig. 10). The data confirmed the model prediction, and showed that the addition of CHIR + FGF at day 4 is not enough to make AN cells transition to more posterior fates as the high levels of Otx2 until day 5 clearly prove. All protein levels distributions are shown in figure SI63.

#### 7.2 CE bifurcation

Figure SI64 shows the cell proportions obtained by the clustering algorithm explained in Section 2 comparing the effect of PD and RA in the population proportions. The complete set of population proportions can be found on Supplementary table T2. The

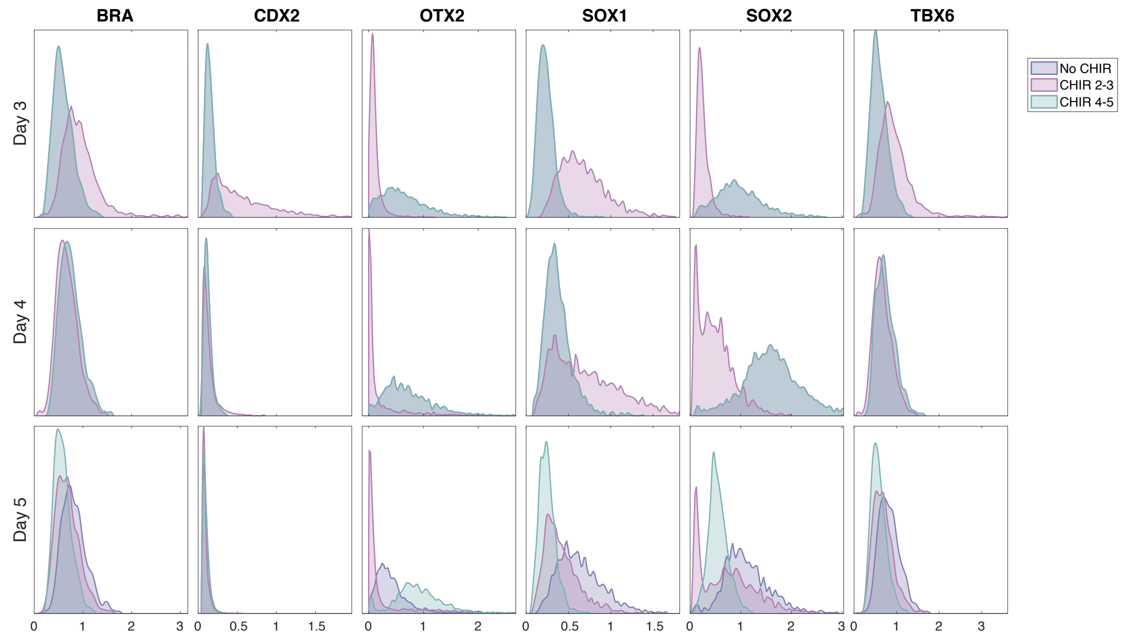

Figure SI63: Distributions comparing the reference experimental conditions No CHIR and CHIR 2-3 with the experimental condition designed to test the commitment to AN. The Otx2 levels for the new condition remain very high at day 5 showing the anterior identity of the cells on these samples.

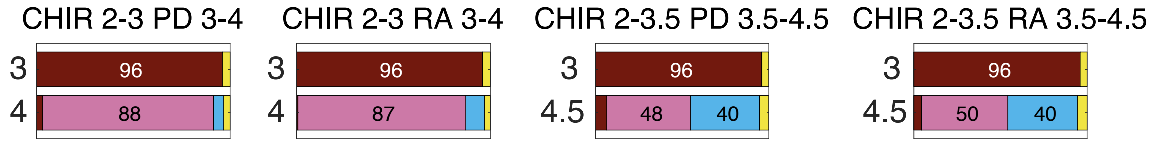

Figure SI64: Clustering proportions comparing the effect of RA and PD as an FGF inhibitor. The proportions are almost identical showing their equivalent effect in this system.

proportions are essentially equal, showing an equivalent effect of these two signals in our system.

Figures SI65 to SI67 show the fate predictions by our model for experiments testing the bifurcation of CE under No CHIR + PD signalling conditions.

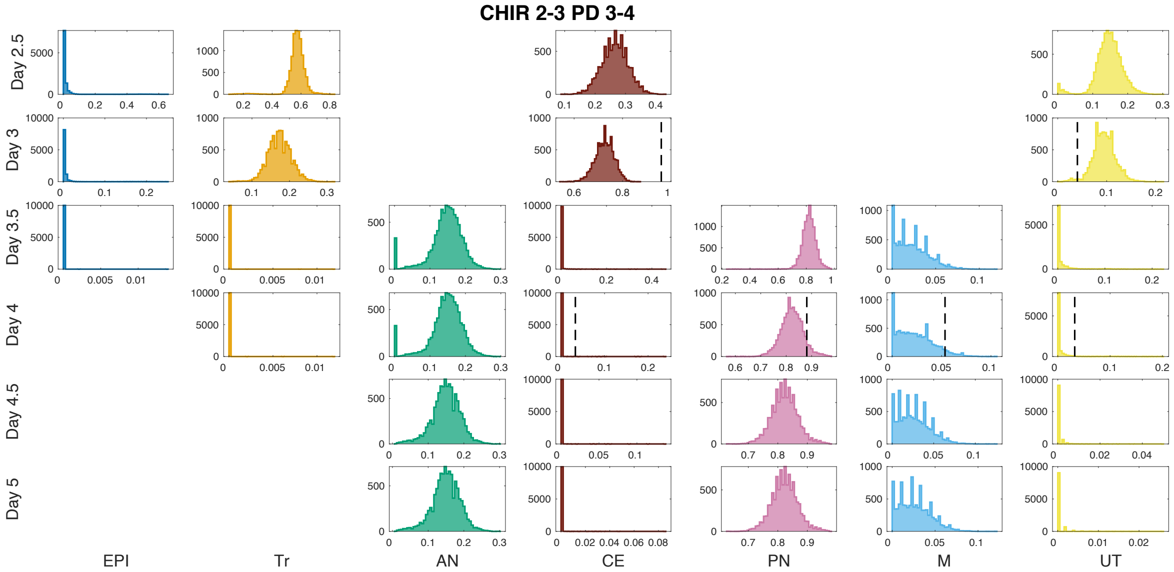

Figure SI65: Proportion distributions for CHIR 2-3 PD 3-4 regime simulations using the 10000 parameter vectors obtained in the last iteration of the refitting.

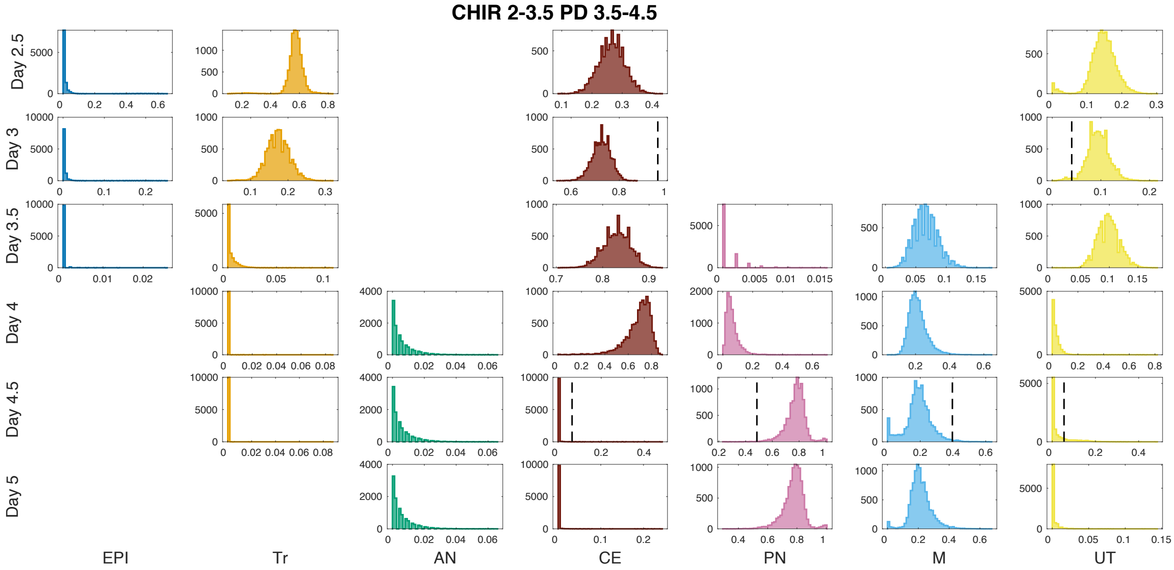

Figure SI66: Proportion distributions for CHIR 2-3.5 PD 3.5-4.5 regime simulations using the 10000 parameter vectors obtained in the last iteration of the refitting.

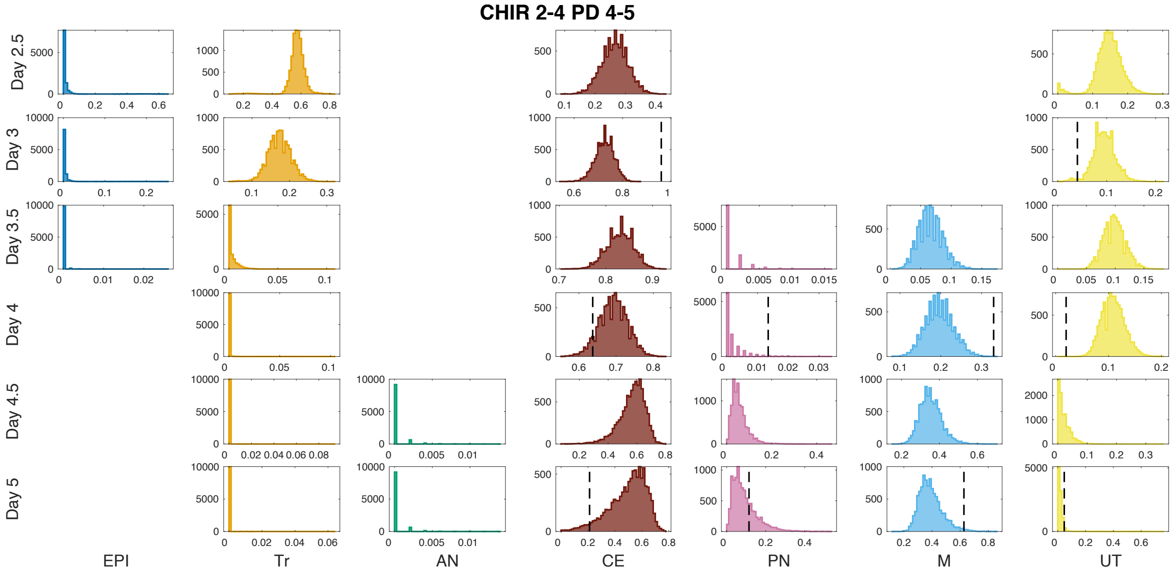

Figure SI67: Proportion distributions for CHIR 2-4 PD 4-5 regime simulations using the 10000 parameter vectors obtained in the last iteration of the refitting.

### References

[1] Robin Hartshorne. *Varieties.* Springer New York, New York, NY, 1977.

[2] Stephen Hawking. Stable and generic properties in general relativity. *Gen. Rel. Grav.*, 1:393–400, 1971.

[3] Stephen Smale. On gradient dynamical systems. *Annals of Mathematics*, pages 199–206, 1961.

[4] et al Rand, DA. Geometry and genetics. 2021.

[5] Christopher Zeeman and David Trotman. The classification of elementary catastrophes of codimension $\leq 5$. In *Structural stability, the theory of catastrophes, and applications in the sciences*, pages 263–327. Springer, 1976.

[6] Vladimir Igorevich Arnold, VS Afrajmovich, Yu S Il'yashenko, and LP Shil'nikov. *Dynamical systems V: bifurcation theory and catastrophe theory*, volume 5. Springer Science & Business Media, 2013.

[7] BA Khesin. Bifurcations in gradient dynamic systems. *Journal of Soviet Mathematics*, 52(4):3279–3305, 1990.

[8] G. J. McLachlan, S. K. Ng, and D. Peel. On clustering by mixture models. In Manfred Schwaiger and Otto Opitz, editors, *Exploratory Data Analysis in Empirical Research*, pages 141–148, Berlin, Heidelberg, 2003. Springer Berlin Heidelberg.

[9] E C Zeeman. The umbilic bracelet and the double cusp catastrophe. pages 1–39, 2013.

[10] Stefano M Iacus. *Simulation and inference for stochastic differential equations: with R examples.* Springer Science & Business Media, 2009.

[11] Desmond J Higham. An algorithmic introduction to numerical simulation of stochastic differential equations. *SIAM review*, 43(3):525–546, 2001.

[12] Peter E. Kloeden and Eckhard. Platen. *Numerical solution of stochastic differential equations.* Springer-Verlag Berlin ; New York, 1992.

[13] Elena Camacho-Aguilar, Aryeh Warmflash, and David A Rand. Quantifying cell transitions in c. elegans with data-fitted landscape models. *bioRxiv*, 2021.

[14] Jean-Michel Marin, Pierre Pudlo, Christian P Robert, and Robin J Ryder. Approximate Bayesian computational methods. *Statistics and Computing*, 22:1167–1180, 2012.

[15] Mark A Beaumont, Wenyang Zhang, and David J Balding. Approximate bayesian computation in population genetics. *Genetics*, 162(4):2025–2035, 2002.

[16] Mark M Tanaka, Andrew R Francis, Fabio Luciani, and SA Sisson. Using approximate bayesian computation to estimate tuberculosis transmission parameters from genotype data. *Genetics*, 173(3):1511–1520, 2006.

[17] Tina Toni and Michael P H Stumpf. Simulation-based model selection for dynamical systems in systems and population biology. *Bioinformatics*, 26(1):104–10, jan 2010.

[18] Juliane Liepe, Paul Kirk, Sarah Filippi, Tina Toni, Chris P Barnes, and Michael P H Stumpf. A framework for parameter estimation and model selection from experimental data in systems biology using approximate Bayesian computation. *Nature protocols*, 9, 2014.

[19] Tina Toni, David Welch, Natalja Strelkowa, Andreas Ipsen, and Michael PH Stumpf. Approximate bayesian computation scheme for parameter inference and model selection in dynamical systems. *Journal of the Royal Society Interface*, 6(31):187–202, 2009.

[20] Sarah Filippi, Chris P. Barnes, Julien Cornebise, and Michael P.H. Stumpf. On optimality of kernels for approximate Bayesian computation using sequential Monte Carlo. *Statistical Applications in Genetics and Molecular Biology*, 12(1), jan 2013.

[21] Mark A Beaumont, Jean-Marie Cornuet, Jean-Michel Marin, and Christian P Robert. Adaptive approximate Bayesian computation. *Biometrika*, pages 1–8, 2009.
